# Pan-tissue scaling of stiffness versus fibrillar collagen reflects contractility-driven strain that inhibits fibril degradation

**DOI:** 10.1101/2023.09.27.559759

**Authors:** K. Saini, S. Cho, M. Tewari, AA.R. Jalil, M. Wang, A.J. Kasznel, K. Yamamoto, D.M. Chenoweth, D.E. Discher

## Abstract

Polymer network properties such as stiffness often exhibit characteristic power laws in polymer density and other parameters. However, it remains unclear whether diverse animal tissues, composed of many distinct polymers, exhibit such scaling. Here, we examined many diverse tissues from adult mouse and embryonic chick to determine if stiffness (*E*_tissue_) follows a power law in relation to the most abundant animal protein, Collagen-I, even with molecular perturbations. We quantified fibrillar collagen in intact tissue by second harmonic generation (SHG) imaging and from tissue extracts by mass spectrometry (MS), and collagenase-mediated decreases were also tracked. Pan-tissue power laws for tissue stiffness versus Collagen-I levels measured by SHG or MS exhibit sub-linear scaling that aligns with results from cellularized gels of Collagen-I but not acellular gels. Inhibition of cellular myosin-II based contraction fits the scaling, and combination with inhibitors of matrix metalloproteinases (MMPs) show collagenase activity is strain - not stress- suppressed in tissues, consistent with past studies of gels and fibrils. Beating embryonic hearts and tendons, which differ in both collagen levels and stiffness by >1000-fold, similarly suppressed collagenases at physiological strains of ∼5%, with fiber-orientation regulating degradation. Scaling of *E*_tissue_ based on ‘use-it-or-lose-it’ kinetics provides insight into scaling of organ size, microgravity effects, and regeneration processes while suggesting contractility-driven therapeutics.

## Introduction

Physical properties of polymer systems such as stiffness often scale with concentration [1], which raises questions about the most abundant polymer Collagen-I within animals. Such fibrillar collagens confer stiffness to tissues as demonstrated by rapid fluidization occurring in ∼1 hour or less upon collagenase to an intact solid tissue. The variation reflects tissue differences: rigid bone has much more Collagen-I than soft brain, for example [2]; and such systematic differences might reflect scaling relationships that apply across tissue types, species, and development. Robust scaling can help clarify the accuracy of measurements as well as the effects of tissue treatments (e.g. drugs) and diseases such as fibrosis and cancer [3]. Pan-tissue studies with methods such as single-cell RNA-sequencing enable novel comparisons [4], but for the most abundant animal protein, trends and underlying mechanisms remain obscure.

Strong scaling of stiffness with purified Collagen-I as in vitro gels proves reproducible [5] [6] [7]:

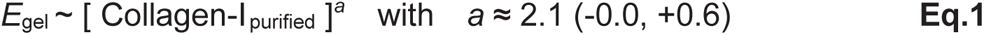

Although this power law conforms to theory of semi-flexible polymers [5] [7], our recent study of nine diverse tissues from adult mouse that ranged from brain to bone [2] plus a second study of several developing organs [8] using Mass Spectrometry (MS) proteomics of extracted proteins proposed a much weaker power law over multiple logs of tissue data:

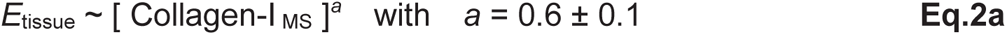

Each tissue type was assumed to have a characteristic stiffness, which continues to be debated [9]. Also, collagen-associated enzymes such as lysyl oxidases (LOX, LOXLs) that crosslink fibrillar collagen and increase *E*_tissue_ can add complexity – even though they scale with Collagen-I [8] [10] - so that Collagen-I scaling implicitly captures many fibril-related factors. Nonetheless, tissue extraction limitations include incomplete yields (e.g., partial tissue digestion) plus complications from different crosslinks and levels in tissues. Moreover, stiffness measurement accuracy can be limited by insensitivity of the technique to collagen fibers due to probed length scales and/or time scales [11] [12] as well as contributions from heterogeneous collagen distribution within tissues [13]. To address these limitations, we hereby study intact tissues by label-free second harmonic generation (SHG) imaging – including response kinetics to molecular perturbations and tissue forces that are cell-generated or external and sometimes paired with stiffness measurements.

SHG imaging is now widely applied, even in the clinic [14], for label-free observations of biointerfaces and particularly for collagen fibrils [15]. Among fibrillar collagens, Collagen-I produces much stronger SHG signal than other types including Collagen-2, 3, 5 [16]. For gels of Collagen-I fibrils, SHG signal increases with collagen concentration [6]. Sensitivity of SHG to local orientation of interfaces [15] and sample thickness scattering effects can in principle be minimized by low magnification imaging with backward signal collection, but this needs to be established for a given microscope and set of samples. Rapid collagenase digestion should also help clarify collagen fibril contributions to SHG signal – which we quantify here across diverse embryonic and adult tissues.

Collagen-I is lacking in early embryos, which are very soft, but collagen accumulation in embryonic heart becomes directly measurable and also implicated by tissue stiffening which helps to pressurize and pump blood through the expanding vasculature [17]. Even higher mechanical stress is generated or sustained by adult muscles, tendons, cartilage and bones, consistent with high collagen in these stiff tissues relative to soft tissues such as brain or liver (**Eq.2**). Whether stress or strain (≈ stress / *E*) in a tissue relates to such different collagen levels remains unclear; a relation is suggested by inhibition of cardiomyocyte contractile forces in diseased hearts that led to decreased fibrotic collagen even though cardiomyocytes do not express collagen [18]. The myosin inhibitor drug Mavacamten is now clinically important [19]. Furthermore, skin wound softening via myosin-II inhibition (using Blebbistatin) in pre-clinical mouse models shows improved healing [20]. More broadly, proposed “use-it-or-lose-it” mechanisms for muscle [21], bone [22], heart [23] and brain [24] require a deeper insight into any mechanically-driven intrinsic remodeling of collagen in extracellular matrix (ECM)– with potential applications to gravity effects relevant to spaceflight [25] [26] and to scaling Organ-size ∼ (Body-weight)^*β*^ scaling across diverse tissues [27] [28] [29] [30].

In highly reductionist studies using purified Collagen-I fibrils [31] and trimers [32], applied force respectively suppresses and accelerates collagen degradation rate. With “cell-free” tissue or constructs treated with bacterial collagenase (BC), externally applied force is thought to modulate degradation as inferred from mechanical measurements [33] [34] [35] [36] [37]. Direct visualization of collagen degradation has often been lacking, especially in animal tissues with endogenous collagen-degrading matrix metalloproteinases (MMPs) that target distinct sites on collagen compared to exogenous BC [38, 39]. Within intact animal tissues, whether forces sculpt both embryonic collagen and tissue mechanics via MMP activity is unclear as is the regulation of degradation by local stress versus local strain in relation or not to stiffness.

Our main result here shows that SHG signal during collagenase-mediated degradation parallels tissue softening and conforms to a pan-tissue stiffness-vs-SHG power law – as summarized below with other relevant scaling results (**Table 1**). Mechanistically, we show contractile forces in tissue normally suppress MMP-collagenase activity, with inhibitor results and also enzymatic crosslinking again fitting the stiffness-vs-SHG power law. Furthermore, the ∼5% beating strains in extremely soft embryonic heart that suppress collagen degradation by endogenous MMPs are the same local strain levels in stiff decellularized tendon that maximally suppress collagenase. Our studies also make use of new modes of tissue deformation and novel peptides for visualization, utimately supporting a “use-it-or-lose-it” basis of pan-tissue collagen scaling.

**Table 1:**
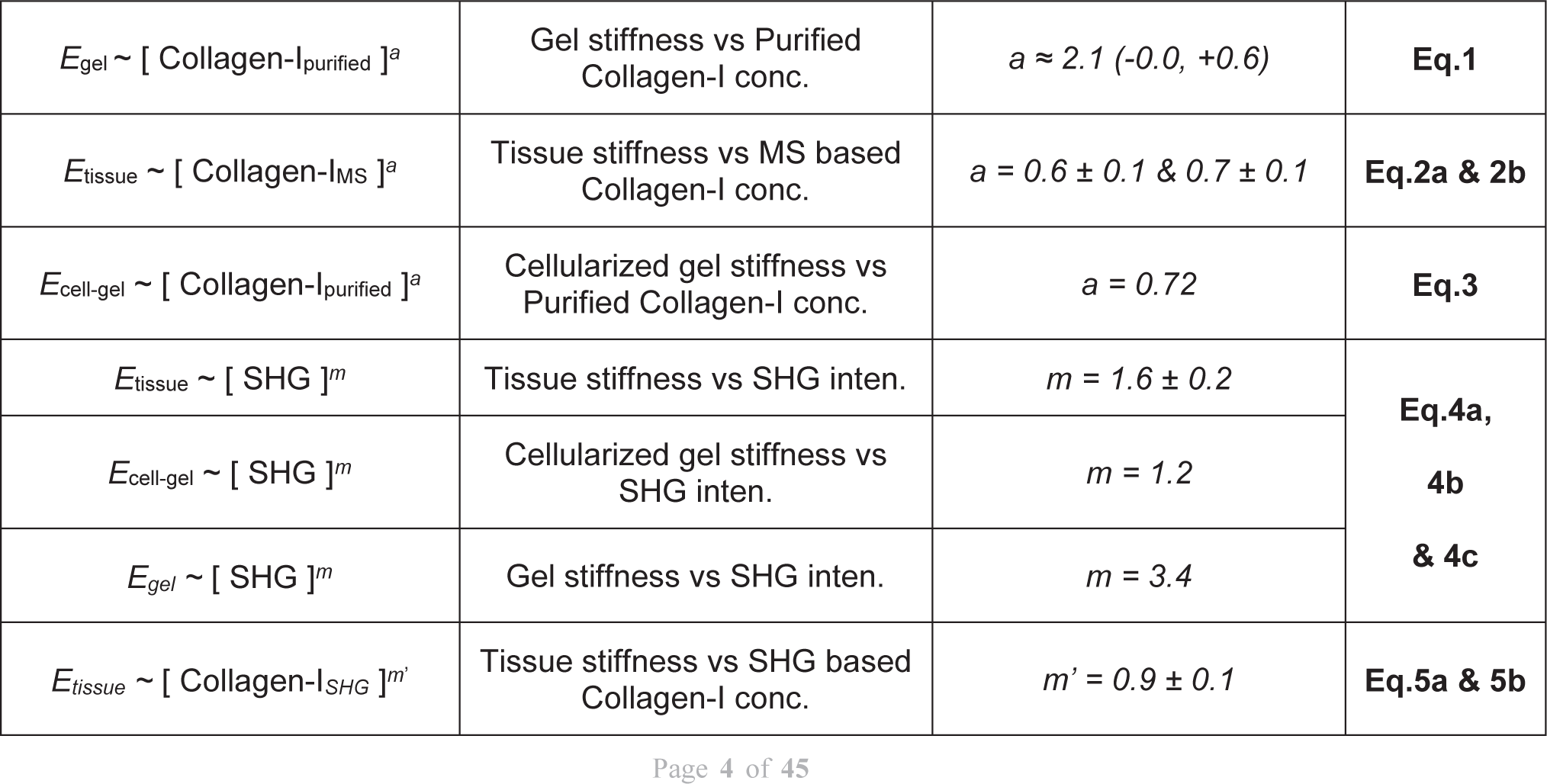
Scaling equations and exponents.

## Results

### Collagenase kinetics of tissue SHG and stiffness in pan-tissue scaling

To first clarify the dependence of the tissue SHG signal and stiffness on fibrillar collagen, we measured the kinetics of collagenase (BC) effects on adult mouse tissues. For stiffness: aspiration into a 50-100 μm diameter micropipette is seen to deform tissue cells and ECM, and collagenase softens adult heart within ∼60 min which tracked well with decreasing SHG signal (**Fig.1A,B**). Post-aspiration: untreated heart recovered completely as an elastic tissue, but collagenase caused tissue fluidization and an inelastic response. Neither SHG signal nor stiffness changed with collagenase-treated soft brain (*E*_tissue_ ∼ 0.4 kPa) having very low collagen levels. Slightly stiffer day-4 chick embryo hearts (E4) (*E*_tissue_ ∼ 1 kPa) show decreased SHG signal and stiffness with collagenase **(Fig.1C-i)**.

**Fig.1.**
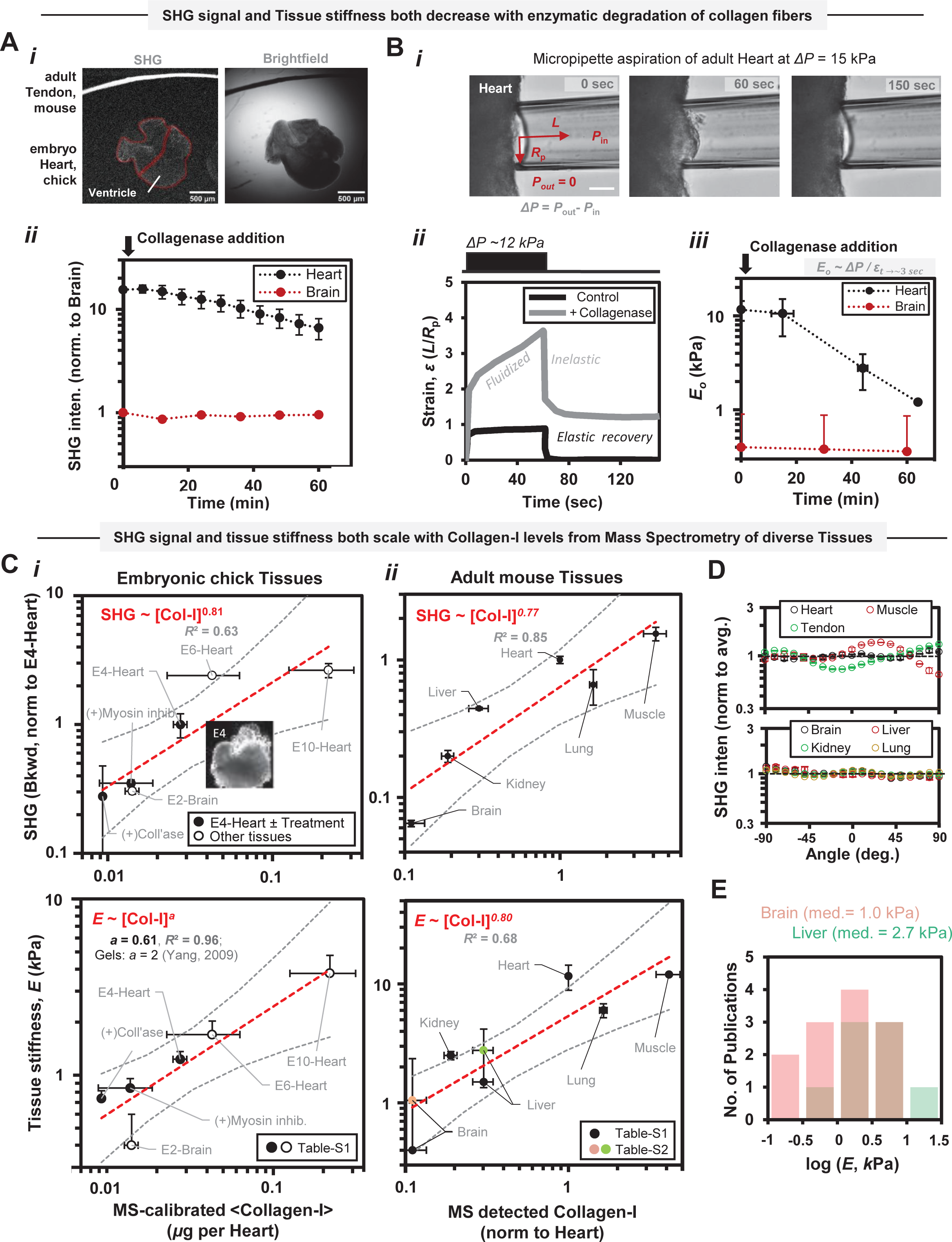
Label-free measurements reveal both Second harmonic generation (SHG) signal and tissue stiffness scale with collagen-I protein levels in embryonic and adult tissues. **A.** Rapid loss of SHG signal in tissue following exposure to exogenous collagenase (i.e., bacterial collagenase (BC)) demonstrates SHG signal as a label-free surrogate for fibrillar collagen levels. (i) SHG signal intensity difference between isolated embryonic day-4 (E4) chick heart and adult mouse tendon shows their highly different fibrillar collagen levels. (ii) SHG signal loss in adult mouse heart upon collagenase treatment confirms fibrillar collagen dependence of SHG signal. Collagenase treatment leads to degradation of abundant fibrillar collagen in the heart tissue while low fibrillar collagen levels and/or low collagenase permeation leads to negligible SHG signal loss in brain tissue. Moreover, SHG signal scales with purified collagen-I concentration (Fig.S1-i). n ≥ 3 samples; Error bars indicate ±SEM. **B.** BC treatment rapidly decreases stiffness (*E* or elastic modulus) of tissues. (i-ii) Measurement of tissue stiffness by micropipette aspiration, based on applied suction pressure (*Δ*P) and instantaneous strain (*ε*) values, in case of adult mouse heart tissue exposed to collagenase. (iii) Collagenase treatment rapidly decreases stiffness of adult mouse heart but not of brain tissues. n ≥ 3 samples; Error bars indicate ±SEM. **C.** Both SHG signal and tissue stiffness increase with collagen-I levels in tissues following a power-law scaling. (i-ii) Top: SHG signal scaling with collagen-I levels, measured by label-free mass-spectrometry (MS), is similar between embryonic chick tissues and adult mice tissues (SHG ∼ [Collagen-1]^a^, *a* = ∼0.8). Bottom: Tissue stiffness scales with collagen-I levels (*E* ∼ [Collagen-1]^a^) in embryonic chick tissues (*a* = 0.6) and adult mice tissues (*a* = 0.8) (Table-S1) and, previously reported stiffness values for brain, liver tissues from diverse species (Table-S2) obey such scaling trends. The dissimilar scaling exponent value a likely results from partial digestion of tissue for MS measurements due to higher collagen crosslink levels in adult tissues than embryonic tissues. **D.** SHG intensity variation with fiber-angle indicates more aligned collagen in stiff tissues (heart, muscle and tendon) than soft tissues (brain, liver, kidney and lung) but such variations might not affect the observed scaling exponent due to log-log nature, random selection of tissue locations for measurements. SHG signal value at each angle (normalized to average value across all angles (−90 to +90 degrees)) is further averaged for backward and forward directions. n ≥ 3; Error bars indicate ±SEM. **E.** Previously reported stiffness values for same tissue isolated from diverse species have same order of magnitude for brain and liver tissues (Table-S2) with observed variance likely resulting from the difference in probed length-scale, deformation rate, etc.

To try to assess inter-relationships among intact tissue SHG signals, *E*_tissue_, and Collagen-I levels, we quantified SHG signal from adult mouse and embryonic chick tissues for which we have MS measurements of extracted Collagen-I levels **(Fig.1C-i,ii top)**. The embryonic data includes tissues from different days of development (E4-E10) as well as perturbations including collagenase. Importantly, both plots for tissue SHG versus MS for Collagen-I (i.e., SHG ∼ [Collagen-I _MS_]^a^) show power law exponents of *a* ≈ 0.8 (**Table 1**). The striated cytoskeletal interfaces of muscle and heart [15, 40] contribute weak SHG signals, and sample orientation effects are also weak, with significance only for tendon and skeletal muscle as expected for tissues with some aligned collagen (**Fig.1D**).

The fractional power law SHG ∼ [Collagen-I_MS_]^0.8^ is close to linear, and reconstituted gel studies with known concentrations of Collagen-I fit with *a* = 1 even though *a* = 0.62 is a better fit (**Fig.S1-i** shows fit of Fig.3A in [6]). As a model for insight into such differences, if fibril radius increases with concentration as *R* ∼ [Collagen-I]*^a^*, then interfacial signal scales with *R* per fiber length as SHG ∼ [Collagen-I]*^a^*, and Bulk signal (such as from MS) scales with *R*^2^ per fiber length as Bulk ∼ [Collagen-I]^2*a*^, which gives an SHG versus Bulk power law with *a* = ½ – except for the case of constant *R* case (*a* = 0). The latter corresponds to *a* = 1 and is closer to our tissue result of *a* = 0.8. This 20% difference could be explained by underestimation of high Collagen-I in stiff tissues by SHG per fiber mechanisms above or else that Collagen-I extractions for MS measurements are less efficient for softer tissues.

Plotting *E*_tissue_ (**Table-S1**) versus MS results **(Fig.1C-i,ii bottom)** incorporates some variation in published *E*_tissue_ (e.g. brain and liver: **Fig.1E**) with median values that respect but de-weight extreme values. The differences for *E*_tissue_ among publications likely reflect differences in species and probed length- or time-scales among other reasons (**Table-S2**), but median data for brain and liver perfectly fit the scaling trend for adult tissues. Moreover, the power law based on MS measuremets of developing (chick) tissues plus perturbations and separately for adult (mouse) tissues including updated stiffness measurements give

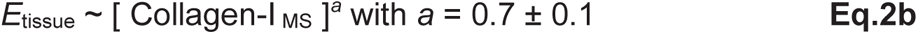

which robustly aligns with the previously cited MS based scaling law for adult tissue (**Eq.2a**). The MS further showed weak power laws for less abundant fibrillar collagens (Col-3, 5, 6, 11, 12 gave *a* = 0.7 to 1.2) that seems consistent with an independent measurement of hydroxyproline in extracted tissue collagen yielding an aggregate power law of *a* = 1.1 (**Fig.S2A**).

Based on MS and required extractions, stiffness scaling is far weaker for tissue than for acelluar Collagen-I gels (**Eq.1**, **Fig.S1-ii**). However, fibroblasts in Collagen-I gels are well known to contract and remodel the gels with the key cited study [6] showing overall softening for a given Collagen-I level compared to an acellular gel and also a scaling that matches our tissue scaling (**Eq.2a,b**):

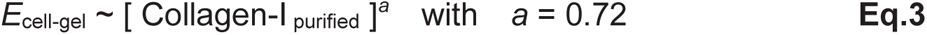

Protein extraction concerns for MS nonetheless motivate further analyses using SHG signal from intact tissues.

### Universal scaling of tissue stiffness with SHG signal fits diverse molecular perturbations

All tissue SHG measurements above were combined into a single plot with corresponding data for stiffness **(Fig.1C-i,ii)** with results also added for extremely stiff adult tendon plus several collagenase-degraded or enzymatically-crosslinked tissues. A scaling relation over many logs (**Fig.2A**, *R*^2^ > 0.95) shows that stiffness and SHG signal vanish together, which is reasonable given the fact that embryos are extremely soft and lack SHG-active collagen fibers [17]. The tissue power law exponent *m* is once again much weaker than for pure gels and only slightly stronger than for a cell-in-gel system:

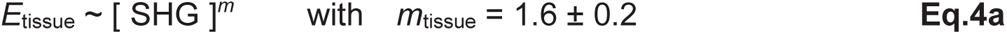

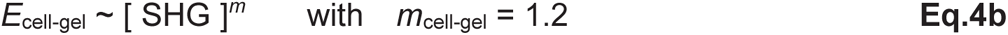

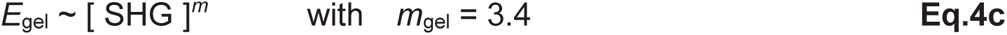

**Fig.2:**
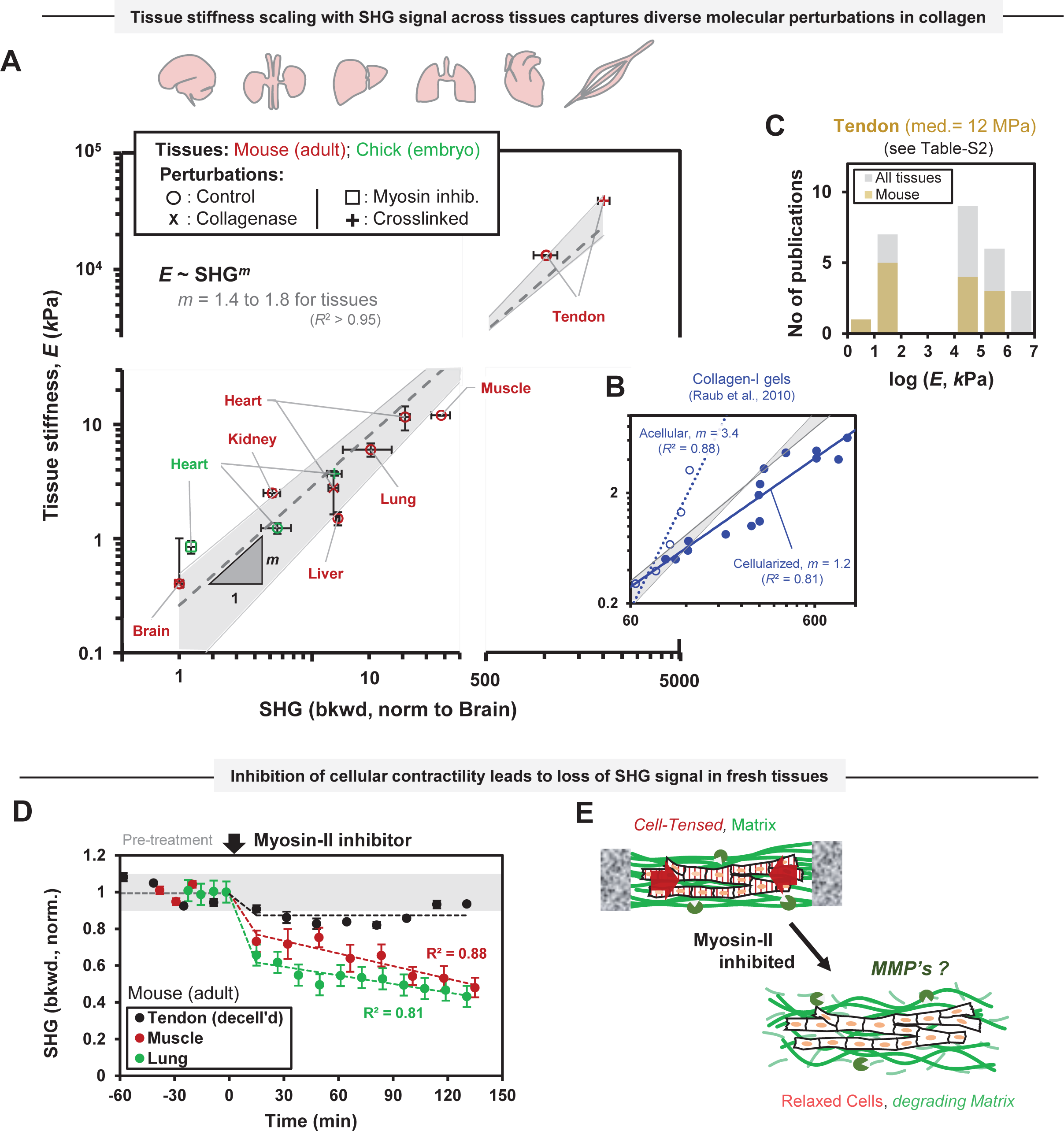
Power law scaling of tissue stiffness with label-free second harmonic generation (SHG) signal produced by different tissues both suggests mechanical strain-suppressed fibrillar collagen degradation in live tissues underlie the scaling relationship and captures diverse molecular perturbations. **A.** E4-chick hearts (green) and tail tendon as well as other tissues (i.e., brain, kidney, liver, lung, heart and muscle (skeletal)) from adult mouse (red) show power-law scaling of tissue stiffness with backward SHG signal (*m* = 1.4 to 1.8). The loss of both stiffness and SHG signal values upon myosin inhibition in E4-chick hearts suggests a loss of fibrillar collagen level due to degradation by endogenous matrix metalloproteinases (MMPs). Both stiffness and SHG signal values decrease in adult mouse heart upon collagenase treatment (per Fig.1A-B) but increase upon collagen crosslinking (with exogenous transglutaminase (TGM)) in E4-hearts and adult mice tendons. For mice tendon stiffness value, see Table-S2 and Fig.2C. Symbols: “circle” = control; “cross” = Collagenase treatment by BC; “square”: Myosin-II inhibition using Blebbistatin; “plus” = Crosslinking by TGM. n ≥ 3 samples per condition; Error bars indicate ±SEM. **B.** The exponent for tissue stiffness versus SHG signal scaling is similar to cellularized purified collagen-I gels (*m* = 1.2) but very different from acellular gels (*m* = 3.4) (Fig.S1-ii). **C.** Previously reported stiffness values for tendon tissues (isolated from diverse species and from different anatomical locations) differ by several orders of magnitude and with observed variance likely resulting from alignment of fibrillar collagen and probed length-scale for local stiffness measurement besides compositional differences. **D.** Inhibition of cellular contractility via addition of myosin-II inhibitor (i.e., Blebbistatin) leads to SHG signal loss in ex-vivo adult mouse tissues namely skeletal muscle and lung. In contrast, SHG signal produced by decellularized tendon tissues is relatively unaltered during myosin-II inhibitor treatment. **E.** Myosin-II based cellular contraction produces a tensed tissue extracellular matrix (ECM) that is difficult to be degraded hypothesized endogenous MMPs, whereas mechanically relaxed ECM of myosin-inhibited cells in a tissue make fibrillar collagen highly susceptible to degradation by MMPs. Thus, mechanical state of a tissue ECM controls its ECM fibrillar collagen levels.

The latter two SHG power laws (per **Fig.2B**) derive from the same SHG-focused study that used known levels of purified Collagen-I [6], and of course conversion of SHG signal scaling to Collagen-I scaling is crucial for insight into the biology. Importantly, the power laws for stiffness versus Collagen-I are both ∼40% smaller than stiffness vs SHG signal : *m* = 3.4 (**Eq.4c**) versus *a* ≈ 2.1 (**Eq.1**), and *m* = 1.2 (**Eq.4b**) versus *a* ≈ 0.7 (**Eq.3**). Thus, SHG underestimates high levels of Collagen-I, with possible mechanisms including increased fibril diameter at high Collagen-I. Our tissue studies with MS likewise showed SHG underestimates high levels of Collagen-I (i.e. 0.8 power law in **Fig.2B**). Hence, for a best estimate of Collagen-I from SHG signal for intact tissue, we apply the 40% correction above to our SHG signal result (*m*_tissue_ = 1.6 ± 0.2 per **Eq.4a**) in order to arrive at a power law *m*’ in terms of Collagen-I in intact tissues:

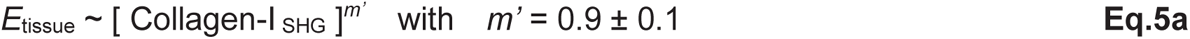

The same scaling is obtained by simply combining the MS-based scaling of both tissue SHG and *E*_tissue_ with Collagen-I (**Fig.1C**):

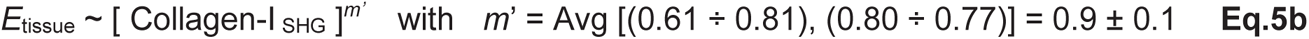

Both results compare well to stiffness scaling with [ Collagen-I _MS_]*^a^* (*a* = 0.7 ± 0.1 in **Eq.2b**). Thus, tissue stiffness scaling is a sub-linear function of Collagen-I levels measured by either SHG or MS.

The scaling result for SHG (**Fig.2A**, **Eq.4a**) should be reproducible across labs and can help sort out tissue stiffness measurements based on the idea that extra scrutiny is required of any outlying data. For tendon results (**Fig.2A**), mouse tail tendon has been measured to have high stiffness (∼10 MPa [41]) in the axial direction related to spindle-shaped tenocyte orientation, but the stiffness value in the plot instead represents the median of a large variance in mouse tendon measurements (**Fig.2C**). Differences are attributable in part to probed lengthscales, directionality of stiffness measurement, and perhaps species (**Table-S2**). Nonetheless, the minimal angular dependence of SHG signal (**Fig.1D**) is unlikely to affect the robust scaling trend due to the insensitivity of log scale data and random selection of tissue location for measurements.

Our key scaling result (**Fig.2A**) shows robustness when challenged by molecular perturbations not only to fibrillar collagen via collagenase (per adult mouse hearts **Fig.1A,B**) and tissue-generated cell forces but also to Transglutaminase (TGM) crosslinking. The latter increases *E_tissue_* in E4-chick hearts [17] and rat tendon fibers [10] by 3-fold but also – surprisingly – increases SHG signal by 2-fold (**Fig.2A,S23**), and we selected TGM as it is regulated independent of tissue Collagen levels whereas many LOX enzymes increase with Collagen-I as previously noted. The E4-chick hearts beat autonomously [17], but inhibiting myosin-II contractility (with blebbistatin) rapidly decreased both SHG signal and *E*_tissue_ (in ∼1 hr). SHG signal is similarly decreased in ex vivo lung and skeletal muscle upon myosin-II inhibition unlike the decellularized tendon (**Fig.2D**), which underscores the mechano-regulation of fibrillar collagen levels by cells in embryonic and adult tissues. These latter studies led us to hypothesize that relaxed tissue allows for fibrillar collagen degradation via endogenous MMPs (**Fig.2E**). Such a mechanism could explain the scaling of normal and perturbed tissue.

### Cellular contractility and external tissue forces suppress collagenases in tissues

To understand which cells generate contractile forces on ECM of beating embryonic heart and which make collagen and MMPs, single-cell RNA sequencing of E4-chick hearts was applied. This revealed fibroblastic cell-types express both collagen-I and MMPs whereas cardiomyocytes expressing myosin-II drive contractile beating but do not synthesize collagen-I heterotrimers (**Fig.3A,S3E**). Note that MMPs target a specific-site on fibrillar collagens (producing ¾ and ¼ fragments) [38] in contrast to BC targeting multiple sites. The observed ∼5% peak strain in E4-chick hearts (beating at ∼1 Hz) is suppressed within minutes upon myosin-II inhibition via Blebbistatin (**Fig.3B-i**), and SHG signal rapidly begins to decrease (by ∼50%) within ∼1-2 hours (**Fig.3B-ii,left**) but such SHG signal loss is rescued in the presence of a pan-MMP inhibitor (GM6001). Exogenous collagenase treatment (via BC) also inhibits the beating and has a similar effect on SHG signal (**Fig.3B-ii,right**), although beating-suppression kinetics seems faster perhaps because the heart stops beeating and thereby unleashes endogenous MMPs to contribute towards ECM degradation. Unperturbed E4-chick hearts show constant SHG signal over several hours (**Fig.S3**) despite considering small changes in collagen-I level due to heart development (∼1% per hour from E4- to E6-day) (**Fig.1C**). Importantly, myosin-II inhibition in E4-chick hearts suggests that eliminating the ∼5% strain or else the stress associated with beating triggers fibrillar collagen degradation by endogenous MMPs.

**Fig.3.**
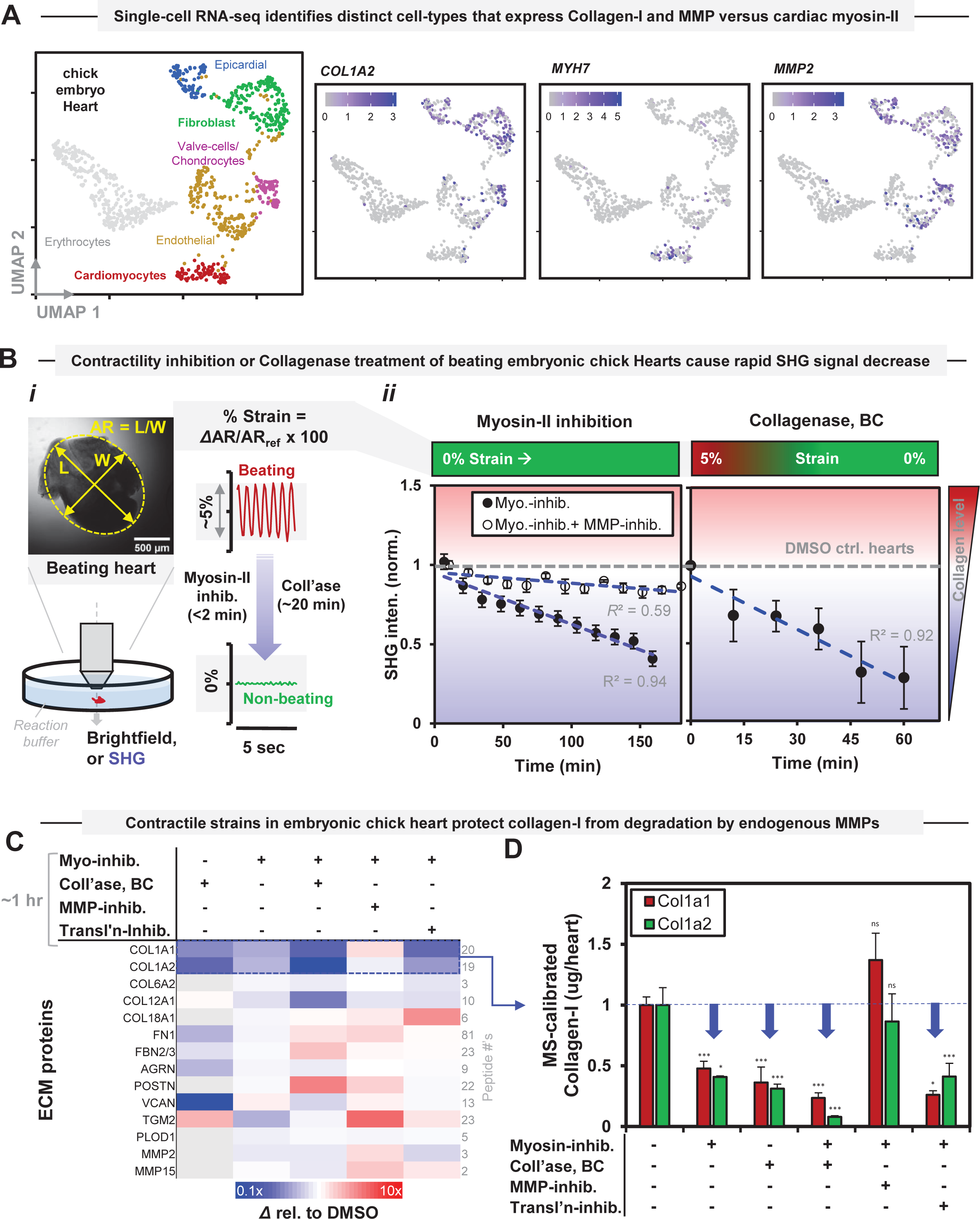
Cardiomyocytes contraction maintains fibrillar collagen level in live hearts via suppression of enzymatic degradation by endogenous MMPs. **A.** Single-cell RNA-sequencing identifies distinct cell-types within E4 chick hearts. Fibroblastic cell-types express collagen-I (*COL1A2*) and abundant MMPs (*MMP2*) while cardiomyocytes express myosin-II (to drive contractile heart beating) not collagen-I. Color bar(s) indicates mRNA expression level (*loo*_c_ normalized). **B.** Beating induced contractile strains maintain fibrillar collagen levels E4 chick hearts while beating inhibition results in collagen level loss. (i) Strains in hearts (beating at ∼1 Hz and measured using temporal aspect-ratio (AR) change) immediately decrease from ∼5% to 0% following myosin-II inhibition (using Blebbistatin). Realtime changes in fibrillar collagen levels in hearts are assessed using forward SHG signal. (ii) Myosin-II inhibition results a rapid loss of ∼50% fibrillar collagen level within 3 hours similar but inhibition of endogenous MMP activity (using GM6001) rescues such SHG signal loss. The loss in SHG signal upon exogenous collagenase (BC) treatment confirms fibrillar collagen basis of SHG signal in live E4-chick heart. Fibrillar collagen levels in DMSO control hearts remain unaltered with time (Fig.S3). n = 4 hearts per condition, averaging ≥ 2 regions per heart; Error bars indicate ±SEM. **C-D.** MS revealed many ECM proteins and showed endogenous MMPs degrade fibrillar collagens in E4-chick hearts. Protein level changes (fold-change relative to DMSO control) following perturbations i.e., myosin-II inhibition with/without BC treatment, inhibition of endogenous MMP activity or protein translation (using Cycloheximide) are shown as heat-map. Contractility-inhibited hearts showed a decay of MS signal for many ECM proteins. Particularly, collagen-I (subunits COL1A1 and COL1A2) levels decreased in the presence of exogenous BC or endogenous MMPs regardless of protein synthesis while the presence of pan-MMP inhibitor rescued the loss of collagen-I as well as many ECM proteins. n = 8 hearts per condition; Error bars indicate ±SEM; * Two-tailed t-test: * = p < 0.05, ** = p < 0.01, *** = p < 0.001.

To directly quantify collagen-I and any strain- or stress-dependent changes in perturbed E4-chick hearts and endogenous enzymes, the protein content was measured using MS proteomics. Calibrations with collagen-I proved linear (**Fig.S3**), and MS also identified endogenous MMP-2 [42] and MMP-15 (**Fig.3C**). Importantly, myosin-II inhibited hearts showed rapid decreases in COL1A1 and COL1A2 by >50% (∼1 h) (**Fig.3C,D**), which is similar to SHG signal loss. Such decreases fit the power-law for tissue stiffness scaling versus SHG signal (**Fig.2A**) and collagen-I (**Fig.1C**). Moreover, a pan-MMP inhibitor rescued collagen-I degradation in the contractility-inhibited hearts (similar to that of SHG signal loss, **Fig.3B-ii**), demonstrating endogenous MMP activity for degradation. Because MMP-2 level decreases in contractility-inhibited hearts and because both MMPs increase with the pan-MMP inhibitor (**Fig.3C**), Collagen-I degradation is not explained by induced changes in MMPs levels; it is also likely that degradation of pro-MMP upon pan-MMP inhibition [43] promotes pro-MMP synthesis. Exogenous BC shows the expected decrease in COL1 levels. Transglutaminase-2 levels (TGM2, with 23 peptides) mediate collagen crosslinking and correlate with MMP levels in both contractility-inhibited and pan-MMP inhibitor treated E4-hearts, showing sensitvity to acute ECM softening with BC treatment. PLOD1 leads to synthesis of lysyl hydroxylase-1 producing collagen crosslinks and does unaffected by any perturbation, wereas Fibronectin increases upon contractility inhibition (81 FN peptides), suggesting E4-hearts either synthesize some proteins and/or their degradation is decreased. Given the possible effects of protein synthesis, we combined mRNA translation inhibition with contractility-inhibition and again found decreased collagen-I (**Fig.3C,D**), and electrophoresis analyses confirm the various perturbations to collagen levels (**Fig.S3**).

To address whether strain or stress in tissue controls collagen degradation rate and to try to generalize tissue ECM interactions with different exogenous collagenases including purified MMP-1 [38], we deformed tendons with their longitudinally aligned collagen [44] and high stiffness relative to E4-chick hearts. This was done in two different modes while quantifying collagenase-driven degradation (**Fig.4**), after making them “cell-free” via freeze-thaw and cold-storage (**Fig.S4**). The mechano-bioreactor imposes heterogeneous strains (via three-point bending) with real-time imaging by fluorescence and SHG (**Fig.4A,B,S5,S6;supp.-methods S1,S2**). It helps overcome some conventional challenges of sample holding and accuracy of local tissue strain measurements by using fluorescence patterned photobleaching (FPP) on tendons labelled with a novel Aza-peptide or widely used Col-F (**Fig.S7,S8;supp.-methods S3**). Aza-peptides (-[azGPO]_3_) exhibit collagen triple-helical propensity similar to Glycine-Proline-Hydroxyproline (GPO) repeat peptides [45] and are expected to possess increased stability against enzymatic degradation [46] and, also – importantly – do not influence tendon degradation by collagenases (**Fig.S9**).

**Fig.4.**
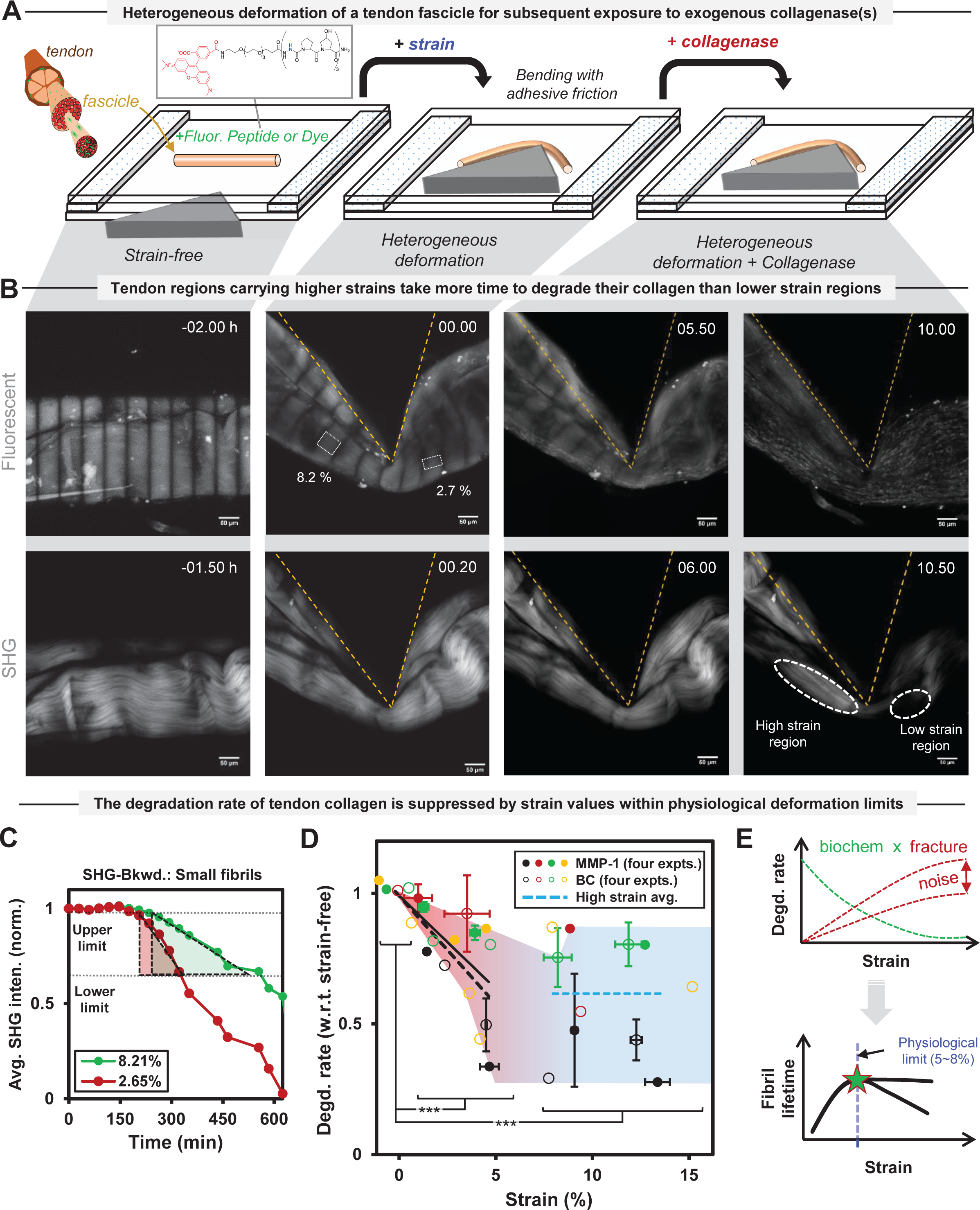
Physiological strains suppress fibrillar collagen degradation in devitalized tendon by exogenous collagenases including purified MMP1. **A.** Three-point bending induced heterogeneous deformation and degradation of “cell-free” mouse tendon fascicles fluorescently labelled with either a novel Aza-peptide or Col-F dye for strain quantification using fluorescent pattern photobleach (FPP). **B.** Undeformed and heterogeneously deformed configurations of a representative tendon fascicle at different time points (L to R). Fluorescent (after pattern photobleach; top-panel) and SHG (bottom-panel) images represent strain distribution and fibrillar collagen content respectively. Native collagen fiber crimps in the undeformed configuration align up along the fascicle-axis following mechanical deformation of the tendon. **C.** For each region of the deformed fascicle, the initial slope of normalized SHG intensity in backward direction (SHG-Bkwd.) versus time represented the degradation rate of fibrillar collagen. **D.** Strain magnitudes of up to ∼5–8% in the fascicles exposed to either matrix metalloproteinase-1 (MMP-1; solid-line) or bacterial collagenase (BC; dashed-line) similarly suppress its collagen degradation rate (*p* = 0.00056 for 2– 8% strain) relative strain-free configuration (i.e., 0±1% strain) regardless of the difference in cleavage activity of MMP-1 and BC on monomeric collagen, whereas the degradation rates are strain-independent at larger strain values (i.e., >8%; dashed blue line, *p* = 0.00016). Fluorescent labelling via either Aza-peptide or Col-F dye does not influence the degradation rate (Fig.S9). n = 4 fascicles per collagenase. Error bars indicate ±SEM; *Two-tailed t-test: * = p < 0.05, ** = p < 0.01, *** = p < 0.001. **E.** Pan-strain dependent degradation is resultant of biochemical regulation and tissue fracture (top). Physiological tissue strains (∼5-8%) maximize collagen fibril lifetime while pathological strains (>8%) cause tissue fracture (bottom).

Local strain quantification by FPP (**Fig.4B-upper**) and tissue collagen content assessment via SHG signal (**Fig.4B-lower,4D;supp.-methods S4**) showed that when deformed tendon is exposed to exogenous collagenases (purified MMP-1 [38] or BC), the initial rate of collagen degradation is suppressed by strain. Such microscale measurements (made on ∼50 *µ*m x 50 *µ*m region of interest (ROI) (**Fig.S9**)) are sufficient to assess collagen fibers within undeformed tendons that exhibit native crimps consistent with low residual strain [47, 48]. Importantly, total times required for either MMP-1 or BC at similar concentrations (i.e., ∼100 nM and assuming all enzyme is active) to degrade fibrillar collagen in tendon tissues were similar (∼hours), which is surprising because BC cleaves monomeric collagen-I >10-fold faster than MMP-1 (**Fig.S10;supp.-methods S5**) [49]. Regardless, widely used trypsin exposure (5 mg/mL for ∼48 hours) did not produce any observable change in tendon tissues (comparable to collagenase treatment) likely because trypsin targets more denatured (or unfolded) collagen triple helices than natively folded collagen [50]. These observations demonstrate that strained tissue rather than the enzyme regulates the degradation reaction. Degradation rate versus strain for both MMP-1 and BC (**Fig.S11**) indeed showed similar degradation rate suppression for up to ∼5% strain values (i.e., physiological strain limit for tendon tissues) whereas degradation rates became strain-independent at pathological strains (i.e., >8% strain) (**Fig.4D**).

Hence, SHG signal and MS measurements of different tissues together show that physiological ∼5% tissue strain (measured at microscale or higher) are sufficient to impede degradation of nanoscale diameter collagen fibrils sustaining lower strain than microscale tissue [47]. Given that stress and strain in a material are constitutively related, simply multiplying tissue strain by *E*_tissss_ gives stress magnitude sustained by a tissue. For E4-chick heart versus adult mouse tendon, ∼5% tissue strain results in similar SHG signal changes in both tissues whereas tissue stress differs by almost 10^4^-fold. Therefore, tissue strain (rather than stress) better explains collagen degradation suppression in tissues.

### Strain-dependent collagen degradation rate relates to local orientation

To identify what strain-driven structural change(s) are associated with degradation of collagen fibers, tendon tissues under physiological uniaxial extension subsequently exposed to BC were imaged (**Fig.5A-i**). Overall, the strain-suppressed collagen degradation of unixially deformed tendons based on SHG signal (**Fig.5A-ii**) overlapped with the measurements from the three-point bending approach (**Fig.4D**). Surprisingly, the FPP-measured strain distribution was heterogeneous initially and became even more heterogeneous with collagen degradation (**Fig.5B**). Nonetheless, after a lag period (also per **Fig.4C**), the instantaneous degradation rate of collagen was slower within strained tendon sublengths (**Fig.5C**).

**Fig.5.**
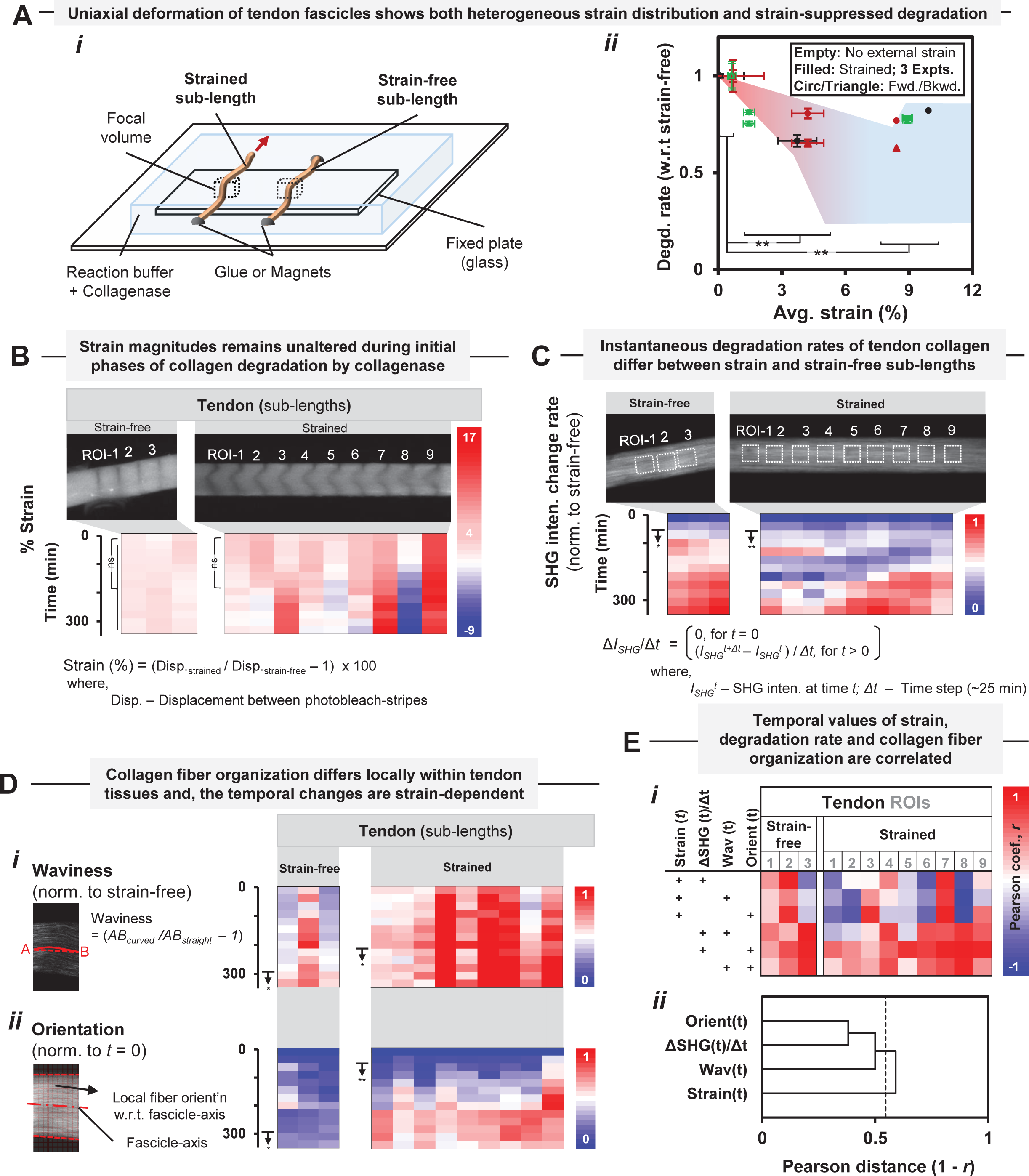
Collagen fiber degradation rate relates to local collagen orientation within a tendon under strain. **A.** (i-ii) Uniaxially deformed “cell-free” mice tendon fascicle sublengths (i.e., strained and external strain-free) under exposure to BC show strain-suppression of collagen degradation rate that is similar to heterogeneously deformed fascicles (for methods, see Fig.4). * Two-tailed t-test: ns = not significant, * = p < 0.05, ** = p < 0.01, *** = p < 0.001; n = 3 samples; Error bars indicate ±SEM. **B.** Temporal local strain distribution within uniaxially deformed tendon remains unaltered only during initial phases of collagen degradation by BC while the degradation do not affect strain distribution in a strain-free tendon. * Two-tailed t-test: ns = not significant, *= p < 0.05, ** = p < 0.01, *** = p < 0.001; n ≥ 3 regions/sample. **C.** The instantaneous degradation rates in the strained sub-length are significantly lower (*p* < 0.05 for >60 minutes) than strain-free counterpart. SHG intensity (*I_SHG_*) is product of collagen fibril density (*N_s_*) and fiber orientation with regard to incident light polarization direction (*γ*). * Two-tailed t-test: ns = not significant, * = p < 0.05, ** = p < 0.01, *** = p < 0.001; n ≥3 regions/sample. **D.** Larger temporal increases in local values of both waviness and orientation in the strained sub-length than their strain-free counterpart during collagen degradation demonstrate strain dependence of collagen fiber organization changes. For waviness and orientation calculations, see supp.-methods S6,S7. * Two-tailed t-test: ns = not significant, * = p < 0.05, ** = p < 0.01, *** = p < 0.001; n ≥3 regions/sample. **E.** (i-ii) Temporal values of different parameters i.e., strain, instantaneous degradation rate and collagen fiber organization, show best correlation between instantaneous degradation rate and fiber orientation. Correlation is defined by Pearson correlation coefficient (*r*) values and shown as color-map. Parameter-wise average *r*-value are used for agglomerative hierarchical clustering based dendrogram with full linkage. The transition between positive and negative *r*-value for a tissue region likely indicates internal stress relaxation due to local collagen degradation.

Organization of collagen fibers (or fibrils) within a tendon during degradation by BC is characterized most simply by waviness and orientation (**Fig.S12,S13;supp.-methods S6,S7**). Waviness, a measure of fiber bending, is initially heterogeneous within both strain-free and strained sublength of tendon tissues (**Fig.5D-i**) but the strained sublength showed rapid increases in waviness values with degradation. Orientation value measures the fraction of collagen fibers inclined with respect to tendon loading-axis, and strained sublengths showed more pronounced increases in local orientation than strain-free regions (**Fig.5D-ii**), with mostly inclined fibers within strained sublengths following degradation (**Fig.S14**). The various spatio-temporal values of strain, collagen degradation rate, and organization were assessed for correlations, and the best correlation is observed between degradation rate and fiber orientation (**Fig.5E**). Orientation and strain are less correlated, consistent perhaps with the idea that more inclined fibers are under low strain and degrade faster as compared to collagen fibers within strain-free regions that degrade almost uniformly irrespective of their orientation (**Fig.S13**).

### Crosslinked fibers’ strain-dependent triple-helical unfolding and degradation

To elucidate strain-dependent multiscale structural changes in a tissue underlying strain-dependent collagen degradation, we began with characterizing orientation of collagen fibers in strained tendons to understand the source of strain heterogeneity and measured the offset between collagen fiber orientation distributions in experimentally-observed and affine-predicted deformed configurations (**Fig.S12,S15A,S16**; **supp.-methods S6**)[51]. The offset provides a measure of non-affine deformation and varies linearly with the fiber orientation relative to tendon-axis (**Fig.S15A-ii**), and likely contributes to the heterogeneity. Globally averaging the fiber inclination over tendon tissue length (∼500 *μ*m) recovered the expected affine deformation.

To address whether the locally complex, non-affine fiber kinematics directly influences collagenase molecules, we considered the strain effects on molecular permeation, collagen fiber density, and molecular mobility. Briefly, collagenase-size fluorescent-dextran molecules (Stoke’s radius, *R*_s_ = 1.4 nm) showed no significant change in spatial distribution within tendon for up to ∼8% strains (**Fig.S15B,S17**). Neither collagen concentration (**Fig.S18**) [52] nor the volume of each tendon region (calculated based on SHG intensity) deviated systematically under strain (**Fig.S15C**), and the product of collagen concentration and volume ratio as tissue mass density reflected the same null result (**Fig.S19**). Notably, SHG measurements avoid limitations of fluorescence photobleaching/quenching, non-specific binding, fluorophore-degradation, etc. (**Fig.S19,S20**). Fluorescence recovery after photobleaching (FRAP) measurements for fluorescent-BC and similar-size fluorescent-dextran (**Fig.S21,S22**; **supp.-methods S8,S9; Table-S3** [53]) showed no strain-dependent mobility for up to ∼8% strain (**Fig.S15D**), whereas larger-size dextran (*R*_s_ = 16 nm) showed strain-suppressed mobility (**Fig.S22**). Interestingly, strains greater than ∼8% increased the mobility of fluorescent-BC and small-dextran molecules (**Fig.S22**), which indicates any suppression of protease degradation at such high strains is offset by increased mobility and higher activity of the protease. Hence, physiological strain-dependent degradation of collagen fibers seems unaffected by properties of the collagenases but controlled locally by the strained fibers as substrate of the enzymes.

We further hypothesized that strain-dependent collagen fiber molecular conformation impedes access to collagenases. To test this hypothesis, artificial collagen crosslinks were first introduced via exogenous transglutaminase (TGM) (**Fig.S23**) to restrain intermolecular sliding within a strained collagen fibril [47, 54, 55] and to subsequently lead collagen triple-helix unfolding [45, 56, 57] favoring degradation by collagenase in contrast to folded triple-helical collagen [50, 58, 59] [60]. Thus, in the uniaxially strained tendon exposed to collagenase, we measured (i) the unfolded collagen triple-helical content by CHP labelling [45] (**Fig.6A**) and (ii) collagen degradation by SHG. In contrast to constant unfolded triple-helical collagen levels in tendon tissues without TGM-crosslinks under uniaxial deformation within the physiological strain regime (**Fig.S24**), TGM-crosslinks carrying heterogeneously strained tendons (**Fig.S25A-B**) showed an increase of ∼3-fold in both unfolded collagen and collagen degradation rate, which are linearly related (**Fig.6B**). Thus, the results are consistent with a model of strain-induced collagen triple-helix unfolding that favors degradation.

**Fig.6.**
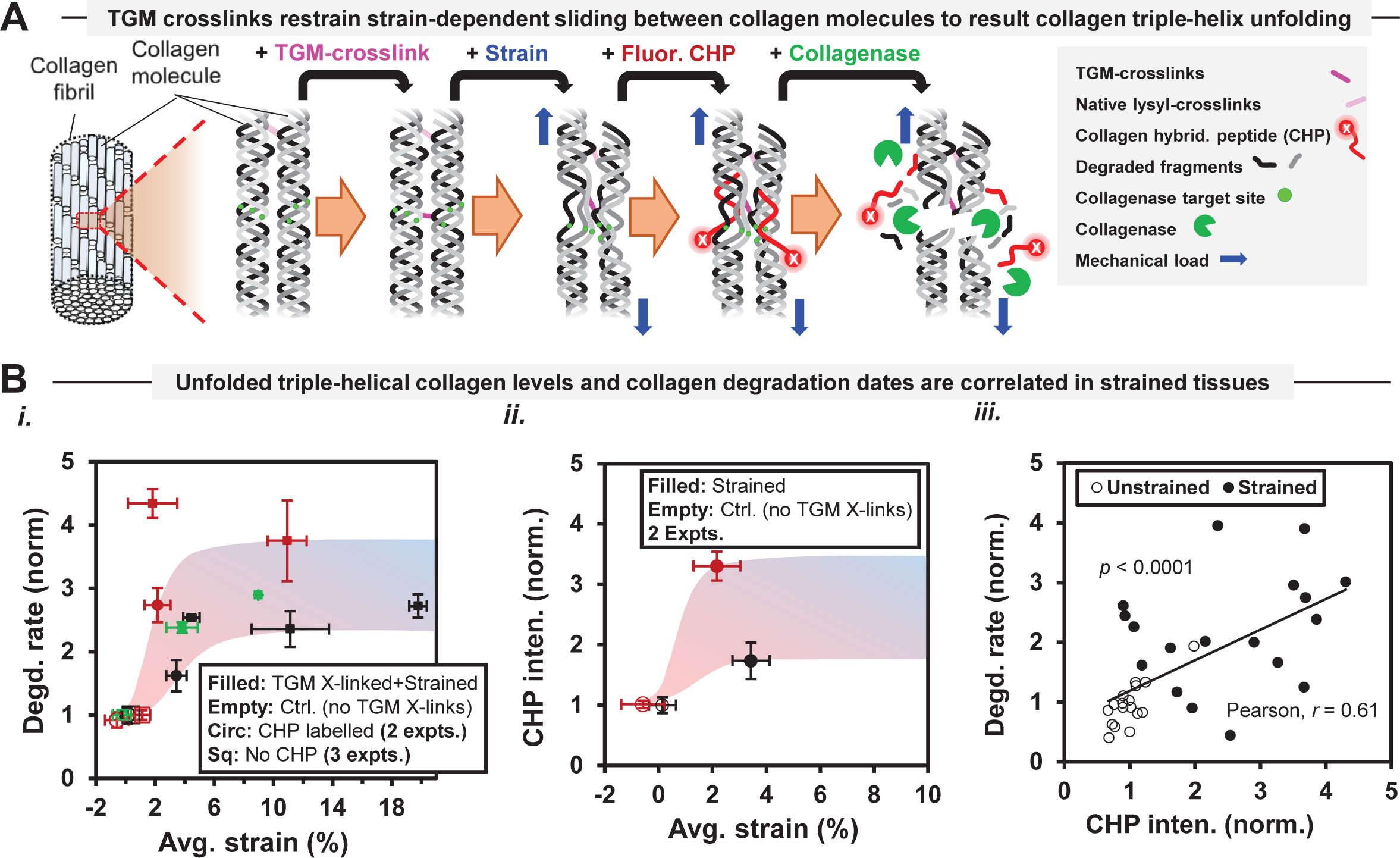
Strain-dependent molecular unfolding and collagen degradation are positively correlated in tendons with increased collagen crosslinks. **A.** Schematic of collagen molecular level structure alterations within a collagen fibril when exposed to transglutaminase (TGM)-crosslinks, mechanical strain, fluorescent collagen hybridizing peptide (CHP) and collagenase molecules. Exogenous TGM molecules induce artificial crosslinks (in addition to the native crosslinks) and, unfolded triple-helical collagen levels are assessed based on binding of fluorescent-CHP. **B.** The presence of TGM-crosslinks increased both collagen degradation rate and unfolded triple-helical collagen levels in uniaxially deformed tendon fascicles. (i) Fascicles carrying TGM-crosslinks show a >3-fold sigmoidal increase in collagen degradation rate by BC as compared to strain-free counterparts (without TGM crosslinks) serving as negative control. n = 3 samples; Error bars indicate ±SEM. (ii) A >3-fold sigmoidal increase in unfolded triple-helical collagen levels in the strained fascicles with TGM-crosslinks is demonstrated by the binding of fluorescent-CHP. n = 2 samples; Error bars indicate ±SEM. (iii) Collagen degradation rate values and unfolded triple-helical collagen levels are positively correlated (Pearson, *r* = 0.61) while both collagen degradation rates and unfolded triple-helical collagen levels in TGM-crosslinked fascicles significantly differ from their strain-free counterparts (with *p =* 0.000024 and 0.000045 respectively based on two-tailed *t*-test).

## Discussion

### Stiffness scaling explains Organ-size scaling via Mechanosensitive transcript levels

Stiffness scaling with SHG signal based fibrillar collagen levels (**Fig.2**), which captures strain-controlled collagen degradation, seems capable of explaining systematic differences among tissues with molecular level sensitivity. In particular, we find that for different organs/tissues among mammalian species (**Fig.7A**), the scaling exponent (*β*) in (Organ-size) ∼ (Body-weight)^*β*^ increases with both the apparent stiffness of tissues *E*_tissue_ (*R*^2^ = 0.95) and fibrillar collagen content (**Fig.7B**). Such change (∼40%) in *β* values among tissues likely results from lower water (or higher protein) levels in stiff tissues compared to soft tissues (**Fig.7C)** given that proteins are ∼40% more dense than water. Thus, the scaling of organ size with body weight from mouse to elephant indicates stronger scaling in stiff load-bearing tissues such as bone and muscle compared to soft tissues such as brain and kidney and, suggests mechanoadaptation of adult tissues.

**Fig.7:**
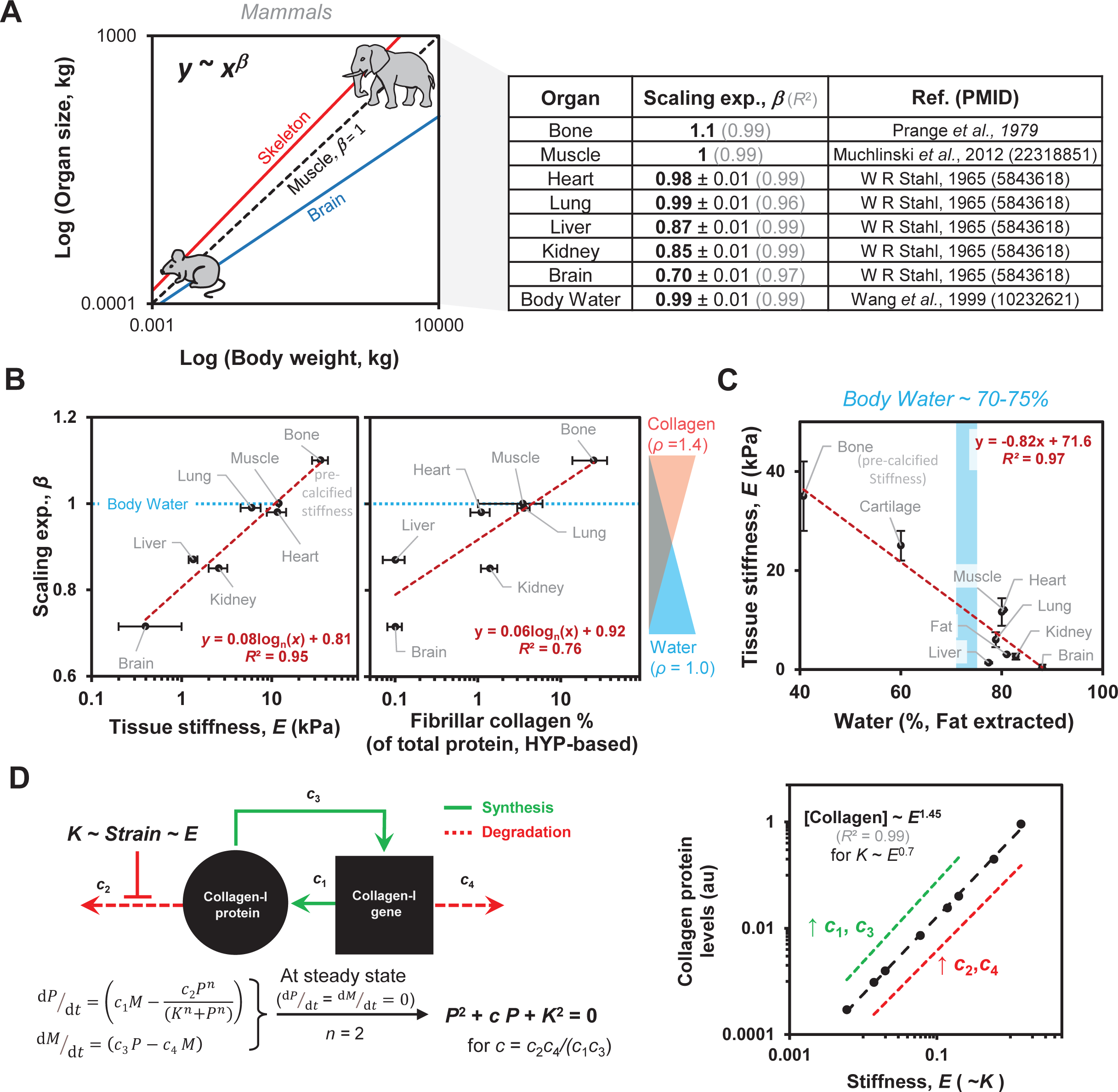
Scaling of organ size with body weight in mammals correlates with tissue stiffness and collagen. **A.** Organ size scales with body weight in diverse mammals (i.e., Organ size ∼ (Body-weight)*^β^* where *β =* scaling exponent). Such scaling is strongest for Bone (*β* = 1.1), weakest for Brain (*β* = 0.7) and other tissues show intermediate values. The scaling of total body water levels with body weight (fat-free or not) is isometric (*β* = 1.0) similar to skeletal muscle tissue (*β* = 1.0) although water level scaling with body weight might differ among tissues. **B-C.** Scaling exponent (*β*) increases with both stiffness and fibrillar collagen (measured based on Hydroxyproline (Hyp) per Tarnutzer et al., 2023) of tissues. Such stronger scaling (*β* = 1.1) in stiff bone carrying high collagen (or less water) levels than soft brain (*β* = 0.7) comprising low collagen (or more water) levels can be expected because proteins in general are 40% more dense than water (density (g/cm^3^), *ρ* = 1.4 for proteins, *ρ* = 1.0 for water). Stiffness decreases linearly with water levels in tissues (fat-free) where total body water levels is ∼70-75%. Data replotted from Wang et al., 1999 (PMID: 10232621) & Smith et al, 2018 (PMID: 29212889). **D.** Positive regulation of collagen-I gene levels by collagen-I protein levels at steady state together with strain-stabilization of collagen protein is sufficient to explain the scaling of Collagen-I with tissue stiffness even varying by several orders of magnitude.

Our hypothesized mechanism for mechanoadaptation [61] is that collagen protein *P* controls its transcript *M* following dynamics of *dP/dt* = *c_1_ M* – *c_2_ P^n^* / (*K^n^ + P^n^*) and *dM/dt = c_3_ P – c_4_ M,* where *n* is Hill Coefficient and *K* is protease affinity towards protein substrate (**Fig.7D**). A strain sensitive *K* reflects strain-stabilizing effect on collagen protein and relates – we postulated – to tissue stiffness as *K* ∼ *E*^0.7^. For *n* = 2 (some cooperativity), the steady state solution gives *P* ∼ *E*^1.45^ (or *E* ∼ *P*^0.7^) which is consistent with **Eq.2**. Although such observations require continued scrutiny, the versatility of power laws in explaining molecular level mechanisms is certainly evident (**Table 1**).

Regardless, the scaling of tissue stiffness with SHG signal from fibrillar collagens in intact tissues is robust in fitting to multiple molecular perturbations and different species (**Fig.1,2**), and it agrees with power laws based on published measurements of Hydroxyproline in fibrillar collagen (**Fig.S25C**). Such relationships also capture physicochemical changes in tissue (defined by stiffness and collagen level) that are common to many pathologies including fibrosis, cancer [62] as well as maturation and adaptation of healthy tissues such as during spaceflight. Our finding that ∼5% tissue strain – rather than stress – impedes the degradation of fibrillar collagens in live tissue ECM by MMPs across soft and stiff adult tissues suggests local degradative sculpting. Such findings on mechanoregulation of fibrillar collagen level kinetics in diverse tissues can be inferred from changes in physical characteristics such as stiffness and cross-sectional area of adult tissue ECM subjected to time-varying mechanical loads [63] [64] [65] [66] [67] [68].

Degradative sculpting of tissues irrespective of stiffness, collagen density and protein synthesis (**Fig.S26A**) define pan-tissue principles as well as a basis for the weak scaling of stiffness with collagen-I levels (i.e. *E* ∼ [Collagen-I]*^a^*) even in cellularized gels. Various cell types (not all) secrete and degrade ECM, and condense stiff ECM more than soft ECM (**Fig.S1**) [6] consistent with increased cellular contractility induced by stiff ECM [2], and the findings here further show that in unstrained directions excess ECM tends to be degraded away by MMPs. The observed collagen level loss in stiff (adult muscle, lung) and soft (embryonic heart) tissue suggest such changes with myosin-II inhibition are more measurable ex-vivo for tissues with “sufficient” collagen levels and cell-contractility, unlike brain tissues. A weak power-law in the steady state level of fibrous protein has thus far been demonstrated with a simple model wherein (i) Protein degradation by MMPs is suppressed by cellular contractility and/or exogenous load driven ECM fibrillar collagen strain, and (ii) protein levels feedback to regulate synthesis by cells [61]. Power law scaling *E*_tissss_ ∼ SHG^m^ (*m* = 1.6 ± 0.2, **Fig.2A**) reflects not only label-free sensitivity to tissue collagen fibers [15, 69, 70] but also distinct effects of cellularization in gels or tissues resulting from factors such as cellular forces, cell type(s) or density, duration, and collagen crosslinks. It leads to the intrinsic scaling result of more basic importance: *E*_tissue_ ∼ [ Collagen-I _SHG_]*^m’^* (*m’* = 0.9 ± 0.1).

Moreover, our findings with purified MMP-1 and BC showing similar times to degrade fibrillar collagen in tissues (**Fig.4D**) contrasts with degradation of soluble collagen-I that is >10-fold faster for BC than MMP-1 (**Fig.S10**) elucidate the differences in cleavage mechanisms including number of target sites per collagen molecule [38, 49]. Pathological strains (i.e., >8%) in tendon producing noisy degradation rates (**Fig.4D**), increased mobile fraction (**Fig.S15D**) and higher unfolded triple-helical collagen levels (**Fig.S24**) collectively suggest multiscale fracture [45, 47] when compared to physiological strains (i.e., ∼5-8%) [45, 47, 54, 55]. Hence, a pan-tissue maximization of collagen fibril lifetime against collagenases is tightly regulated by tissue strain (**Fig.4E**).

Although local degradation rate correlates well with local fiber orientation (**Fig.5E**), sudden bursts of local strain relaxation following collagen degradation [71] could explain considerable noise, strain and waviness changes are also reasonably correlated. Furthermore, fiber orientations that underlie non-affine strains likely relate to reorientation of native crimps [44, 72] parallel to tendon-axis (**Fig.S15A**) and reflect how fibers locally share and transfer load [73]. The evident sensitivity of fiber response(s) to tissue age/health via non-collagenous proteins, load magnitudes, cell density, and pitch of crimps [72, 74] together highlight the remarkable, emergent simplicity of the observed scaling. Conformational control of collagen degradation could explain the lack of a strain effect on tissue permeation, tissue mass density, and enzyme mobilty (**Fig.S15B-D**), but could also explain the effect of crosslinking by TGM (**Fig.6**). TGM-crosslinks are distinct from native lysyl-crosslinks (**Fig.S23**) and are likely inducing changes in fibrillar surface residues (Glutamine and Lysine) due to TGM’s size (*R_s_* > 2 nm [53]), and TGM-crosslinks influence on tendon tissues seems detectable by SHG signal [75] where tendon tissues with native lysyl-crosslinks show little to no triple-helical unfolding of strained collagen (**Fig.S24**). Our TGM results thus support a model wherein strain suppresses conformational accessibility to collagenases in tissue ECM.

### Applications of Strain-inhibited Sculpting of Materials

The scaling results here can help explain the sensitivity of differentiation to ECM stiffness [76], while noting that both structure and stiffness develop at multiple length-scales in proportion to sustained physical loads via mechanisms associated with ECM collagen (i.e.levels, crosslinking, alignment), calcification, etc. and some are captured by SHG signal (**Fig.1,2**). Moreover, strain-inhibited degradation of fibrillar collagen by MMPs can help explain past observations of improved tissue mechanics upon inhibition of endogenous MMPs [77]. Such effects synergize with mechanoregulation of collagen and MMP transcription via cytoskeletal tension on long timescales of ∼days [78] [79] [80] [81] [82] [83] [84] [85] [86]. They also synergize with stimulation of synthesis by exercise [80, 87] and non-physical factors [88] [89].

Perturbations on short timescales studies here are particularly relevant to heart dysfunction associated with an acute heart attack or infarct. However, contributions in other tissues of MMPs, collagen crosslinking enzymes (TGM, LOXs) and collagen synthesis with different perturbations on long times scales of days or weeks require further investigation. Understanding such tissue process by SHG will likely benefit from deconvolution of optical scattering effects [90], of contributions from different collagen-crosslinks [75], and estimations of surface/core area based on fibril size distributions [15]. Findings here could help assess and perhaps treat fibrosis and even solid tumors [91–96], and potentially limit atrophy in the microgravity of spaceflight (e.g. with MMP inhibitors). Therapeutic targeting of cellular contractility (per our study on several tissues and some past studies [67] [68]) could also help regenerate or rejuvenate tissue ECM, and SHG among other label-free imaging methods also show useful signal differences (**Fig.S26B**). Although scaling over many logs shown here is useful as a holistic approach, more limited perturbations to a single tissue might not clearly show significant tracking along the broader trend. Tissue constructs for regenerative medicine and broader classes of materials might benefit from monitoring the strain-inhibited sculpting that underlies stiffness scaling: simply, if the material at some location is not strained in normal operation, then that material is simply unnecessary and should be removed and recycled for use where needed most.

## Acknowledgments

We appreciate support from the National Institutes of Health National Cancer Institute under Physical Sciences Oncology Center Award U54 CA193417, Human Frontiers Sciences Program grant RGP0024, National Heart, Lung, and Blood Institute award R01 HL124106, SERB Indo-US Postdoctoral Fellowship 2016-157, the US–Israel Binational Science Foundation, National Science Foundation, Materials Science and Engineering Center grant to the University of Pennsylvania and also grant agreement CMMI 15-48571. This content is solely the responsibility of the authors and does not necessarily represent the official views of the National Institutes of Health or other granting agencies. Leica SP8-MP upright microscope at Penn Vet imaging core was supported by NIH grant S10 OD021633-01. K.Y. is supported by Versus Arthritis Career Development Fellowship (Grant 21447). Authors acknowledge J.C. Andrechak, L.J. Dooling, J. Irianto and A.A. Anlas for providing key materials or editing the final draft.

## Author Contributions

Conceptualization: K.S., D.E.D.; Investigation: K.S., S.C.; Formal Analysis, K.S., D.E.D.; Validation: AA.R.J., M.T., M.W.; Key materials/Resources: A.J.K., D.M.C., K.Y.; Writing: K.S.; Editing: K.S., AA.R.J., D.E.D.

---

**Table S1:** Stiffness (*E*) values for embryonic chick and adult tissues.

**A.** Stiffness values for embryonic chick tissues (E2: embryonic day-2, E4: embryonic day-4, etc.)
| # | Tissue | *E* (kPa) | Ref. (PMID) |
| --- | --- | --- | --- |
| 1 | E2-Brain | 0.4 ± 0.2 | Majkut *et al.*, 2013 (24268417) |
| 2 | E4-Heart (collagenase treated) | 0.738 ± 0.078 | Majkut *et al.*, 2013 (24268417) |
| 3 | E4-Heart (myosin-II inhibited) | 0.85 ± 0.11 | *Present work* |
| 4 | E4-Heart (untreated) | 1.23 ± 0.13 | Cho *et al.*, 2019 (31105008) |
| 5 | E6-Heart (untreated) | 1.7 ± 0.33 | Cho *et al.*, 2019 (31105008) |
| 6 | E10-Heart (untreated) | 3.78 ± 1.0 | Cho *et al.*, 2019 (31105008) |

**B.** Stiffness values for adult tissues (per Swift *et al.*, 2013)
| # | Tissue | *E* (kPa) | Reference (PMID) |
| --- | --- | --- | --- |
| 1 | Brain | 0.4 (+0.6 / -0.2) | Georges *et al.*, 2006 (16461391) |
| 2 | Liver | 1.35 ± 0.15 | Georges *et al.*, 2007 (17932231) |
| 3 | Kidney | 2.6 ± 0.6 | Wyss *et al.*, 2011 (21123730) |
| 4 | Lung | 6 ±1.5 | Lai-Fook *et al.*, 1985 (10904048) |
| 5 | Muscle | 12 | Engler *et al.*, 2006 (16923388) |
| 6 | Heart | 11.6 ± 2.8 | *Present work* |

**Table S2:**
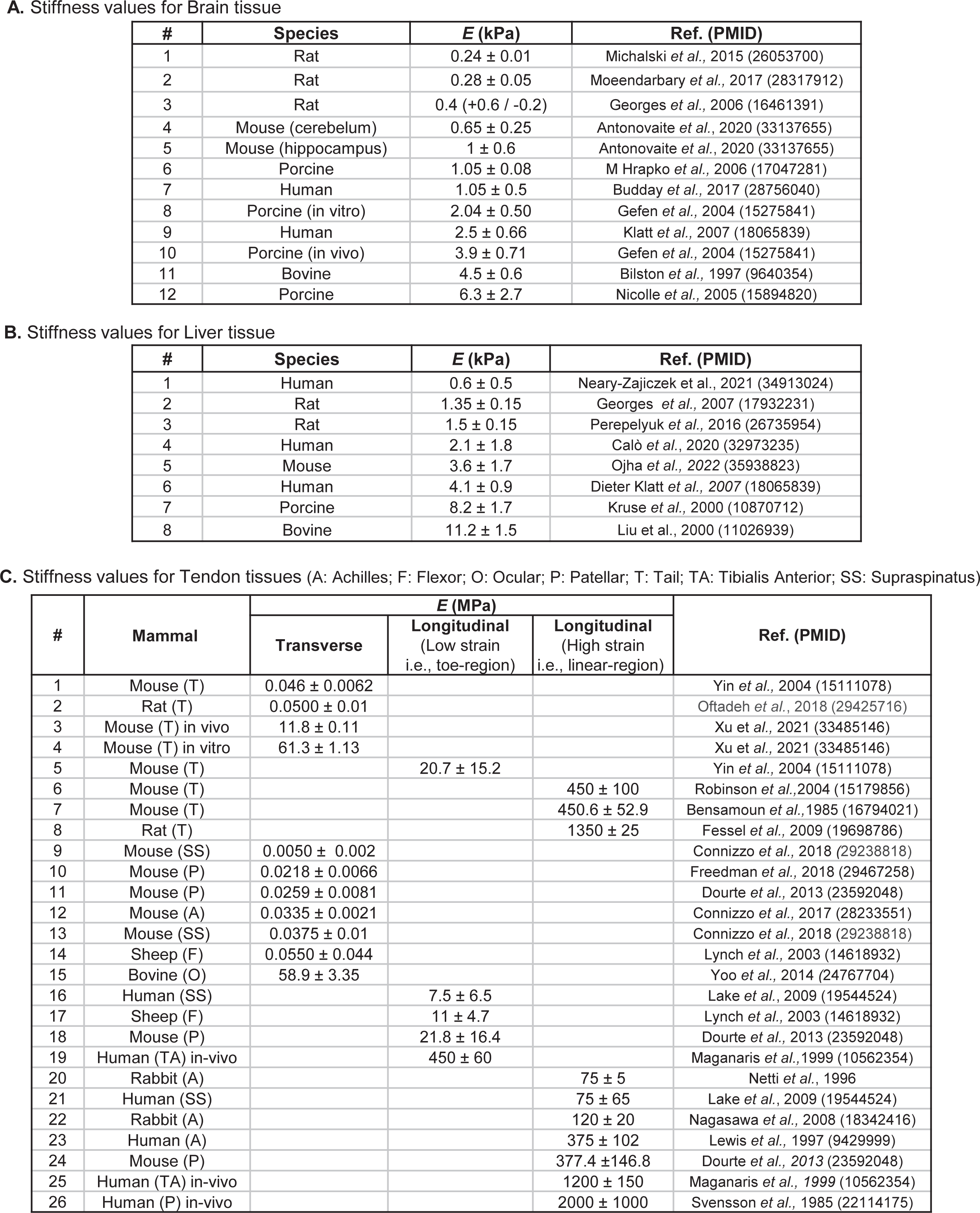
Previously reported stiffness (*E*) values for Brain, Liver and Tendon tissues from different species.

**Table S3:** Size of BC and dextran molecules.

| **S. No.** | **Item** | **Mass (kDa)** | **Stoke's radius, $R_s$ (nm)** |
| --- | --- | --- | --- |
| 1 | BC | 60–130 | ~3 |
| 2 | Dextran TRITC | 4.4 | 1.4 |
| 3 | Dextran TRITC | 155 | 8.5 |
| 4 | Dextran TRITC | 500 | 16 |

**Table S4:**
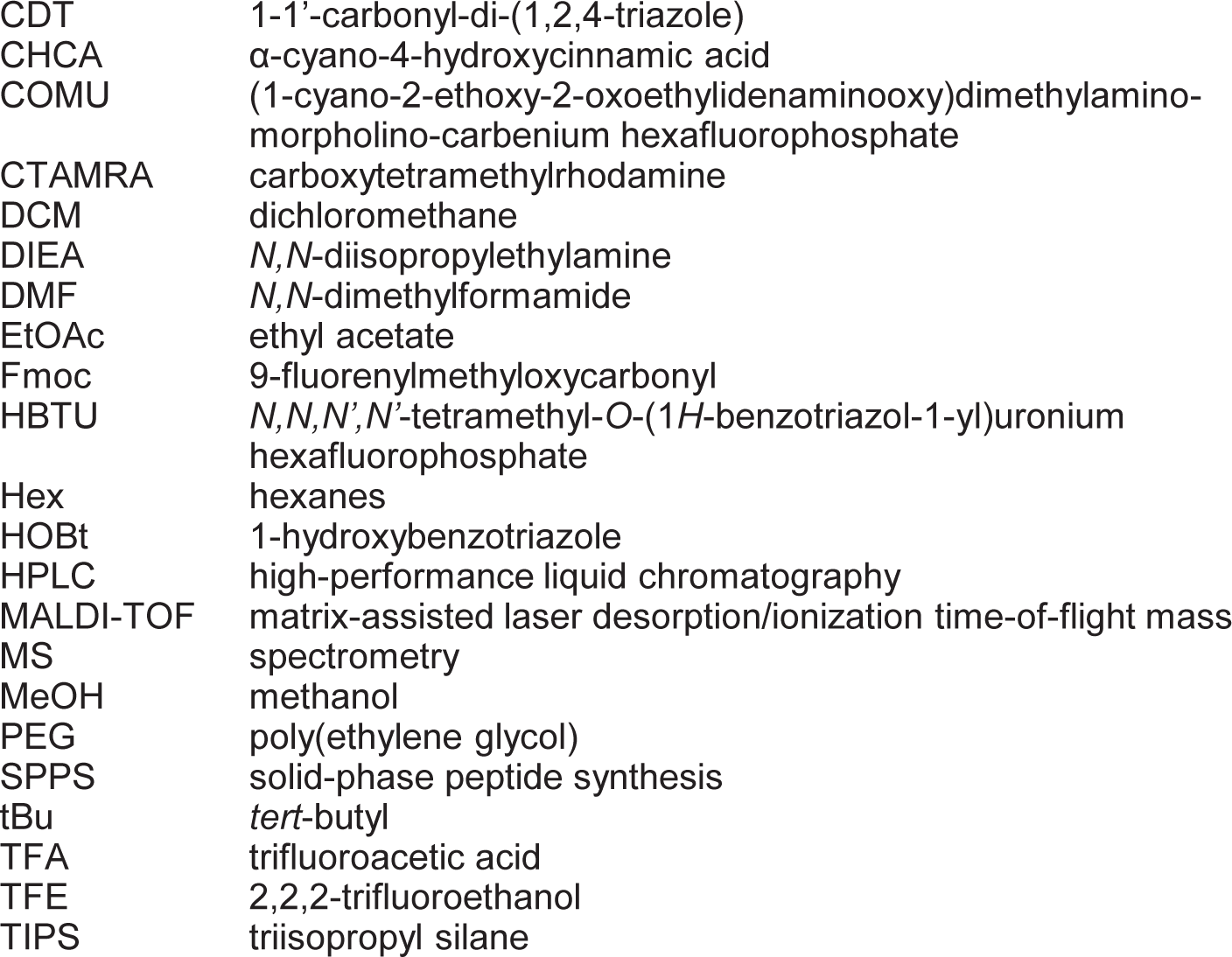
Abbreviations.

## Materials and Methods

### Embryonic chick hearts

White Leghorn chicken eggs (Charles River Laboratories; Specific Pathogen Free (SPF) Fertilized eggs, Premium #10100326) were used to extract embryonic hearts. SPF chicken eggs from Charles River are produced using filtered-air positive-pressure (FAPP) poultry housing and careful selection of layer flocks. Every flock’s SPF status is validated in compliance with USDA memorandum 800.65 and European Pharmacopoeia 5.2.2 guidelines. SPF eggs were incubated at 37°C with 5% CO_2_ and rotated once per day until the desired developmental stage (e.g., four days for E4; Hamburger-Hamilton stage 23-24 (HH23-24)). Embryos were extracted at room temperature (RT) by windowing eggs, carefully removing extra-embryonic membranes with sterile sonicated forceps, and cutting major blood vessels to the embryonic disc tissue to free the embryo. The extracted embryo was then placed in a dish containing PBS on a 37°C heated plate, and quickly decapitated. For early E2-E5 embryos, whole heart tubes were extracted by severing the conotruncus and sino-venosus [8]. All tissues were incubated at 37°C in pre-warmed chick heart media (α-MEM supplemented with 10% FBS and 1% pen-strep, Gibco, #12571) for at least 1 hour for stabilization, until ready for use. BC (Sigma-Aldrich, C7657) comprising multiple enzymes was used as an exogenous collagenase in chick heart experiments. Actomyosin perturbation was done by blebbistatin (EMD Millipore, #203390), protein synthesis was inhibited by blocking translation step using cycloheximide (Sigma, #C7698) and MMP activity was inhibited by ilomastat (GM6001). For single-cell RNA-seq, the hearts were disaggregated using dispase (Corning, Catalog no. 354235) supplemented with 4 mg ml-1 collagenase IV (Thermo Fisher Scientific, Catalog no. 17104-019) and DNAse I (Thermo Fisher Scientific, Catalog no. 18068-015) at 1 *µ*L per 1 mL of dispase solution. Hearts were allowed to dissociate for 30-45 min (until mostly dissociated) while incubated at 37°C. Dissociated cells were centrifuged at 300Xg, washed with Dulbecco’s phosphate-buffered saline (PBS; Gibco, Catalog no. 10010-023), and then gently passed through a 40-micron filter to further dissociate suspension and remove extracellular material. Cells were washed once more and then used for single-cell sequencing experiments.

### Mouse tissues

In accordance with institutional animal care and use procedures, euthanized wild-type (WT) adult mouse (C57BL/6J, 9-12 weeks) were selected for extraction of the tissues. For inter-tissue comparison, tail tendon and other tissues namely skeletal muscle (from hindlimb), heart (ventricle region), brain (cerebrum) and lobes of lung, liver and kidney were isolated, kept overnight in 4% PFA, stored in 70% ethanol (4°C) and imaged within 10 days. For tendon fascicles extraction, the tails were sectioned between the coccygeal vertebrae at the base and distal tip of the tail using a sterile scalpel blade to result lengths of up to ∼70 mm. The tail was immediately washed and placed in PBS (Corning 21-031-CV) at room temperature for extraction of the fascicles using a sharp tweezer. For enzymatic reaction purposes, each group of 3-4 isolated fascicles were kept in small Eppendorf tubes carrying PBS and stored at −20°C. Before start of each experiment involving enzymes, the stored fascicles were thawed at 4°C and washed few times in PBS at room temperature. For enzymatic reactions involving bacterial collagenase (BC i.e., collagenase from Clostridium histolyticum (Sigma-Aldrich, C7657)), PBS with Ca/Mg (Thermo Scientific, 14040117) was used as a reaction buffer. For experiments involving purified and activated MMP-1, 50 mM Tris-HCl/ 150 mM NaCl/ 10mM CaCl_2_ with 0.01%, bovine serum albumin (BSA) solution were freshly prepared for use as reaction buffer [38, 97]. DQ™ Collagen, type I From Bovine Skin, Fluorescein Conjugate (Thermofisher, D12060) was used for comparing collagenase activity of MMP-1 and BC. For experiments involving treatment with Trypsin (Sigma-Aldrich, T4174), tendon samples were incubated in 10x trypsin solution for ∼48 hours at 37°C. Fluorescent labelling of fascicles was done using Aza-peptide ([5(6)-CTAMRA-NH(PEG)_3_CO-(azGPO)_3_-NH_2;_ M.W. = 1481.58 Da) or Col-F dye (Immunochemistry Tech., 6346). Dextran solutions of 4.4 kDa, 155 kDa or 500 kDa (Sigma Aldrich, T-1037, T-1287, 52194) (0.1 mg/mL in PBS) were used for FRAP studies. Fluorescent-BC (0.01 mg/mL in PBS) was prepared (**supp.-methods S8**) by labelling Clostridium histolyticum (Sigma-Aldrich, C7657) at N-terminus using Alexa Fluor™ 555 NHS Ester (Invitrogen™, A20009) per manufacturer protocol. For artificial crosslinking, the tendon fascicles were incubated in ∼2.5 mg/mL transglutaminase (TGM) (Sigma-Aldrich, T5398) solution in DMEM (Thermo Scientific, 21063029) having 10% FBS + 1% P/S. For detection of unfolded collagen within samples, Biotin conjugate of collagen hybridizing peptide (CHP) (3Helix, Inc., B-CHP) and fluorescent Streptavidin conjugate Alexa-594 (Thermofisher-S11227) were used following endogenous biotin blocking using Biotin-blocking kit (Life Technologies Corporation/Thermo fisher, E21390). For Aza-peptide preparation, the following reagents/ solvents were used as received: 2-chlorotrityl chloride resin (ChemPep.), Low-loading rink amide MBHA resin (Novabiochem), Fmoc-Pro-OH, CDT, and Fmoc-NH(PEG)_3_-COOH (Chem-Impex); Fmoc-Hyp(tBu)-OH (AK Scientific); COMU CreoSalus (Advanced ChemTech); DIEA (Acrōs); TFE, TIPS, HBTU, and Fmoc-carbazate (Oakwood); Piperidine (EMD Millipore); 5(6)-CTAMRA (Carbosynth/ Novabiochem); TFA (Alfa Aesar); all other solvents were acquired from Fisher or Sigma.

### Tissue stiffness measurements

Following application of internal pressure (*P*_in_), tissue aspiration into glass micropipettes (measured by aspirated length *L,* **Fig.1B-i**) having inner diameter (*R*_p_ = ∼50 *µ*m), which is sufficient to probe dozens of cells plus ECM, was used to quantify tissue stiffness. Given that control adult heart tissue deformed instantaneously when compared to collagenase treated hearts (taking tens of seconds) based on strain (*ε = L*/*R*_p_) values (**Fig.1B-ii**), adult heart tissue behaved elastically compared to inelastic behavior of collagenase treated hearts under similar applied suction pressure. Considering instantaneous tissue strain (within initial ∼3 seconds) and applied suction pressure (Δ*P* = *P*_out_-*P*_in_, where *P*_out_ = atmospheric pressure), stiffness values (*E* ∼ Δ*P*/*ε*) were measured for adult tissues (**Fig.1**) and embryonic heart tissues (**Fig.S3**).

### Environment control mechanical manipulation of the fascicles

To mechanically deform tendon fascicles, the fascicles obtained following several freeze-thaw cycles (**Fig.S4**) were placed the mechano-bioreactor was installed on a MP microscope following their fluorescent labelling (**Fig.S5**; **supp.-methods S1**). Due to smaller displacement between tissue sample and the lens, SHG signal was captured easily in both backward and forward directions using the mechano-bioreactor. The movable plate facilitates in-plane deformation of the sample via three-point bending and subsequent exposure to collagenase or dextran molecules. The temperature is maintained at 37°C by the bottom heating plate using a separate temperature control unit. Notably, our measurements are unaffected by the evaporation losses of the reaction buffer from the mechano-bioreactor edges (**Fig.S6;supp.- methods S2**). For uniaxial deformation tests (**Fig.5A-i**), each fascicle was cut into strain-free and strained sub-lengths and was kept on a glass surface to avoid the adhesion contact friction. The samples were immersed in reaction buffer medium during the experiment. The strain-free sub-length was glued at both ends to avoid deformation whereas to-be-strained sub-length was glued at one end while other end was subjected uniaxial pulling and holding.

### Fluorescent labeling of the fascicles

To quantify the strain magnitudes, the fascicles were fluorescently labelled with either Aza-peptide or Col-F dye (see methods). For fluorescent labelling with Aza-peptide, the fascicles were kept in 30 *μ*M Aza-peptide solution in PBS (for synthesis and purification of Aza-peptide, see methods) [98]). Aza-peptide solution was heated to 80°C for 5-7 minutes (to obtain monomeric peptide) and quickly cooled to 4°C (by keeping on ice for ∼25 sec) before application to the fascicles by an overnight incubation at 4°C (**Fig.S7**). The low molecular weight fluorescent probe i.e., Col-F (fluorescein conjugated to Physostigmine), exhibiting non-covalent affinity towards ECM fibers [99], was used also to label the fascicles. Fascicles were kept in Col-F dye solution (at ∼50 *μ*M) for 30 minutes at room temperature, followed by at least two PBS washes of 30 minutes each using tube revolver to remove the free dye molecules followed by immediate use (or kept on ice in longer waiting durations).

### Strain quantification by fluorescent pattern photobleach (FPP) method

To measure strain distribution within fluorescently labelled tendon fascicles (**Fig.S8**), FPP method was used. A half-diameter deep region into the fascicle was selected to create ten equally spaced rectangular photobleach-stripes (∼5 *μ*m width with ∼50 *μ*m inter-stripe spacing) using FPP method. Given that fluorescent intensity in each photobleach-stripe region followed a Gaussian distribution, Gaussian peak was used to represent each stripe position within undeformed and deformed configuration of the fascicle. The displacement between each pair of photobleach-stripes were measured on a projected sum composite image along a path parallel to the longitudinal axis of the fascicle (i.e., thick yellow line in **Fig.S8B-ii**). A fixed number of points on the ROI path were selected for Gaussian curve fitting using custom-built MATLAB script (**supp.-methods S3**).

### Second harmonic generation (SHG) imaging

An upright multi-photon microscope TCS SP8 MP (Leica) was utilized for SHG imaging of native tissue structure with minimal processing (e.g., paraffin embedding, sectioning, etc.). While backward SHG signal was captured at different locations of each tissue and used for absolute SHG intensity comparison among tissues, corresponding forward SHG could only be obtained in tissue regions with less thickness (< ∼200 *µ*m). The tunable Coherent Chameleon Vision II Ti:Sapphire laser (680-1080 nm) produced linearly polarized pulses spectrally centered at 910 nm of 1.99 W power. Incident light at 910 nm was focused onto the sample with a 20X water immersion HCX APO L (1.00 NA) objective for backward SHG signal. Due to collagen fibril size dependent emissivity [100], SHG signal was additionally captured in forward direction (by a 0.9 NA condenser lens). The laser power was adjusted to obtain SHG signal without any obvious damage to samples [15, 101]. Each z-stack acquired tissue volume (2 *µ*m spacing) with 200 Hz line frequency using LAS X (Leica Application Suite X).

### Stiffness scaling with SHG signal at steady state captures strain-inhibited degradation

Assuming first-order collagen-collagenase interactions, with collagen concentration defined in terms of SHG signal ([Col] ∼ [SHG]^n^, n = 1.6 for collagen-I per **Fig.S1**), and collagen degradation rate (- *Δ*[Col]/*Δt*) and its strain-suppression respectively defined by

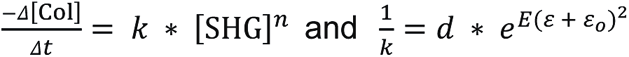

where, *ε_o_* and *ε* are set-point strain and applied strain values (*ε* ≥ *ε_o_*) respectively that suppress collagen degradation rate in a tissue of stiffness (*E*) via modulating reaction rate constant (*k*), and *d* is a constant. As collagen synthesis rate equals its degradation rate at steady state of a tissue, and assuming constant collagen synthesis rate (*s*) in a tissue

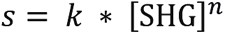

Taylor series expansion at smaller values (say ∼1% strain and *E* in MPa) gives

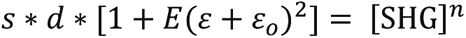

For each strain value (e.g., 5% strain as observed in E4-chick hearts), and considering *c* = *s* x *d* x (*ε* + *ε_o_*)^2^ gives

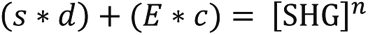

Thus, if (*s* x *d*) << (*E* x *c*), then

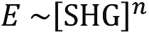

which shows that stiffness scaling with SHG signal is preserved during strain-suppression of collagen degradation rate.

### Evaluation of collagen fiber deformation behavior and degradation dependent fascicle structure changes

To understand the relationship between the fascicle structure and tissue mechanical strains, we evaluated collagen fiber kinematics within different regions of the fascicles. SHG signal was used to calculate local collagen fiber orientation within the fascicles based on the structure tensor (**supp.-methods S6**). The fiber orientation distribution within undeformed configuration and strain magnitude of a ROI (i.e., region between a pair of consecutive photobleach-stripes of the fascicle) were used to predict theoretical collagen fiber orientation distribution in ROI’s deformed configuration based on affine deformation assumption. The following equation was incorporated into a customized script in MATLAB for making the calculations

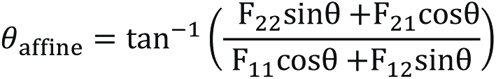

where, *θ* represents collagen fiber orientation with respect to the fascicle-axis in the undeformed configuration and *F*_ij_ are elements of deformation gradient matrix of the ROI. For a deformed ROI within a fascicle, *F*_11_ represents the change in displacement between the photobleach-stripes (along longitudinal-axis of the tissue) while F_22_ represents the change in ROI size in the transverse direction (i.e., fascicle-radius change based on averaged transverse cross-sectional area (CSA) along fascicle-axis). F_21_, F_12_ were assumed negligible (adjusted to zero) due to both uniaxial deformation of the ROIs along the fascicle-axis and parallelism of the photobleach-stripes following ROI deformation. Experimentally observed and affined predicted fiber orientation distributions in deformed configuration of a fascicle region were compared offset values obtained using projection plots (**Fig.S12;supp.- methods S6**) [51] in order to quantify the extent of affine deformation.

### Fluorescence recovery after photobleaching (FRAP)

Mobility of various size solute particles within mechanically strained regions of the fascicle were quantified using FRAP method. Here, fluorescent-BC molecules labelled at N-terminus (see methods) or several size TRITC-dextran molecules (**Table-S3**) were used (**Fig.S15D,S22**). The deformed fascicles carrying a solute solution were kept overnight at 37°C (protected from light and evaporation) before making FRAP measurements. For each solute solution (fluorescent BC or each size TRITC-dextran), the sub-volumes of the fascicles carrying different strain values were selected for mobility measurements. FRAP data from each strained sub-volume of the fascicle was normalized and binned using a custom MATLAB script. The obtained data was then used for determination mobility characteristics i.e., mobile fraction (m) and time constant (t_1_), using a single exponential fitting (**supp.-methods S9**).

### Fluorescent labeling of BC molecules at N-terminus

For determining the mobility of collagenase-size molecules in a mechanically deformed fascicle, fluorescent labelling of BC molecules at their N-terminus was done using Alexa Fluor™ 555 NHS Ester (**Fig.S21**;**supp.-methods S8**). Here, BC molecules and fluorophore (Alexa Fluor™ 555 NHS Ester) were reacted at two different stoichiometric ratios. The labelling efficiencies were observed using UV (ultraviolet) scanning and high performance liquid chromatography (HPLC) measurements.

### Artificial crosslinking by transglutaminase (TGM)

For determining the influence of collagen molecular structural perturbations on strain-dependent collagen degradation, the fascicles were artificially crosslinked using ∼2.5 mg/mL transglutaminase (TGM) solution in DMEM (with FBS+P/S) for ∼24 hours incubation at 37°C. After TGM treatment, the samples were washed twice with PBS before further use. One sub-length of each fascicle was crosslinked using TGM while the other one served as a strain-free control without TGM crosslinks.

### Detection of unfolded collagen content within fascicle samples

For detection of unfolded collagen within mechanically deformed the fascicles, Biotin conjugate of collagen hybridizing peptide (CHP) was used per manufacturer’s protocol while fluorescent streptavidin conjugate Alexa-594 was used for its detection. Briefly, following endogenous biotin blocking of the fascicles, Biotin-CHP stock solution (50 *μ*M) was heated at 80 °C for 10 min to thermally dissociate trimeric CHP to its monomeric state, and quenched by immersion in 4 °C water for ∼15 seconds. Each fascicle sample was kept in 10 µM CHP solution at least for ∼2 hours at 4 °C followed by twice washing with PBS. The fascicles were then incubated in fluorescent streptavidin conjugate Alexa-594 solution in dark for ∼30 minutes and were subsequently washed twice with PBS. Fluorescent imaging of tissue bound CHP was made using two-photon laser (excitation: 810 nm, emission: 610-660 nm) by creating z-stacks through the tissue depth. Composite images obtained using projection of the z-stacks were used to quantify fluorescence from the CHP, and intensities were averaged over rectangular areas (at least 50 *µ*m x 50 *µ*m) for making further analysis.

### Mass spectrometry (LC-MS/MS) of whole heart lysates

Mass spectrometry (LC-MS/MS, or ‘MS’) samples were prepared using procedures outlined [2]. Briefly, ∼1 mm^3^ gel sections were carefully excised from SDS–PAGE gels and were washed in 50% 0.2 M ammonium bicarbonate (AB), 50% acetonitrile (ACN) solution for 30 minutes at 37°C. The washed slices were lyophilized for >15 minutes, incubated with a reducing agent (20 mM TCEP in 25 mM AB solution), and alkylated (40 mM iodoacetamide (IAM) in 25 mM AB solution). The gel sections were lyophilized again before in-gel trypsinization (20 mg/mL sequencing grade modified trypsin, Promega) overnight at 37°C with gentle shaking. The resulting tryptic peptides were extracted by adding 50% digest dilution buffer (60 mM AB solution with 3% formic acid) and injected into a HPLC system coupled to a hybrid LTQ-Orbitrap XL mass spectrometer (Thermo Fisher Scientific) via a nano-electrospray ion source. Raw data from each MS sample was processed using MaxQuant (version 1.5.3.8, Max Planck Institute of Biochemistry). MaxQuant’s built-in Label-Free Quantification (LFQ) algorithm was employed with full tryptic digestion and up to 2 missed cleavage sites. Peptides were searched against a FASTA database compiled from UniRef100 gallus (chicken; downloaded from UniProt), plus contaminants and a reverse decoy database. The software’s decoy search mode was set as ‘revert’ and a MS/MS tolerance limit of 20 ppm was used, along with a false discovery rate (FDR) of 1%. The minimum number of amino acid residues per tryptic peptide was set to 7, and MaxQuant’s ‘match between runs’ feature was used for transfer of peak identifications across samples. All other parameters were run under default settings. The output tables from MaxQuant were fed into its bioinformatics suite, Perseus (version 1.5.2.4), for protein annotation and sorting.

### Gel electrophoresis and Coomassie Brilliant Blue stain

Hearts isolated from chick embryos (E4, E6, and E10; n ≥ 8, 4, and 2 respectively) were rinsed with pre-warmed PBS, excised into sub-millimeter pieces, quickly suspended in ice-cold 1x NuPAGE LDS buffer (Invitrogen; diluted 1:4 in 1x RIPA buffer, plus 1% protease inhibitor cocktail, 1% β-mercaptoethanol), and lysed by probe-sonication on ice (10 x 3 second pulses, intermediate power setting). The samples were then heated to 80°C for 10 minutes and centrifuged at maximum speed for 30 minutes at 4°C. SDS-PAGE gels were loaded with 5-15 *μ*L of lysate per lane (NuPAGE 4-12% Bis-Tris; Invitrogen). Lysates were diluted with additional 1x NuPAGE LDS buffer if necessary. Gel electrophoresis was run for 10 minutes at 100 V and 1 hour at 160 V. Gels were then stained with Coomassie Brilliant Blue for >1 hour, then washed using de-stain solution (10% acetic acid, 22% methanol, 68% DI water) overnight.

### Ex-vivo drug perturbations of embryonic hearts

For actomyosin perturbation experiments, intact E4-chick hearts were incubated in 25 *μ*M blebbistatin (stock solution 50 mg/ml in DMSO) and were kept away from light to minimize photo-deactivation. The activity of endogenous MMPs was inhibited by ilomastat while protein synthesis was inhibited by 1 *μ*M Cycloheximide. Treated hearts were compared to a control sample treated with an equal concentration of vehicle solvent DMSO in heart culture media. For collagen perturbations, E4-chick heart tissue was incubated in BC (1 mg/mL) for ∼45 min. Enzyme activity was blocked by replacing with chick heart media containing 5% BSA. A minimum of 8 hearts were treated and pooled per lysate/experimental condition.

### Aza-peptide synthesis and purification

Aza-peptide was synthesized and purified using procedures outlined (**supp.-methods S10**) [98]. Briefly, HPLC was performed using Agilent 1260 Infinity II systems equipped with Phenomenex PFP(2) columns [98]. Flash chromatography for purification of building block Fmoc-azGPO(tBu)-OH was performed using a Teledyne ISCO CombiFlash Rf chromatography system. MALDI-TOF MS was performed using a Bruker MALDI-TOF Ultraflex III mass spectrometer. UV-vis spectroscopy was performed using a JASCO V-650 UV-vis spectrophotometer. Probe aliquots were dried using an Eppendorf Vacufuge plus concentrator for storage prior to use in subsequent assays. Fmoc-azGPO(tBu)-OH was synthesized as described in **supp.-methods S10**.

## Supplementary Methods

### S1. Mechanical manipulation mechano-bioreactor design, and installation on MP microscope

The mechano-bioreactor has glass coverslips of thickness 130–160 *μ*m (cover glasses from Fisherbrand™) as upper plate and middle plate that are glued to each other and bottom polystyrene surface of petri-dish using nail-paint (**Fig.S5**). Together with nail-paint film (thickness ∼40 *μ*m) and side glass plates, a height of ∼220 *μ*m is obtained between bottom polystyrene surface and upper-plate that provides a sufficient room to guide the motion of movable glass plate (VWR 48366-205) for in-plane mechanical deformation of the fascicles (for sizes up to ∼220 *μ*m) without rolling. To place the fascicles into the mechano-bioreactor, wet fluorescently labelled fascicle were gently put on a larger size PBS drop lying on bottom polystyrene surface using sharp tweezers. Careful soaking of the drop reduced its volume ensured minimum residual strains. The sample adherence to the hydrophobic polystyrene surface was evident from the absence of any floating of the sample when PBS was immediately added to maintain sample hydration. Adhesion forces between wet samples and dry polystyrene surface produced local strain gradients within the sample. The rehydrated fascicles following dehydration were found to be degraded by the collagenases (BC or MMP-1). The presence of collagen fiber crimps within SHG image (**Fig.4**) confirmed residual strain-free configuration of the fascicles adhering to polystyrene surface after the transfer from Eppendorf tube to mechano-bioreactor (otherwise, this step wise repeated). Although, the adhesion between fascicle and polystyrene surface was sufficient to retain the deformed configuration of a fascicle sample at least during initial phase of the degradation.

### S2. Reaction buffer composition and packing of collagen fibrils

Due to the use of an upright microscope carrying water-immersion lens, the evaporation losses from edges of the mechano-bioreactor are likely to influence the composition of reaction buffer as well as collagen fibril packing. The evaporation of the reaction buffer carrying different solutes (i.e., BC or dextran) did not change the pH values considerably (**Fig.S6**). Further, the effect of reaction buffer composition changes involving different solutes (i.e., dextran, calcium chloride (CaCl_2_) salt or sodium chloride (NaCl)) on SHG based tissue collagen fibril packing was assessed (**Fig.S6**). Dextran concentration variations did not affect the collagen packing (**Fig.S6B**) but an increase in CaCl_2_ and NaCl concentration in the presence of strain (**Fig.S6C,D**) promoted collagen fibril disassembly and/or signal directionality as indicated by a decrease in forward/backward (F/B) signal ratio [15]. The salt concentration-dependent and mechanical strain-associated changes in the SHG signal are indicative of collagen fibrillar architecture changes as a result of altered molecular packing within a collagen fibril and/or assembly of smaller collagen fibrils into larger fibrils. The compensation for the evaporation losses during an experiment was made by adding DI water at fixed intervals (∼2 hours) based on the evaporation rates of the buffers (**Fig.S6A-ii**).

### S3. Strain quantification with fluorescently labelled tendon fascicles

#### Laser settings for FPP on fascicle samples

Leica SP8 Confocal/Multiphoton Upright Microscope having 20X water immersion lens (of 1.0 NA) and Leica Application Suite (LAS) software with FRAP wizard was used to create a photobleach pattern on the fluorescently labelled fascicles using a non-polarized 20 mW laser beam emitted from the resonator passed through focusing optics. Considering the sample curvature, tissue heterogeneity, fluorescent intensity variation due to sample depth, etc., an optimal z-position based on higher fluorescent intensity as well as maximum area on the sample (at a depth of ∼half diameter) was selected for creating a photobleach pattern with following settings: line frequency = 200 Hz; exposure time = 5.24 sec per image; intensity of laser for photobleach/non-photobleach regions = 90/0.1%; no. of iterations = 180. For an image of size 553 *μ*m x 553 *μ*m, the photobleached region results a total area of ∼7500 *μ*m^2^ (ten ∼5 *μ*m wide stripes for a 150 *μ*m diameter fascicle = 5×10×150) that covers approximately 2.45% of total area of the image where the laser stays at high intensity for 0.128 sec (2.45% of 5.24 sec) during each iteration. The total fluence in the photobleached region, computed as laser power × exposure time/square area i.e. 0.307 x 10^-7^ J/mm^2^ for each iteration, for 180 iterations as 55.29 J/mm^2^ seems a reasonable choice towards good photobleached pattern contrast and minimum sample damage due to laser exposure [102].

#### Strain quantification by Gaussian fitting

In order to make the local displacement measurements between each pair of consecutive photobleach-stripes on the sample in undeformed and deformed configurations, we fitted Gaussian curves to the ROI path (i.e., yellow line covering two-consecutive stripes (**Fig.S8B-i**) [103]. Here, a single composite image was obtained by summing up fluorescent intensities of all images within a stack covering fascicle dimensions diametrically. Using ImageJ package, a ROI path on the composite image was created to capture fluorescent intensity magnitude as function of length parallel to fascicle’s longitudinal-axis. The fluorescent intensity was transversely averaged (parallel to the stripe-length) over certain distances considering the parallelism between two neighboring stripes. We found a fixed ROI length (∼25 *μ*m) optimal for Gaussian curve fitting optimized considering the sample heterogeneity (**Fig.S8D**). For fitting a Gaussian curve to each valley of a ROI path carrying several photobleach-stripes, a custom-built MATLAB script was used. Based on the mentioned criteria, Gaussian peaks were used to calculate the displacement between each pair of consecutive photobleach-stripes in undeformed and deformed configurations (**Fig.S8B,C**). The displacement values between each pair of consecutive photobleach-stripes were then used to calculate the average value within that region between them (**Fig.S8E**).

### S4. SHG signal measurement in tissues

The SHG signal from different tissues measured at microscale represents an ensemble of nanoscale diameter collagen fibrils. In tendon tissues, collagen fibers (comprising of collagen fibrils) are mostly aligned along the longitudinal axis that result SHG intensity dependence on the angular orientation between the fascicle-axis and the direction of polarization of incident light (**Fig.1D** and **Fig.S18A**) [15]. Given that smaller diameter collagen fibrils (∼λ_SHG_/10) emit more backward SHG signal while the larger ones emit forward SHG signal predominantly [100], SHG imaging facilitates collagen fibril size based comparison within a tissue. During degradation by collagenase molecules, the insoluble collagen fibrils within tendon tissue are converted into soluble peptide fragments incapable of producing SHG signal (**Fig.S9**). For comparison of the absolute SHG intensities within a tendon tissue, the obtained SHG signal values were scaled based on the angular orientation collagen fibers with respect to incident light polarization direction and, volumetric average of SHG intensity over ∼25 *µ*m depth was considered for each tendon region (**Fig.S18A-F**, **supp.-methods S4**). For SHG intensity comparison of among tissues (including a tendon fascicle maintained at fixed orientation), ∼150 *µ*m deep image stacks (considering “non-flat” surface) with 2 *µ*m step size were captured (at minimum three randomly selected locations) and, volumetric average of SHG intensity over ∼25 *µ*m depth was considered for each tissue region (similar to **Fig.S18D-F**). For comparison of fibrillar collagen degradation rate among different regions of a tendon tissue following collagenase addition, a projected sum composite image from SHG image stack(s) covering tendon fascicle diametrically (within the photobleached region) was obtained in forward and backward directions, and the slope of SHG intensity (normalized to *t* = 0 min) versus time was used (**Fig.S9**). Such normalization procedure excluded the influence of tissue heterogeneity (resulting from troughs in tendon crimps or variation among different tendon samples) on degradation rate measurements. Temporal changes in E4-chick heart fibrillar collagen levels following molecular perturbations (myosin inhibition or collagenase treatment) were quantified based on forward SHG signal.

### S5. Comparison of MMP-1 and BC activity on soluble collagen-I

The comparison of MMP-1 and BC activity on collagen-I was made using fluorescence-based assay where normalized intensity change (with respect to negative control) was used for the degradation rate quantification. The normalized intensity was obtained using the following.

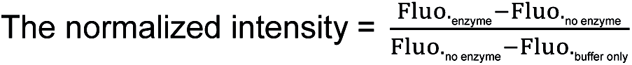

The degradation rate was measured using the following.

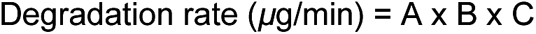

where, A = slope of normalized intensity change (with respect to negative control) versus time (min); B = Final collagen concentration (*μ*g/mL) in a reaction well; C = Volume of solution in the well (mL). BC molecules cleaves soluble collagen molecules >10 times faster than MMP-1 (**Fig.S10A**). The observation of catalytic activity of MMP-1 and BC (**Fig.S10B**) on soluble collagen suggests that the catalytic activity of MMP-1 is sensitive to presence of BSA molecules (0.01% solution) while BC activity remains almost unaffected by BSA molecules. Considering the sensitivity of MMP-1 activity and minimize reaction buffer composition changes due to evaporation from the mechano-bioreactor edges, the petri-dish carrying the fascicles exposed to MMP-1 was covered and moved to an incubator between imaging time points.

### S6. Quantitation of collagen fiber orientation distribution and deformation behavior

Although tendon fascicles carry bundles of parallel collagen fibers, there are native crimps where these fibers show an inclination to the fascicle-axis. Interestingly, we observed a collagen fiber orientation change when moving diametrically into the fascicle (**Fig.S12A-*i***). Upon mechanical deformation of the fascicle, collagen fibers tend to align along longitudinal axis (**Fig.S12A-*ii***). Local collagen fiber orientation distribution with respect to the fascicle-axis (within each pair of consecutive photobleach stripes) was calculated by means of structure tensor obtained using SHG signal for each 3 x 3 pixels group (corresponding to ∼1 x 1 *μ*m) [104, 105]. Using the collagen fiber orientation distribution within a ROI of the undeformed fascicle, respective affine predicted collagen fiber orientation distribution in the deformed configuration was estimated based on local tissue strain magnitudes (see methods). As stepwise listed below, the comparison between affine predicted and experimentally observed collagen fiber orientation distributions in a deformed ROI has been made based on quantile-quantile (Q-Q) plots (**Fig.S12B,C**) using offset and range values (**Fig.S12D**) calculated from the projection plots [51].

I. Collagen fiber orientation distribution histograms were obtained for each ROI of the fascicle using forward SHG images stack (each image taken 2 *μ*m apart) for angles taken between - 90 to +90 deg (0 deg = representing horizontal reference line)
II. The orientation of each ROI (i.e., local fascicle-axis) with respect to the horizontal reference line was determined based on projected-sum composite image the ROI stack. Straight lines on ROI edges (upper and lower) were fitted to determine the average inclination of the local fascicle-axis of the ROI.
III. Histograms representing distribution of collagen fibers with respect to the local fascicle-axis within each ROI were obtained for angles between −90 to +90 deg.
IV. The change in the displacement between the consecutive photobleach-stripes were used to represent the deformation gradient along the fascicle-axis (i.e., F_11_) for each ROI of a fascicle. Average cross-sectional area (CSA) along the local fascicle-axis of each ROI was obtained in undeformed and deformed configurations by fitting an ellipse to the fascicle cross-section. The change in radius of the fascicle cross-section was used to estimate the deformation gradient in transverse direction (i.e., F_22_). F_21_ and F_12_ were assumed as negligible (set to zero) because mechanical loading was along the fascicle-axis and parallel photobleach-stripes were considered for local strain estimations within a tissue.
V. Orientation distributions of each ROI in undeformed and deformed configuration were grouped into 100 quantiles (with 1% increment) using a custom MATLAB script. The angular values were approximated with linear interpolation if required.
VI. Quantile grouped data for the undeformed configuration was used to obtain affine predicted orientation distributions using respective deformation gradient matrix.
VII. Experimental and affine predicted orientation distributions were compared using Q-Q plots. Experimental and affine predicted values (100 groups each) were used to obtain projection plots where quantile difference (i.e., Exp. - Affine) were plotted along ordinate axis and the average of two (i.e., (Exp.+Affine)/2) was plotted as abscissa.
VIII. Offset and range were estimated to quantify deviation between experimental and affine predicted distributions in each ROI (**Fig.S12D**). Offset value represents the deviation (in degrees) of the median of quantile differences from 0 whereas the range values represent +/-34.1% spread about the median.
IX. Offset and Range values across all ROIs were compared.

To obtain the primary collagen fiber orientation with respect to the fascicle-axis within different regions (i.e., ROIs) of an undeformed fascicle, obtained collagen fiber orientation distributions were grouped into 100-quantiles (for angles with respect to fascicle-axis). Based on the extreme values of obtained quantile angles, collagen fiber distribution histograms were obtained for each class-size of 2.5 deg (for angles between −45 to +45 deg). Gauss-peak was used to represent the primary collagen fiber orientation while Gauss-width indicated a spread of orientation values with respect to the fascicle-axis.

### S7. Waviness value estimation

Tendon tissue waviness is calculated by the spatial averaging of local collagen fiber orientation changes when moving along the longitudinal-axis of a fascicle (**Fig.S13A**) based on the following steps

I. Local collagen fiber orientation (with respect to horizontal reference line) vector-field per slice (pixel-size = ∼1 x 1 *μ*m) was obtained to represent local orientation for a ROI.
II. Pixels lying within each slice of the tissue were selected for further investigation while considering the rest as background.
III. Based on the orientation of each ROI (local fascicle-axis orientation on projected sum composite image of forward SHG images stack) with respect to horizontal reference line, a slice-wise collagen fiber orientation with respect to the fascicle-axis and corresponding pixel coordinates (x, y) were determined.
IV. x-values and corresponding angles values were locally grouped and averaged into 30 bins (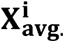 and 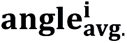, for i = 1:30) i.e., every ∼2 *μ*m along the fascicle-axis.
V. Based on 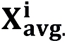 and 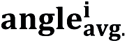 values and setting 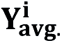 = 0 for *i* = 1 was used (assuming starting 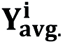 on fascicle center line). 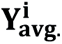 (for 1 < *i* <= 30) values were obtained using following relation to predict local waviness of slice based on local fiber orientation 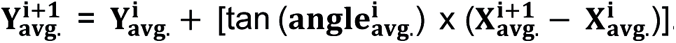.
VI. Line segments between consecutive points 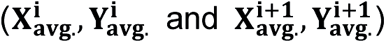 were then used to represent local length of curve and estimate total length of the curve (L_curve_).
VII. Displacement between starting and end points of the curve were then used to calculate end- to-end displacement (L_e2e_).
VIII. Waviness within each slice of a ROI was estimated based using the following relation i.e., Waviness = ((L_curve_ /L_e2e_)-1) x 100.
IX. Above steps were then used to estimate the waviness values for all slices lying within an ROI of a fascicle covering entire tissue-depth.
X. Median value of waviness for each ROI was selected to represent the final waviness of the ROI.
XI. Normalized waviness value (i.e., with respect to waviness value before fascicle deformation) for a ROI with time (following addition of BC) were estimated.

### S8. Fluorescent labelling of BC molecules at N-terminus

BC molecules were fluorescently labelled (**Fig.S21**) by mixing with NHS-Ester 555 (per supplier protocol) in two different proportions using following steps

I. Two different relative proportions of BC with respect to the fluorophore were reacted for 2 hours at 4 deg. i.e., 50 *μ*L (10 mg/mL) of fluorophore was added to 0.5 mL of 20 mg/mL (producing 4X-BC) or 1 ml of 10 mg/mL of BC (producing 1X-BC)
II. Overnight dialysis of the samples was done using 10 kDa cut-off (given mass of BC lies in 65-130 kDa range) in PBS at 4 deg C.
III. UV scanning of labelled BC samples and control samples (without labelling) were compared to verify BC labeling by fluorophore.
IV. Higher concentration (4X) of BC in reaction resulted in higher labelling efficiency.
V. Obtained concentration for 1X solution was 6.31 mg/mL whereas for 4X, it was 13.157 mg/mL (calculated based on extinction coefficient of unlabeled-BC)

HPLC was used to determine if all different masses of BC molecules present in the solution were labelled by the fluorophore (**Fig.S21B**) as BC solution represents a cocktail of at least 7 different proteases (molecular weight range ∼68–130 kDa).

I. A comparison of two different concentrations of fluorescent-BC solutions obtained by the labelling protocol was made
II. Unlabeled BC samples (1 mg/ml), Fluorescent-BC (13.1 mg/ml and 6.3 mg/ml) in PBS were stored at −20 deg. and thawed at 4 deg. for 24 hours before testing.
III. Samples were diluted in PBS to similar concentrations i.e., unlabeled-BC (1 mg/ml), fluorescent-BC (1.31mg/mL) and fluorescent-BC (0.63 mg/mL) before HPLC run
IV. Samples were run in polarity-based HPLC by a transition of mobile phase from polar to non-polar ratio of 100:0 to 60:40 (DI water: Acetonitrile) for 30 minutes
V. Signal intensity (peak height) at 280/555 was observed proportional to the sample concentration values
VI. Fluorescent-BC samples show small polar masses absorbing at 280 nm (highlighted sky-blue) resulting from the self-degradation of BC due to the presence of Alexa fluorophore at N-terminus of BC.
VII. Both fluorescent-BC samples show absorbance at 555 nm (highlighted sky-blue) for the range of masses while 555 nm absorbance is absent in the unlabeled-BC indicating all masses are fluorescently labelled

### S9. Fluorescence recovery after photobleaching (FRAP) measurements

For each experiment, the stock solutions of TRITC-dextran or fluorescent-BC in PBS were added to the medium carrying heterogeneously strained tendon fascicle and kept overnight at 37 deg C before making FRAP measurements. All measurements were performed on Multiphoton microscope TCS SP8 MP (Leica) in FRAP mode using a 20 mW laser with a 20X /1.0 numerical aperture objective, with a 5.24 A.U. pinhole, which yielded a 7 *μ*m thick optical section. At center of image (of size 69.33 x 69.33 *μ*m), a square region (5 *μ*m x 5 *μ*m) was selected for photobleaching. Each measurement was made into deep regions of the tissue (at least than 25 *μ*m away from the surface) while the acquisition of each image (with 512 x 512 pixels taking 1.295 sec) represented a time-point. Minimum value of laser intensity was selected to avoid photobleaching during the image acquisition while the intensity was set to 90% of maximum laser power for the photobleach sessions. All FRAP experiments consisted of pre-bleach, bleach and post-bleach sessions taking 303.3, 151.15 and 3367 sec respectively (using bi-directional scan). Pre-bleach images were used for quantifying photobleaching during image acquisition at 0.01% laser intensity and making necessary corrections/normalization to the obtained data. The compensation evaporation losses from reaction buffer during imaging was made every ∼2 hours by addition DI water (**Fig.S6**).

For extracting mobility characteristics based on FRAP experiments, a single exponential curve was fitted to the normalized recovery data. To avoid fluctuations in the data for curve fitting purposes, the obtained ROI intensity values were binned over constant intervals. Double normalization of the data was performed for the intensity values obtained from photobleached regions with respect to the average background intensity as well as photobleaching during image acquisition [106]. Following double normalization, the data was scaled between 0 to 1 with respect to the maximum value during pre-bleach session and minimum values at end of beach session. For normalization protocol, we used the intensity values within three different regions i.e., bleached spot, background intensity and all image, at various time points of pre-bleach, bleach and post-bleach sessions. We used following relation to calculate the normalized intensity in the bleached region for all data sets

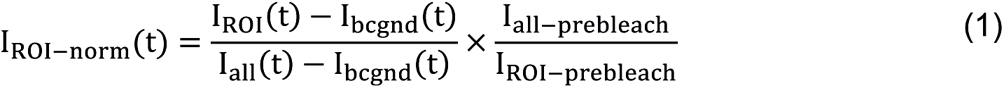

where, I_ROI−norm_(t) is normalized intensity of bleached region at different time points, I_ROI_(t) average intensity from bleached region, I_bcgnd_(t) average background intensity, I_all_ (t) average intensity of the full image, I_all−prebleach_ average intensity of all images averaged over all time points in pre-bleach session and I_ROI−prebleach_ as average intensity of bleached region averaged over all time points in pre-bleach session. Recovery characteristics of fluorescent molecules after photobleaching are obtained by using the following relations where amplitude(s) and time constant(s) of the curve represent mobile fraction(s) and time taken for recovery respectively

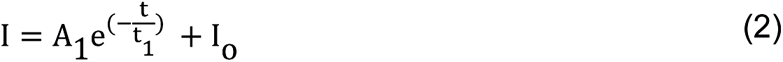

For single exponential curve fitting to recovery data, the mobile fraction value is given by

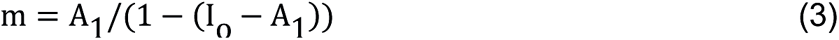

### S10. Probe synthesis and purification

#### Solid-Phase Synthesis

General probe synthesis procedures were adapted from those of Mason Smith based on his previous work in peptide synthesis [107]. All probes were synthesized on solid phase using Fmoc chemistry. Low-loading rink amide MBHA resin was swelled for 10 min in DCM followed by 3 min in DMF. N-terminal Fmoc group was removed by stirring in 20% piperidine/DMF for 5 min, after which solution was replaced and stirred for an additional 15 min. Resin was then washed 5 times with DMF. Coupling solutions were prepared by combining reagents and DMF solvent in conical vials, mixing, and allowing to activate briefly at room temperature (≥5 min). Each coupling solution contained: 5 eq. Fmoc-protected building block, 5 eq. COMU, and 10 eq. DIEA, with all equivalents determined relative to the resin. Activated coupling solutions were added to deprotected resin and stirred for ≥45 min. Resin was then washed again with DMF (5x). Following coupling of PEG linker during synthesis of probe-1, resin was washed an additional 5 times with DCM. When stopping synthesis at any point, resin was washed an additional 5 times with DCM and dried by vacuum; when resuming synthesis, resin was swelled as described above. These steps of Fmoc deprotection, washing, coupling, and washing were repeated for coupling of each Fmoc-protected building block. Final fluorophore labeling reaction was adapted from established protocols [108]. Briefly, 6 eq. 5(6)-CTAMRA, 6 eq. COMU, and 12 eq. DIEA were combined in DMF and allowed to activate as described above. Activated coupling solution was then added to deprotected resin and stirred for 3 days. For synthesis of probe-1, resin was subsequently washed with DMF (5x) then stirred in 20% piperidine/DMF for 30 min.

#### Final Cleavage & HPLC Purification

After completing synthesis, each probe was purified as follows: Resin was washed with DMF (5x), followed by DCM (5x), and then dried by vacuum. Probe was cleaved from resin and tBu protecting groups were cleaved from hydroxyproline (Hyp) residues by stirring in TFA:TIPS:H_2_O (95:2.5:2.5) for 1.5-2 hours. Crude probe solution was then filtered through frit in reaction vessel into pear-shaped flask. Resin was then rinsed twice with TFA, and rinses were added to flask. Solvent was evaporated in vacuo, then product was resuspended in 10% citric acid or 1N HCl. Organic fraction was extracted ≥3 times with DCM. Remaining aqueous fraction was concentrated to minimal volume in vacuo, resuspended in 50:50 H_2_O:ACN, 0.22 *μ*m filtered, and purified by HPLC in gradient of ACN/H_2_O (0.1% TFA) at 80 °C (Due to low yield for synthesis of 6-CTAMRA-PEG control probe-2, it was subsequently determined that organic fraction contained large amount of product. For this probe, organic fraction was dried in vacuo, resuspended, filtered, and purified as described above, then combined with purified product from aqueous fraction). Temperature of column/solvent was elevated to prevent oligomerization of probes during separations. Because labeling of probes with 5(6)-CTAMRA produced two isomers (i.e., probes labeled with 5-CTAMRA or 6-CTAMRA), isomers were separated by HPLC during purification and 6-CTAMRA labeled probes were used for subsequent assays (“Validation Data” below).

#### Preparation of Peptide-Loaded Resins

To streamline general SPPS of CMP-based fluorescent probes, rink amide resin was loaded with peptide sequence shared by multiple probes (i.e., Fmoc-[azGPO(tBu)]_3_-resin). Loaded resin was thoroughly washed and dried, after which small peptide aliquot was cleaved from resin in order to check purity and mass by analytical HPLC and MALDI-TOF MS, respectively. Resin loading density was determined using published UV-vis spectroscopy-based assay. Loaded resin was then stored under desiccation and used as starting material for multiple peptides used in this and other studies, including probe-1.

#### Probe Stock Preparation

Stock concentrations were determined by UV-vis spectroscopy. To prepare samples for UV-vis measurement, concentrated aqueous solutions of probes were diluted and added to 1-cm quartz cuvettes. 4-5 dilutions were prepared from each stock for replicate measurements. UV-vis scans were collected in continuous scan mode with a 200 nm/min scan speed, 1.0 nm data interval, and 1.0 nm bandwidth. Because it has been observed that fluorophores conjugated to N-termini of trimeric collagen peptides can self-quench when peptides assemble into triple helices [109]. The measurements were recorded at 65 °C to inhibit self-assembly. Absorbance of N-terminal CTAMRA fluorophore in each dilution was measured at 555 nm and averaged, after which probe concentrations were calculated using average A_555_ and molar extinction coefficient of 89,000 M^-1^ cm^-1^ [110]. Concentrated probe stocks were then aliquoted and dried by vacuum centrifuge for storage prior to use, at which time they were resuspended as needed to achieve necessary concentration for subsequent assays.

#### Synthesis and Purification of Fmoc-azGPO(tBu)-OH, and validation data

To streamline Aza-peptide probe synthesis, protected tripeptide Fmoc-azGPO(tBu)-OH was synthesized and used as principal building block for SPPS. General protocol outlined below was developed using standard principles of Fmoc SPPS on 2-chlorotrityl chloride resin; one notable reference is included here [111].

#### Initial Resin Loading with Hydroxyproline

To a 500-mL SPPS vessel, Fmoc-Hyp(tBu)-OH (30.71 g, 75 mmol, 1.5 eq.) and DCM (200 mL) were added and stirred briefly. To this mixture, 2-chlorotrityl chloride resin (34.25 g, 50 mmol, 1.46 mmol/g) and DIEA (26.1 mL, 150 mmol, 3 eq.) were added. Reaction mixture was stirred for 5 hours. After 5 hours, MeOH (70 mL, ∼2 mL/g resin) was added to cap remaining reactive trityl groups, and resin was stirred for 1 hour. Resin was washed with DMF (5x).

#### Proline coupling

N-terminal Fmoc group was removed by stirring resin in ∼200 mL 20% piperidine/DMF (2 x 30 min). Resin was washed with DMF (5x). Fmoc-Pro-OH (50.60 g, 150 mmol, 3 eq.), HBTU (56.89 g, 150 mmol, 3 eq.), DIEA (52.3 mL, 300 mmol, 6 eq.), and DMF (300 mL) were combined in a 1000-mL round bottom flask and stirred to activate for several minutes. Activated coupling solution was added to resin and stirred overnight (15.5 hours). Resin was washed with DMF (5x).

#### Aza-glycine coupling

Deprotection and wash steps were carried out as described above. CDT (24.62 g, 150 mmol, 3 eq.), Fmoc-carbazate (38.14 g, 150 mmol, 3 eq.), and DMF (350 mL) were combined in a 1000-mL round bottom flask and stirred to activate for several minutes. Activated coupling solution was added to resin and stirred for 2 days (45.5 hours). Resin was washed with DCM (3x) followed by MeOH (3x) and dried by vacuum.

#### Cleavage & Drying

Dry resin was transferred to a 2000-mL round bottom flask. Protected tripeptide was cleaved from resin by stirring in a 1-L cleavage cocktail of 30.0% TFE, 0.3% TFA in DCM for 4 hours. Reaction mixture was filtered through filter paper into separate 2000-mL round bottom flask. Resin was rinsed several times with DCM and rinses were filtered into crude peptide solution. Solvent was evaporated in vacuo.

#### Purification

Crude product was purified by flash chromatography (ISCO) in gradient of 70-100% EtOAc/Hex (EtOAc containing 0.1% TFA). Fractions containing product were pooled and dried in vacuo. Purified product was azeotroped 3 times with ∼300 mL toluene. Final product was dried on hi-vac and aliquoted as needed.

#### Validation Data

HPLC gradients and conditions are as noted. CHCA was used as matrix for all MALDI-TOF MS. For each probe, HPLC and MALDI-TOF MS data for corresponding 5-CTAMRA-labeled isomers (removed during purification) are shown alongside data for purified 6-CTAMRA products used in assays. See **Fig.S27,S28** for validation data, and **Table-S4** for abbreviations.

---

**Fig.S1:.**
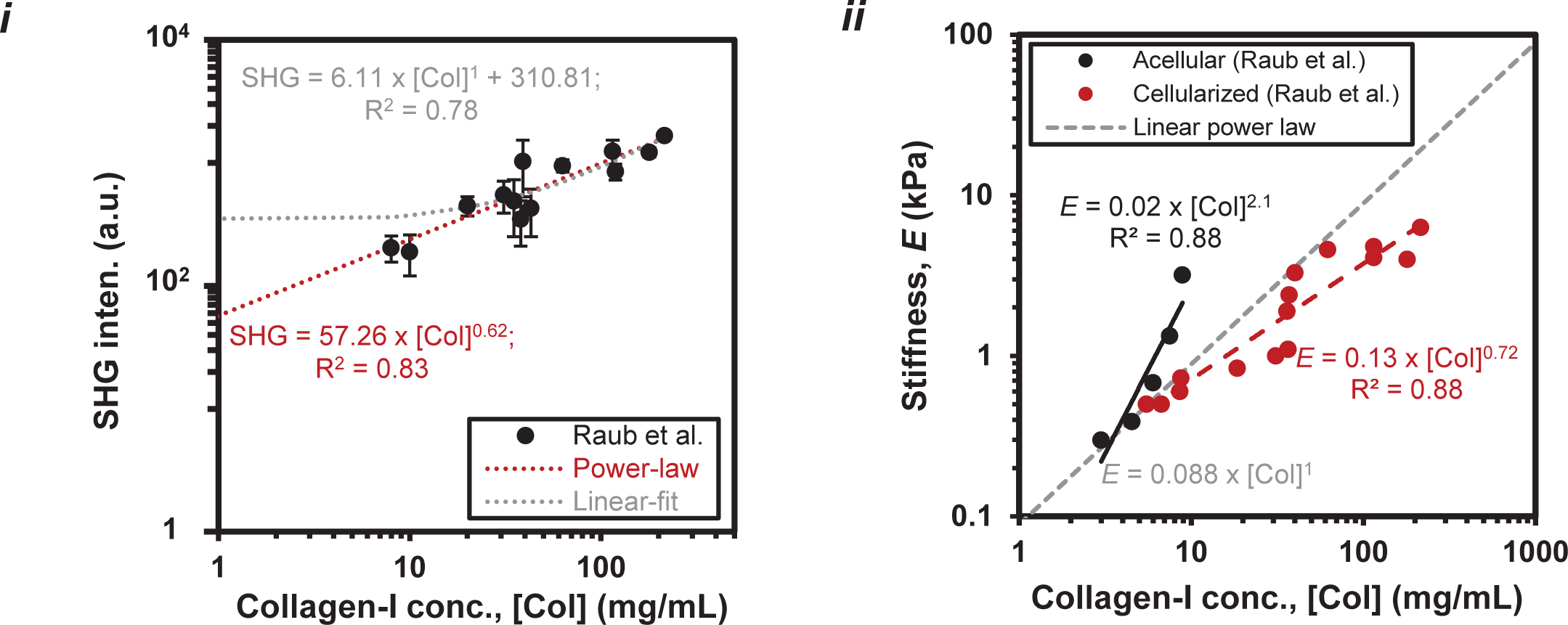
SHG intensity and stiffness scale with concentration of purified rat tail collagen-I in gels. (i) Backward SHG intensity versus collagen concentration is better represented by non-linear power-law (SHG ∼ [Col]^0.62^) as compared to a linear-fit with non-zero intercept (adopted from Raub et al., 2010) as shown by respective R-square values of 0.83 and 0.78, and its extrapolation to zero concentration. (ii) Stiffness versus collagen concentration (adopted from Raub et al., 2010) also follow non-linear power-law scaling (*E_gel_* ∼ [Col]^a^, a = scaling exponent) in case of acellular and cellularized (with normal human lung fibroblasts (NHLFs)) collagen gels where cellularity decreases the scaling exponent from 2.1 to 0.72. For cellularized gels, Raub et al. estimated major effects of gel contraction after weeks, observed some degradation of labeled collagen, but ECM synthesis by the cells is unclear

**Fig.S2:**
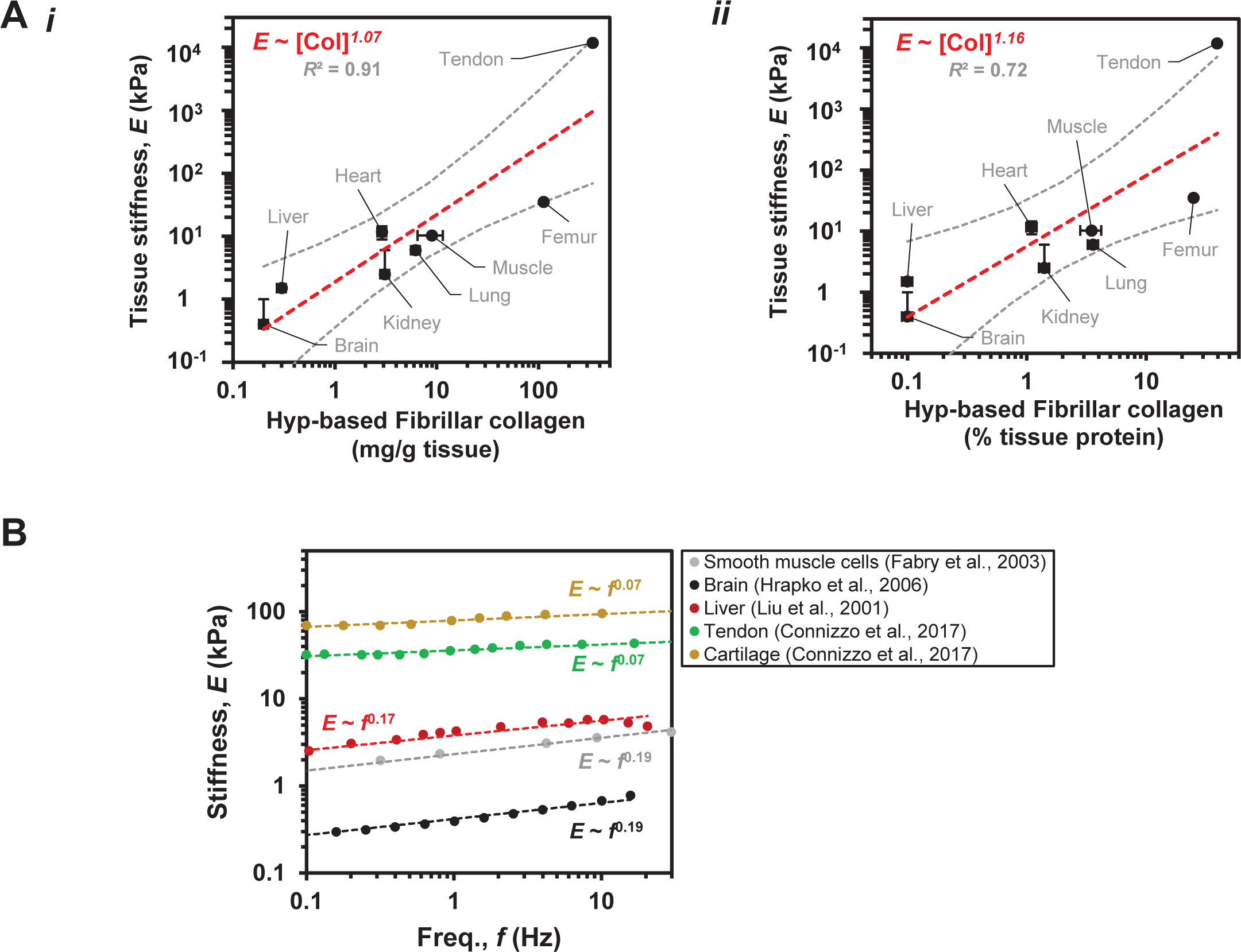
**A.** Tissue stiffness scales with hydroxyproline (Hyp)-based fibrillar collagen level in tissues with fibrillar collagen levels defined as mg/g of wet weight or as percentage total protein in tissue (per Tarnutzer et al., 2023 PMID: 36934197). **B.** At low strain rates, tissue stiffness scales weakly with strain rate.

**Fig.S3.**
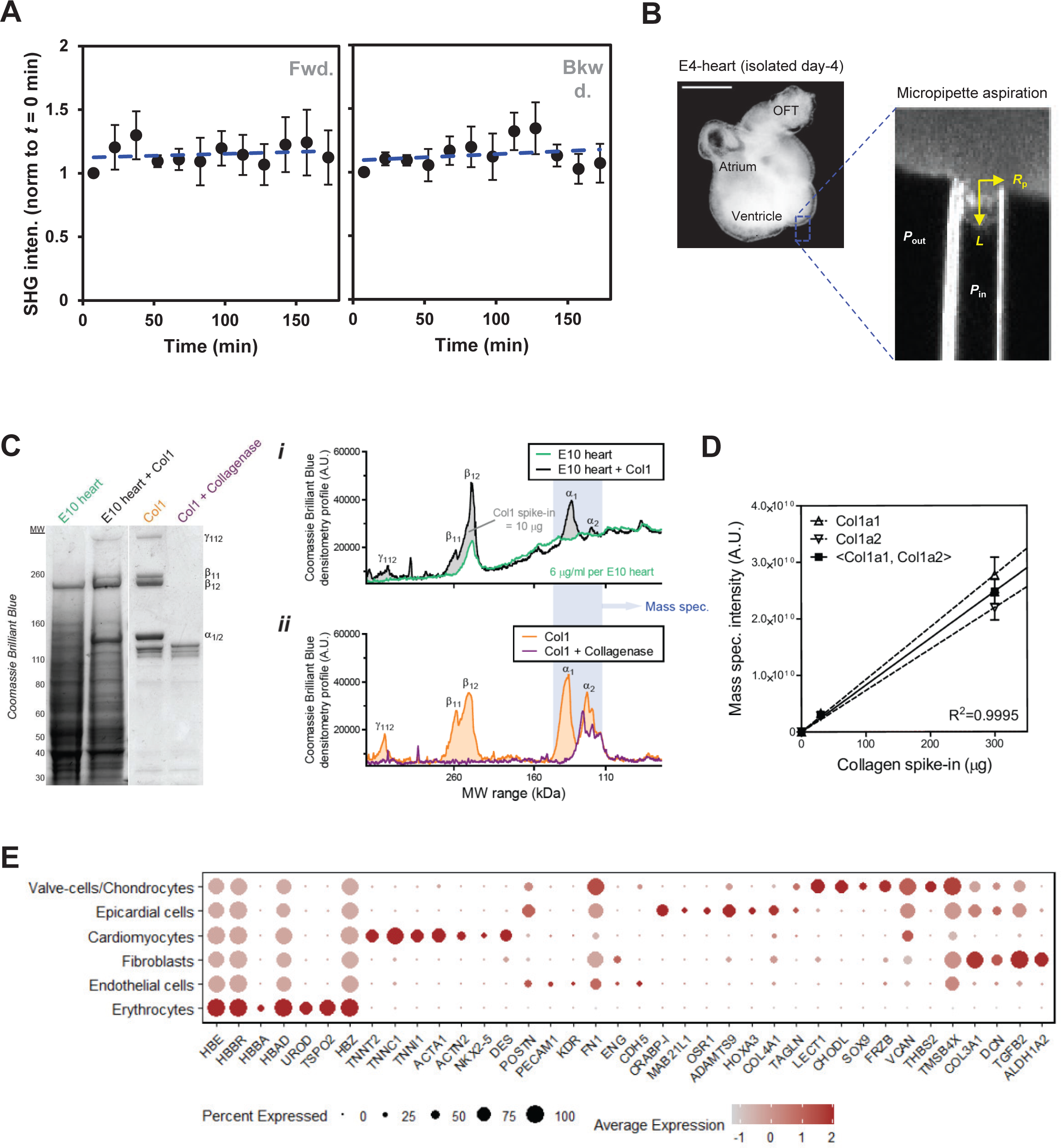
Quantitation of collagen levels in isolated embryonic chick hearts based on SHG signal and mass-spectrometry (MS) measurements. **A.** Temporal measurements of SHG signal (normalized w.r.t. time, *t* = 0 min) show that collagen levels within beating embryonic day-4 (E4) chick hearts remain unaltered in the presence of endogenous collagenases during the experimental duration of 3 hours. Absolute SHG signal was stronger in forward direction than backward direction and, thus forward SHG signal was selected for data analysis (n ≥ 4 hearts; All error bars indicate ±SEM). **B.** (i) Image of an E4 chick heart. ‘OFT’: Outflow tract, ‘Atr’: Atrium, ‘Ventr’: Ventricle (scale bar = 500 *µ*m). Right inset: Tissue stiffness (*E_t_*) measured by micropipette aspiration. **C-D.** Analysis of collagen-I present in embryonic chick hearts. **(C)** Coomassie Brilliant Blue gel image of E10 heart lysates, E10 hearts with spike-ins of purified rat-tail collagen-I, purified collagen-I only, and collagen-I with BC treatment (n = 2 hearts per condition). (i-ii) Densitometry profiles show collagenase degrades collagen-I and removes all trimer (γ), dimer (β), monomer (α) bands. **(D)** MS analysis of gel bands (100–140 kDa) spiked in with known amounts of purified collagen-I provide calibration curves that can be used to approximate total collagen content (in *µ*g’s) in embryonic hearts (Error bars indicate ±SEM). E. Gene expression of cell-type specific markers. Dot-size encodes the percentage of cells within a cluster, while the color encodes the average expression level (log_e_ normalized) across all cells within a cluster (red is high).

**Fig.S4.**
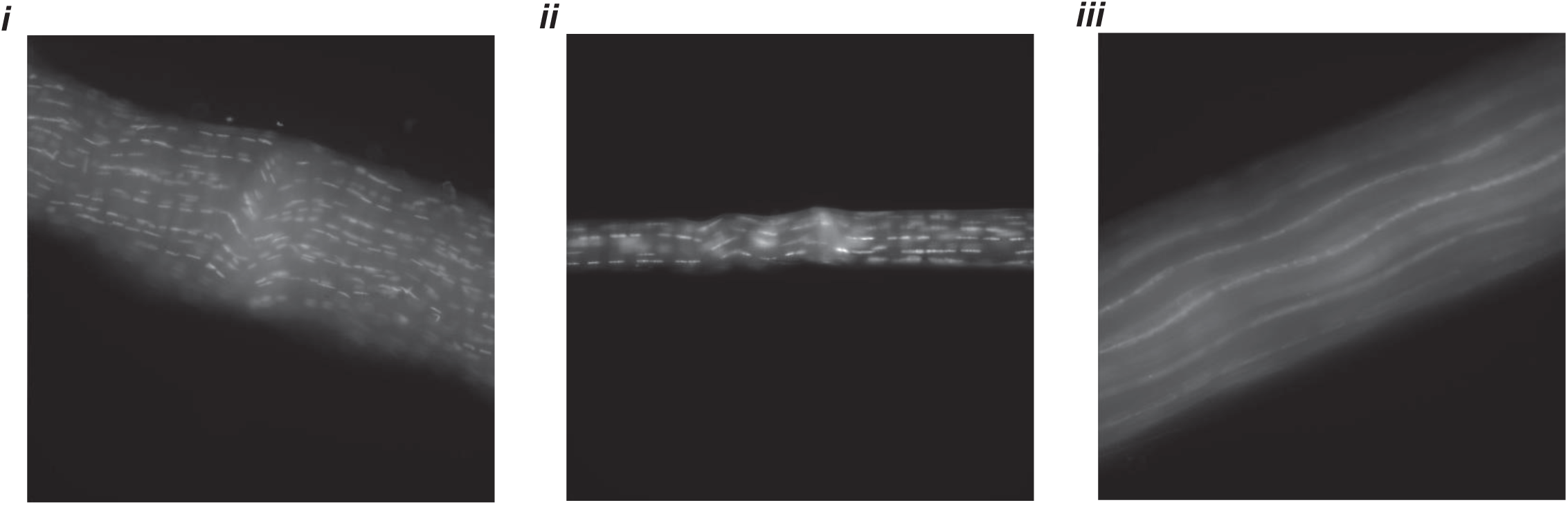
Freeze-thaw cycles produce a “cell or protein-synthesis free” ECM of tendon fascicles. Localized chromatin distribution based on Hoechst staining in case of tendon fascicles (i-ii) from C57BL/6J mice tail obtained immediately after euthanizing and, form NSG mice tail after euthanizing and paraformaldehyde fixing together suggest non-ruptured cells. (iii) Starved tendon fascicles from C57BL/6J mice after several freeze thaw cycles and long-term storage (∼few months) at −20°C results cell death as shown by chromatin distribution along the fascicle length. Fascicle diameters up to 200 *µ*m.

**Fig.S5.**
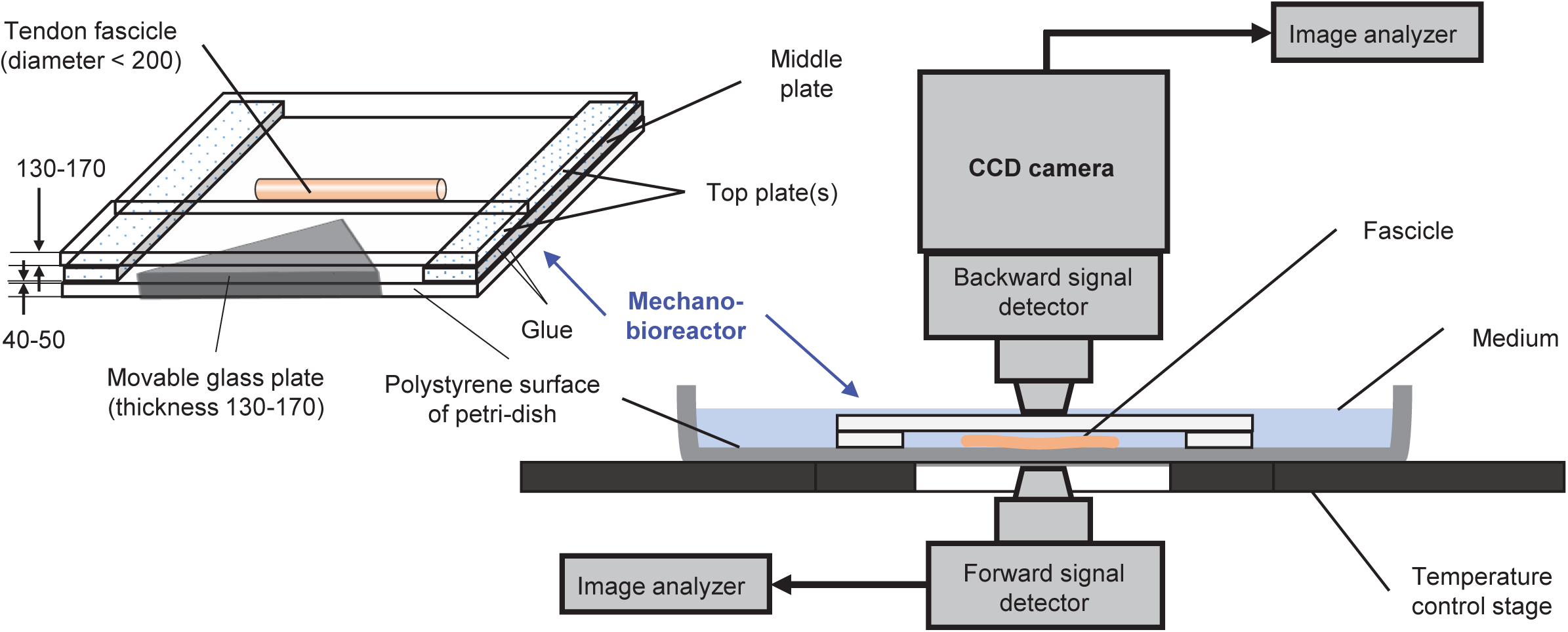
Schematic of the mechano-bioreactor for heterogeneous deformation of a fascicle via in-plane three-point bending and, its installation on a MP-microscope for SHG imaging in both forward and backward directions. All dimensions are in microns.

**Fig.S6.**
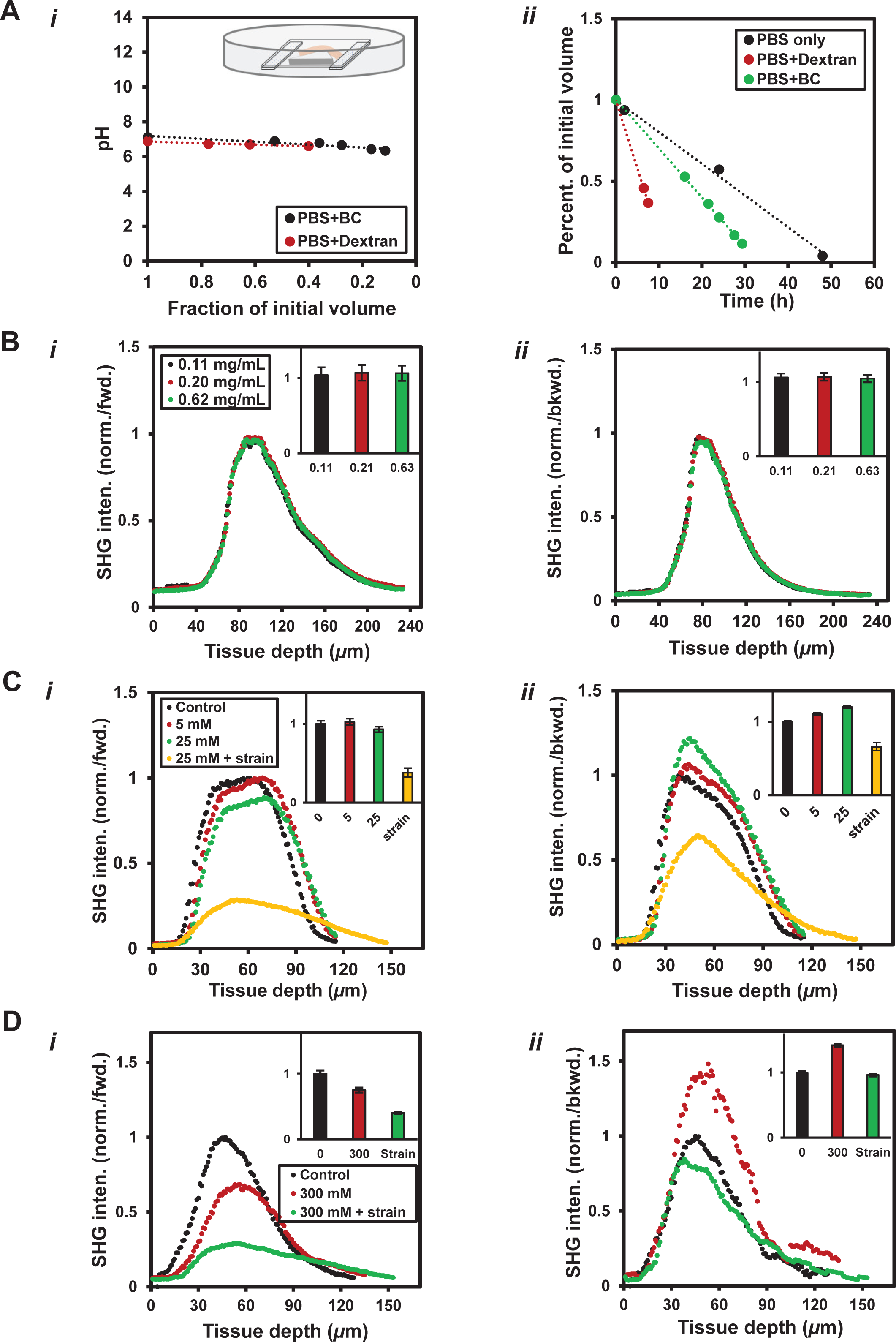
Reaction buffer composition and its effect on packing of collagen fibrils within tissues. **A.** (i-ii) Although the evaporation of the reaction buffer from the edges of mechano-bioreactor (kept at 37 °C; initial volume = 5 mL) does not affect its pH, the higher evaporation rate of reaction buffer (i.e., PBS) carrying TRITC-dextran as solute than that of BC suggests need for compensation against evaporation losses. **B.** (i-ii) Dextran concentration changes in the reaction buffer do not affect the packing of larger and smaller size collagen fibrils within fascicles as shown by SHG signal values in forward and backward directions respectively. **C.** (i-ii) SHG signal values in forward and backward directions show that the increase in CaCl_2_ concentration promotes disassembly of larger diameter collagen fibrils into the smaller diameter fibrils. The superimposition of mechanical strain on a fascicle in addition to the presence of CaCl_2_ promotes disassembly of all size collagen fibrils. CaCl_2_ concentrations: Physiological = 1.1–1.4 mM, Fluctuation range = 0.5–80 mM (in ion channels). **D.** (i-ii) SHG signal in forward and backward direction in a fascicle shows that the increase in NaCl concentration promotes disassembly of larger diameter fibrils. The superimposition of mechanical strain on a fascicle in presence of NaCl results disassembly of all size collagen fibrils. NaCl concentration: Physiological = 120 mM.

**Fig.S7:**
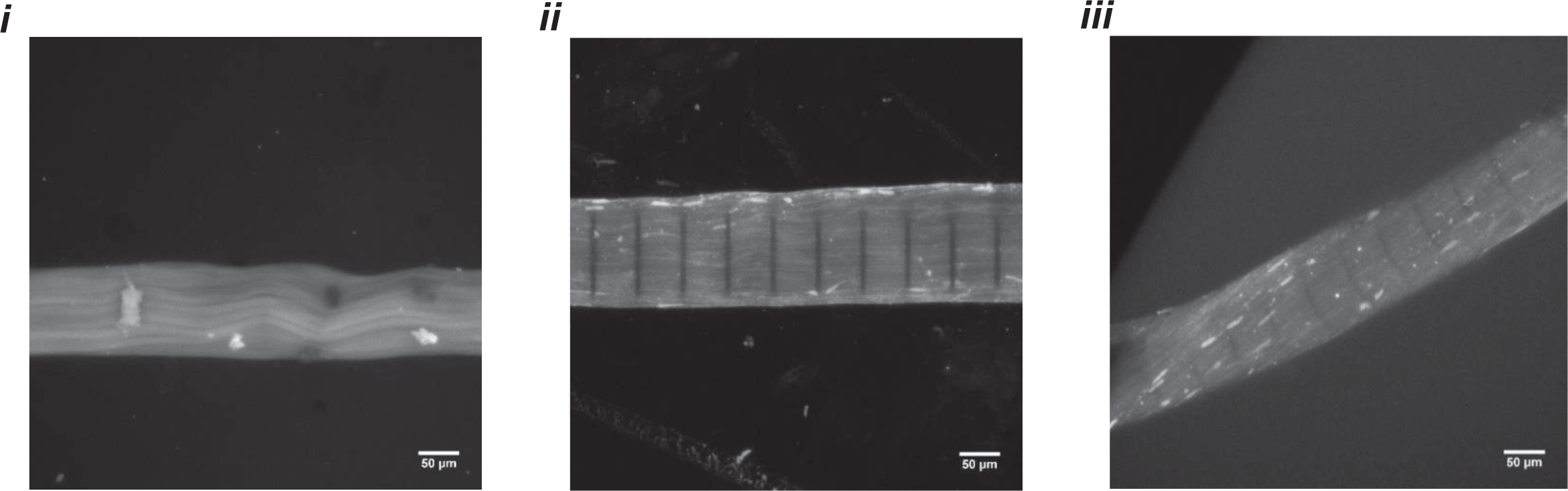
The novel Aza-peptide shows affinity towards mechanically unstrained and strained tissue ECM. (i) Mechanically unstrained tendon fascicles labelled with fluorescent Aza-peptide. (ii) Unstrained fascicle after labelling with fluorescent Aza-peptide and pattern photobleaching. (iii) Mechanically strained fascicle following labelling with fluorescent Aza-peptide.

**Fig.S8.**
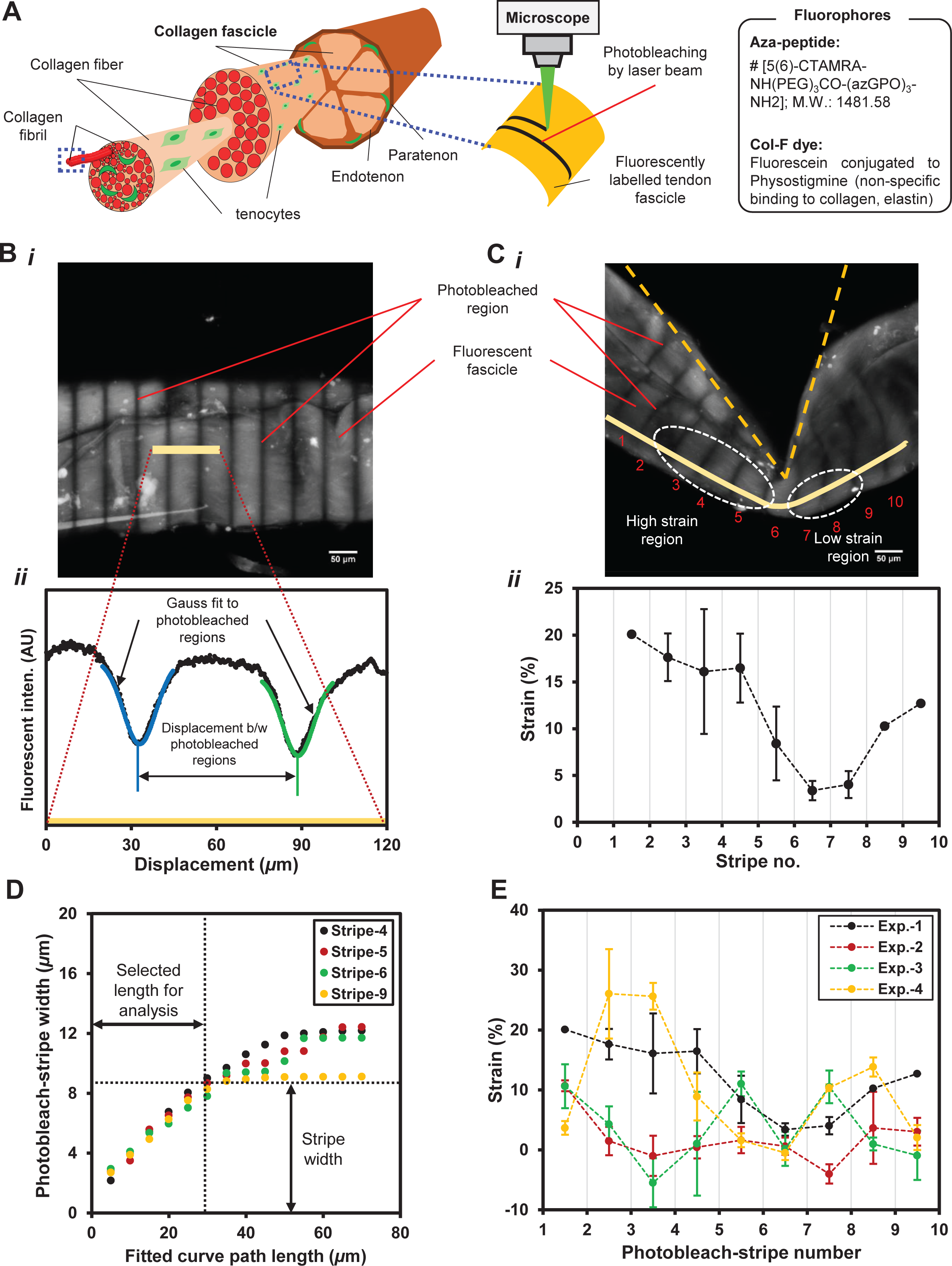
Structural hierarchy in a tendon fascicle and, strain quantification using fluorescent pattern photobleach (FPP) method. **A.** Fluorescent labelling of the fascicles by either Aza-peptide or Col-F dye for pattern photobleaching purposes. **B.** (i) A projected sum intensity image of an unstrained fluorescently labelled fascicle after pattern photobleaching, and a representative path across consecutive photobleach-stripes (i.e., light-yellow bar) selected for intensity-based measurements. (ii) The intensity profile along the representative path and curve fitting to obtain Gaussian-peak position in order to calculate the displacement between the photobleach-stripes. **C.** (i) High and low strain regions of a mechanically deformed fascicle. (ii) Strain distribution in the regions between each pair of consecutive photobleach-stripes on the fascicle along the representative path. **D.** The measured increase in Gaussian-width as a function of path length across a photobleach-stripe during curve fitting suggests a local tissue heterogeneity. A path length of 25 *μ*m for Gaussian curve fitting is selected to obtain Gaussian-peak position in order to represent photobleach-stripe position on the fascicle. **E.** The strain distributions among the fascicles differ despite their deformation in three-point bending mode and suggest a material heterogeneity.

**Fig.S9.**
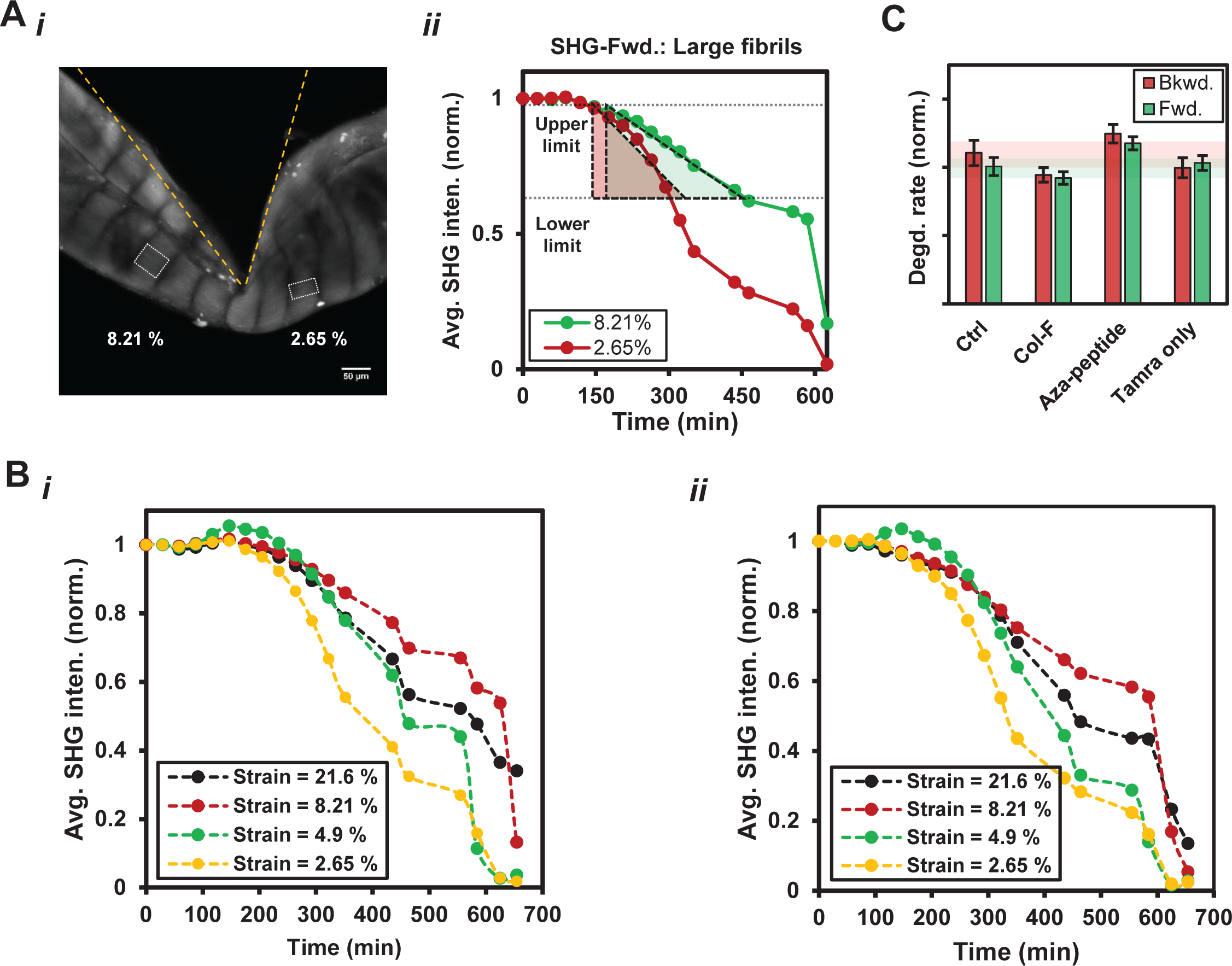
SHG signal based collagen degradation rate measurements in different regions of a tendon fascicle and, the influence of fluorescent labelling of the fascicle on the measured degradation rate. **A.**(i-ii) Representative areas in high and low strain regions of a fascicle over which SHG signal was spatially averaged. Average SHG (forward) signal from these regions decreases at different rates when the rate of each strained region is represented by the slope of the normalized SHG intensity (w.r.t. *t* = 0 min) versus time. The selection of initial SHG intensity change (from 0.95 to 0.60) minimizes influences of strain relaxation or collagenase/substrate concentrations changes on the measurements. **B.** Representative temporal changes in average SHG signal intensity within different regions of a strained fascicle that is subsequently exposed to BC in (i) Backward direction (ii) Forward direction. **C.** The degradation rates of the fascicles labelled with different fluorophores i.e., Col-F dye, Aza-peptide, TAMRA-only (fluorophore with peg-linker) or control (i.e., unlabelled) under exposure to BC were not significantly different. Aza-peptide with TAMRA fluorophore showed more affinity to the fascicles than TAMRA-only as indicated by their absolute intensities (n ≥ 3 fascicles per condition; All error bars indicate ±SEM).

**Fig.S10.**
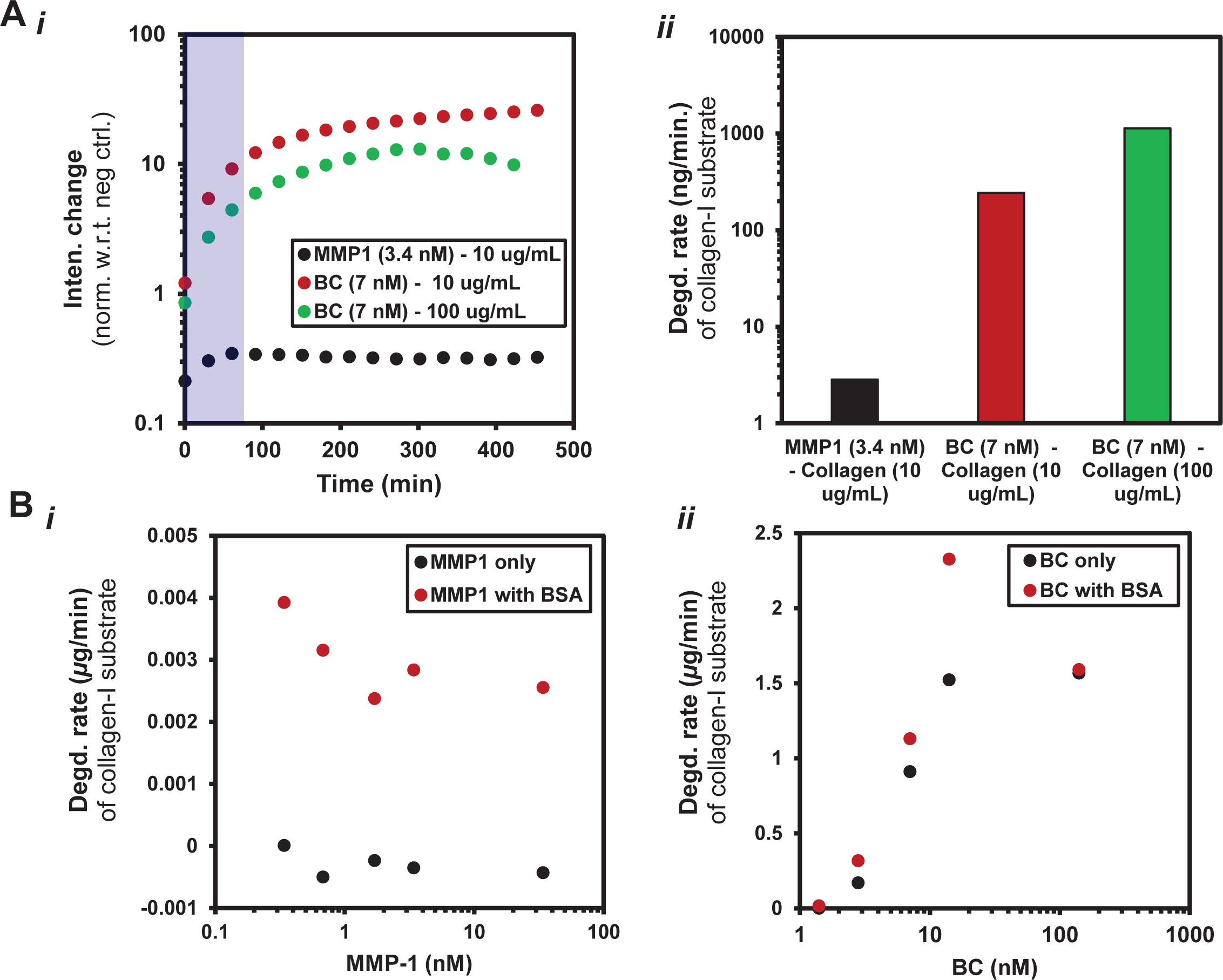
The comparison of cleavage activity of MMP-1 and BC on monomeric collagen-I fluorescein conjugate. **A.** (i-ii) BC molecules cleave monomeric DQ^TM^ collagen-I (from bovine skin) >10-fold faster than MMP-1 as indicated by fluorescence-based assay. **B.** (i-ii) MMP-1 activity on monomeric DQ^TM^ collagen-I is influenced by the presence of bovine serum albumin (BSA) (0.01% solution) whereas the activity of BC is unaffected by the presence of BSA.

**Fig.S11.**
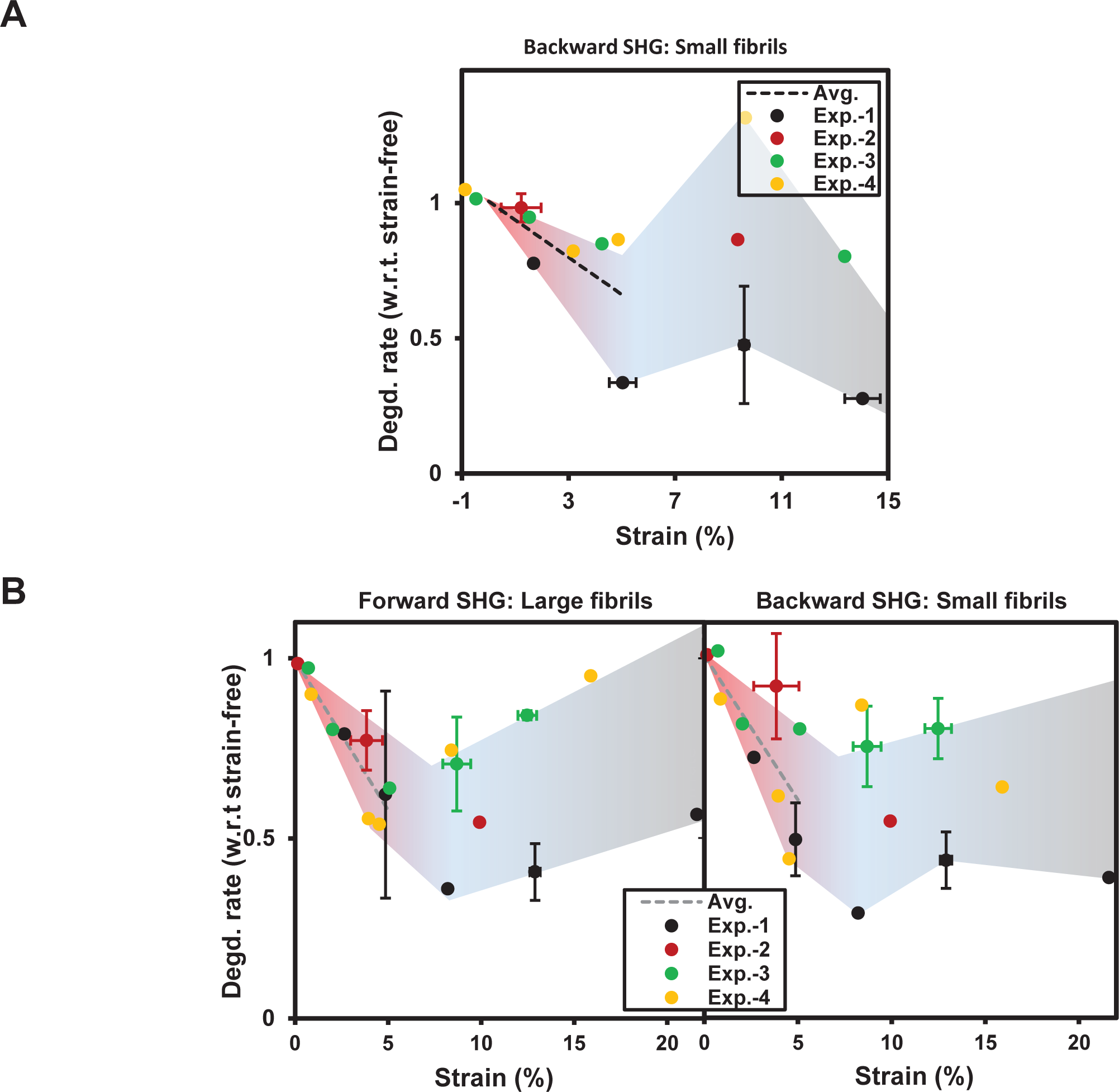
Heterogeneous strains suppress collagen degradation rate in the fascicles exposed to different collagenases when strains are kept within physiological limits. **A.** Backward SHG signal based collagen degradation rate measurements of the heterogeneously deformed fascicles exposed to purified MMP-1 shows tight suppression when strain values are up to ∼8%. **B.** (i-ii) SHG signal based collagen degradation rate of the heterogeneously deformed fascicles exposed to BC shows a similar suppression when strain values are up to ∼8%. SHG signal in forward and backward direction is produced by large and small size collagen fibrils respectively (n = 4 fascicles per MMP1 or BC; All error bars indicate ±SEM).

**Fig.S12.**
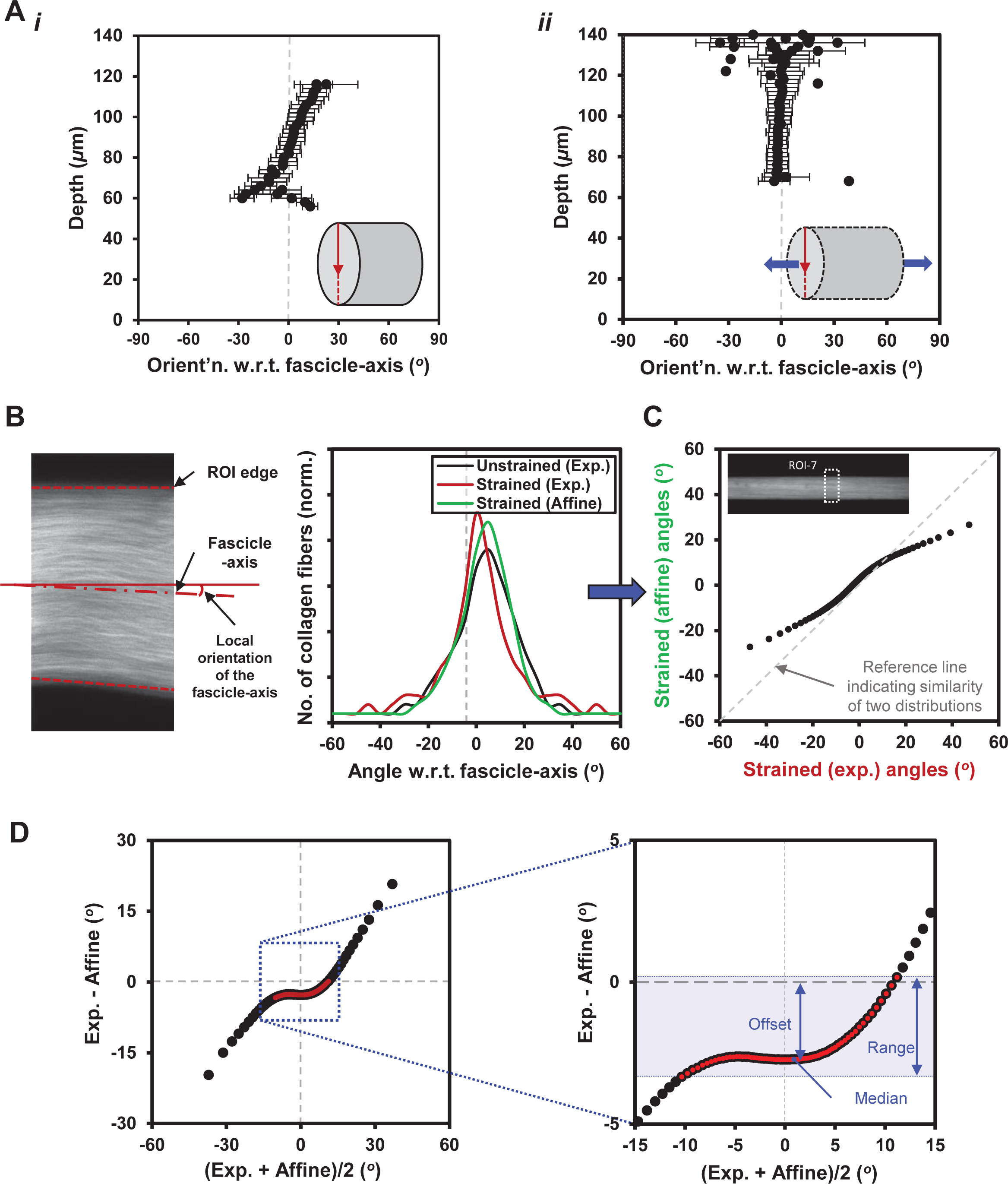
Analysis of deformation behavior of tendon fascicles using projection plots. **A.** (i-ii) Primary collagen fiber orientation (w.r.t. the fascicle-axis) within a region of the fascicle as a function of tissue depth (diametrical direction in the fascicle as indicated by red-arrow) shows that the inclined collagen fibers in the undeformed configuration tend to align along the fascicle-axis upon mechanical deformation of the fascicle. **B.** Local orientation of the fascicle-axis (i.e., within a region of interest (ROI)) is calculated based on its local geometry defined using forward SHG signal. The distributions of collagen fiber orientations (w.r.t. the fascicle-axis) within a ROI of a fascicle in both experimentally observed unstrained and strained configurations and in affine-predicted strained configuration. **C.** The comparison of collagen fiber orientation distributions between experimentally observed and affine predicted strained configurations within a ROI by means of quantile-quantile (Q-Q) plot. **D.** Projection plot for quantitative comparison of angular distributions of collagen fibers in experimentally-observed strained and affine-predicted strained configurations of a ROI. Abscissa represents mean of experimentally-observed and affine predicted angular values while ordinate represents the difference between experimental-observed and affine predicted angular values. Median of the difference between two populations on the projection plot and ±*σ* spread (i.e., ±34.1% indicated by red-dots) about the median is used to estimate offset and range values respectively.

**Fig.S13.**
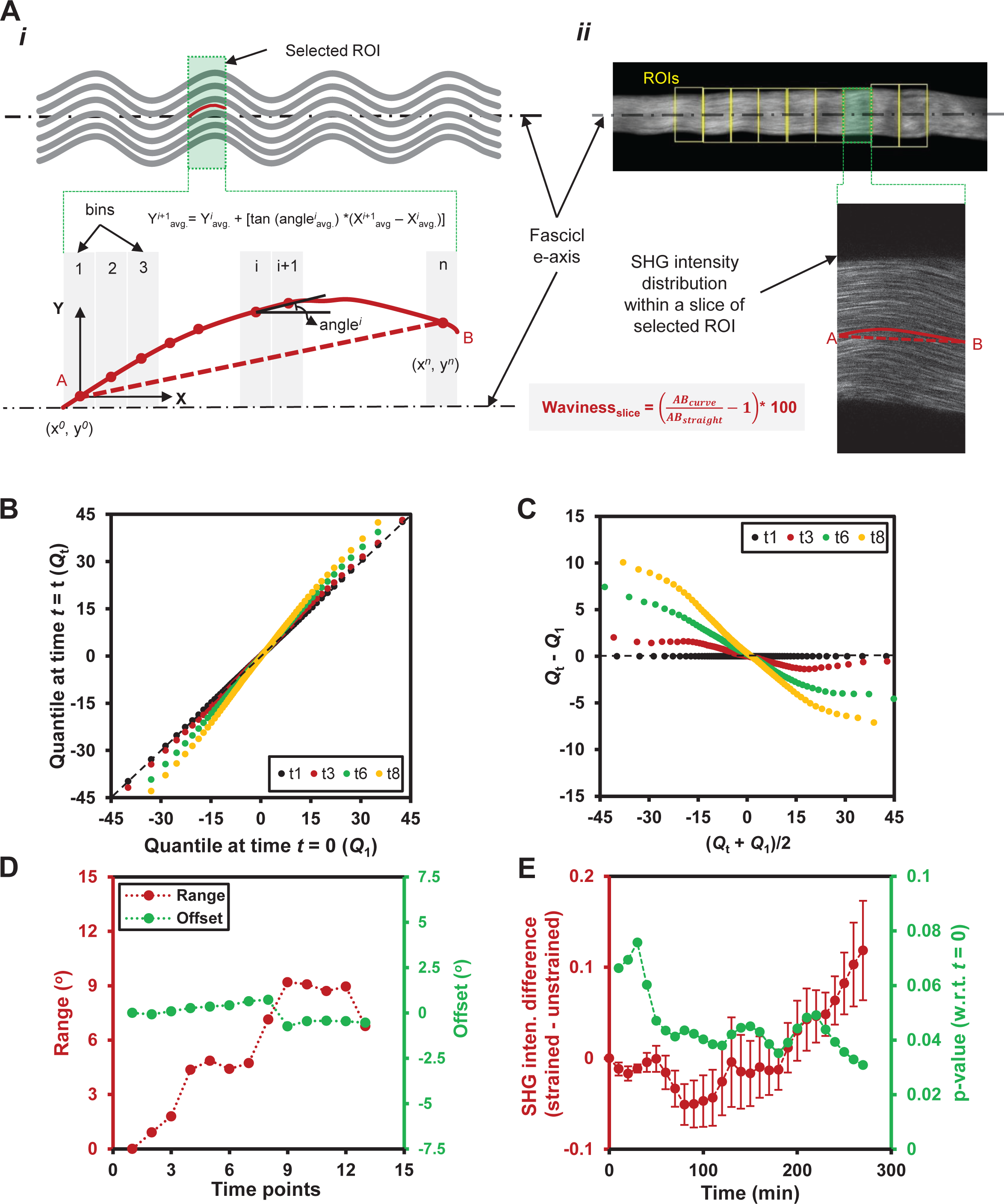
Measurement of waviness in the fascicle based on SHG signal and, temporal comparison of collagen fiber orientation distributions within a fascicle region using projection plots during degradation by BC. A. (i) Waviness measurement at a slice or image plane reflects a curvature of collagen fibers over each 50 µm and, is calculated based on spatial averaging of collagen fiber inclinations (w.r.t. the fascicle-axis). The median of slice-wise measurements of waviness values is used to represent the waviness of a ROI. (ii) Forward SHG image (projected sum intensity) of a tendon fascicle indicating various ROIs (top) and a representative slice within a ROI (bottom). **B.** Quantile-quantile (*Q-Q*) plot for collagen fiber orientation distributions within a representative ROI at four time points (i.e., time-point-1,3,6,8 of total 13 time-points spanning over ∼6 hours) during degradation by BC. The “fat-tails” on Q-Q plots suggest an appearance of more inclined fibers following the fascicle degradation by BC. **C.** Corresponding projection plots at same time points representing mean and difference of the observed quantiles. **D.** Range and offset magnitudes of a ROI at various time points shows that the offset values remains almost unaltered while range values change considerably. Median of the difference between two populations on the projection plot and ±*σ* spread (i.e., ±34.1% indicated by red-dots) about the median is used to estimate offset and range values (for methods, see Fig.S12). **E.** The difference between average SHG intensities of uniaxially strained fascicle and its unstrained counterpart under exposure to BC increases with time (averaged over three experiments, red markers). Following an initial phase of collagen degradation (∼1 hour), SHG intensities become significantly different (p-values < 0.05, green markers) from the initial SHG intensity values (i.e., *t* = 0) (n = 3 fascicles).

**Fig.S14.**
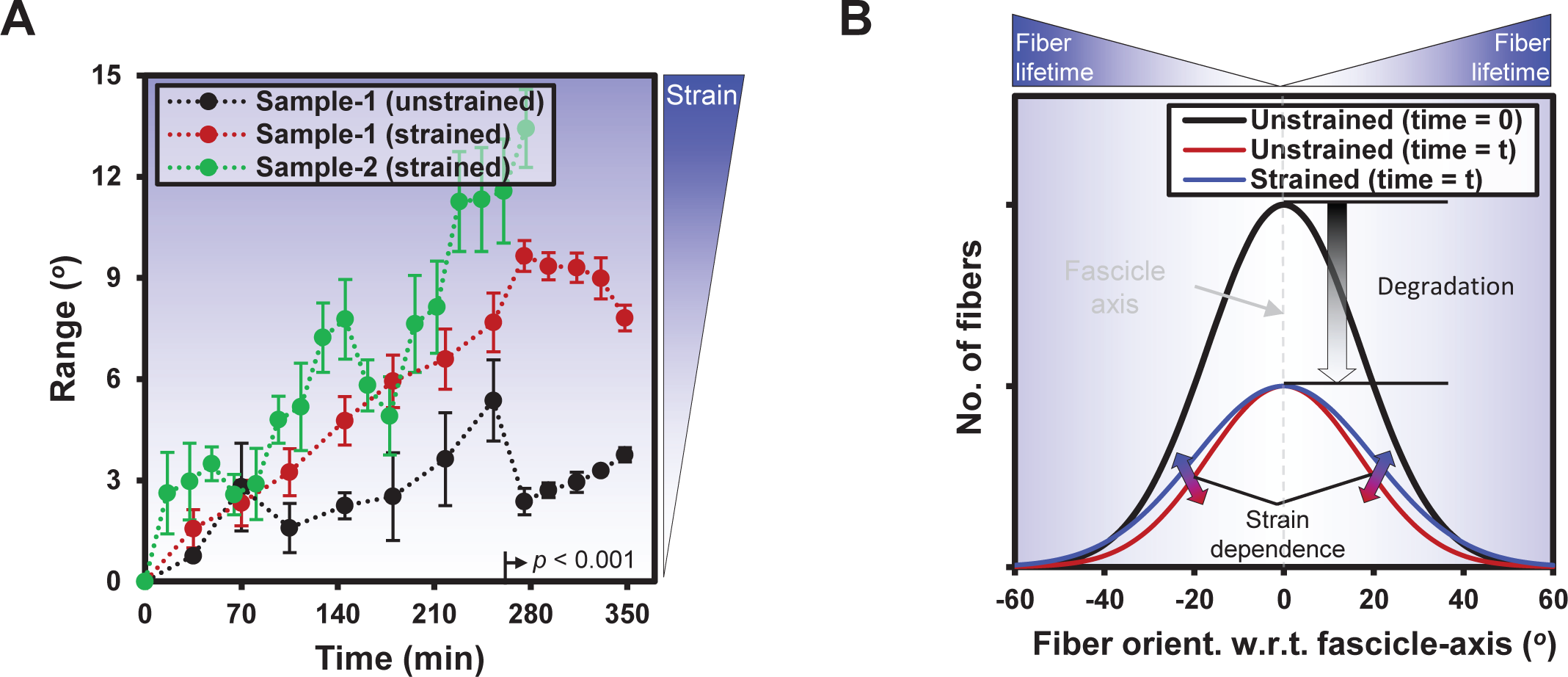
Enzymatic degradation (by BC) produces more inclined collagen fibers (w.r.t. the fascicle-axis) in uniaxially deformed tendon fascicles than its strain-free counterparts. **A.** Quantitative comparison of collagen fiber orientation distributions between strained and unstrained fascicles based on range values (obtained using projection plots, Fig.S12). Temporal increase in range values suggest more inclined fibers at later time points (i.e., >255 minutes) suggest mechanically strained fascicles end with more inclined fibers compared to unstrained counterparts. *p*-values based on Two-tailed t-test. **B.** Collagen fibers in an unstrained fascicle (marked as black-line) might follow uniform degradation (marked as red-line) as indicated by smaller change in range values with time (Fig.S14A). Mechanically strained fascicles ends up with more inclined fibers (marked as blue-line) relative to aligned ones as indicated by relatively larger range values with time.

**Fig.S15.**
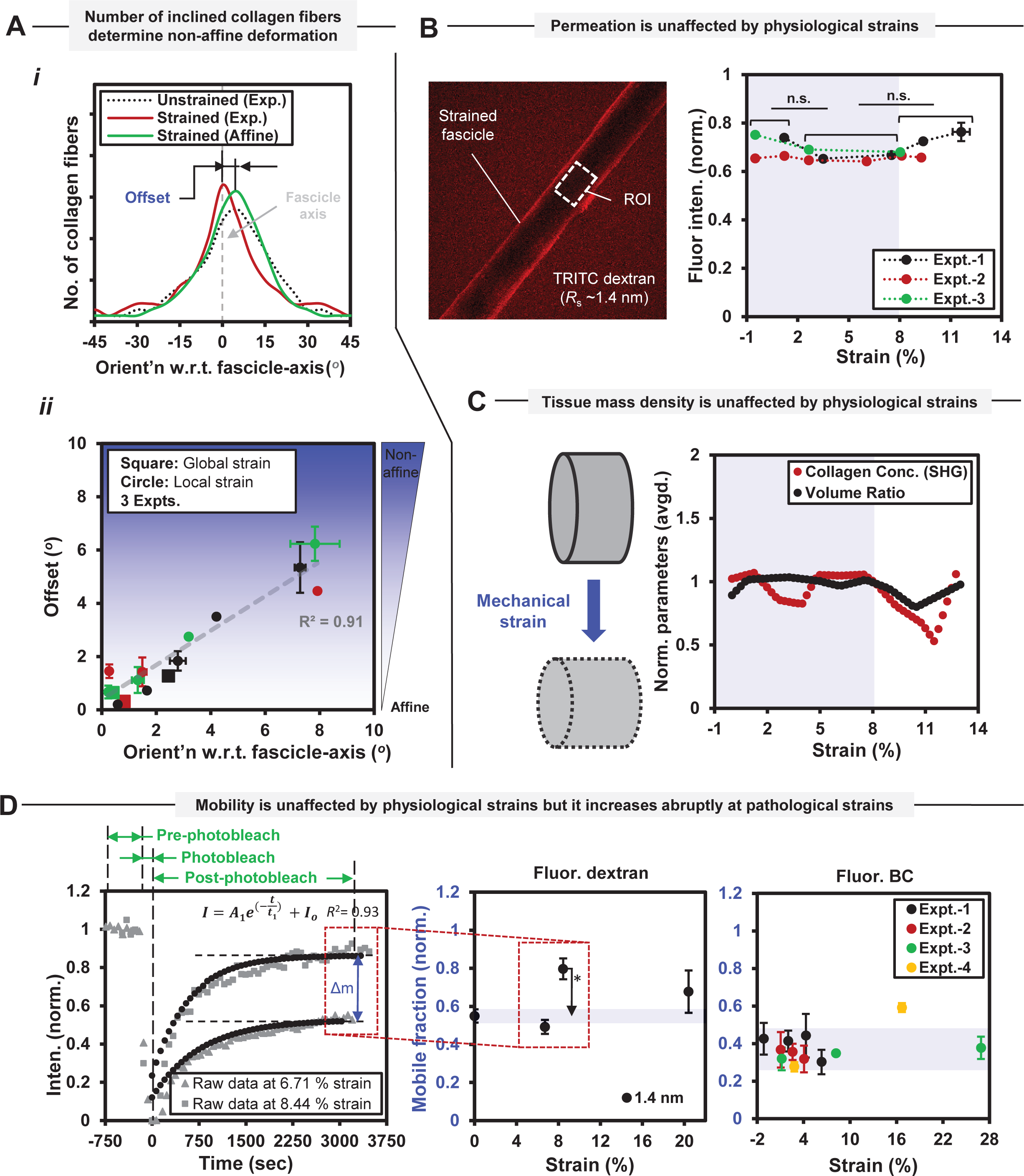
Non-affine strains within tendon fascicle do not affect permeation, mass density, or enzyme mobility. **A.** Non-affine deformation in a tendon fascicle region is determined by collagen fiber orientation distribution within undeformed tendon tissues. (i-ii) The extent of non-affine deformation, defined by the offset between collagen fiber orientation distributions within experimentally-observed and affine-predicted deformed configurations, correlates linearly with collagen fibers orientation value (relative to fascicle-axis) within undeformed tendon. n = 3 samples; Error bars indicate ±SEM. **B.** Tissue permeation is unaffected by up to ∼8% strain. (L to R) Representative heterogeneously deformed tendon fascicle (at ∼half-diameter depth) following an overnight exposure to TRITC-dextran (Stoke’s radius (*R_s_*) = 1.4 nm) with permeation values quantified using average fluorescence intensity (normalized w.r.t. dextran solution). n = 3 samples; Error bars indicate ±SEM; n.s.: non-significant based on two-tailed t-test *p* > 0.05. **C.** SHG based normalized collagen concentration and volume ratio values do not deviate systematically for strains of up to ∼8% as shown by piecewise line fits to the normalized parameter values. Product of collagen concentration and volume values shows tissue mass-density stays within ±10% of its average value (Fig.S19). n = 3 samples. **D.** Mobility of collagenase-size TRITC-dextran (R_s_ = 1.4 nm) or fluorescent-BC molecules is unaffected for up to ∼8% fascicles strain whereas the increases in mobile fraction values at larger strains (i.e., >8%) suggest a local tissue fracture. The mobility of relatively larger-size dextran molecules within the fascicles show strain-dependence based on fluorescence recovery after photobleaching (FRAP) (Fig.S22). BC molecules were fluorescently labelled at N-terminus (Fig.S21). n ≥ 3 samples per collagenase or TRITC dextran molecule; Error bars indicate ±SEM; Two-tailed *t*-test: n.s. = not significant, * = p < 0.05, ** = p < 0.01, *** = p < 0.001.

**Fig.S16.**
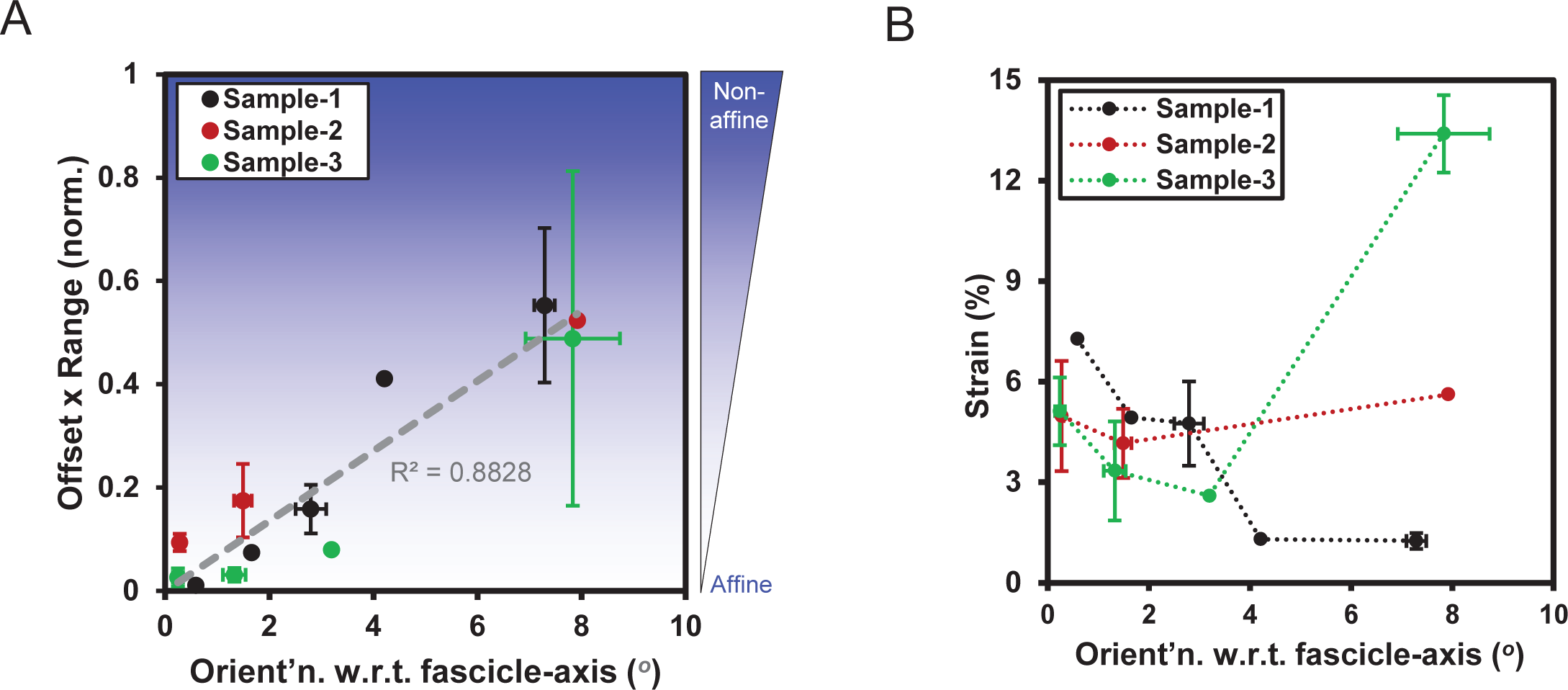
Local collagen fiber orientation distribution w.r.t. the fascicle-axis affects the extent of non-affine behavior and strain distribution in case of uniaxially deformed fascicles. **A.** When non-affine deformation behavior quantified as the product of offset and range values (in contrast to just offset values), the regions of the fascicle carrying aligned fibers (w.r.t. the fascicle-axis) follow affine deformation whereas the regions dominated by inclined collagen fibers tend to follow non-affine deformation. Offset and range values, estimated based on projection plots (Fig.S12D), are normalized w.r.t. maximum observed value within an experiment. (n = 3 fascicles; All error bars indicate ±SEM). **B.** Local strain magnitudes under uniaxial deformation of fascicle depends on the local collagen fiber orientation (w.r.t. fascicle-axis) in the undeformed configuration (n = 3 fascicles; All error bars indicate ±SEM).

**Fig.S17.**
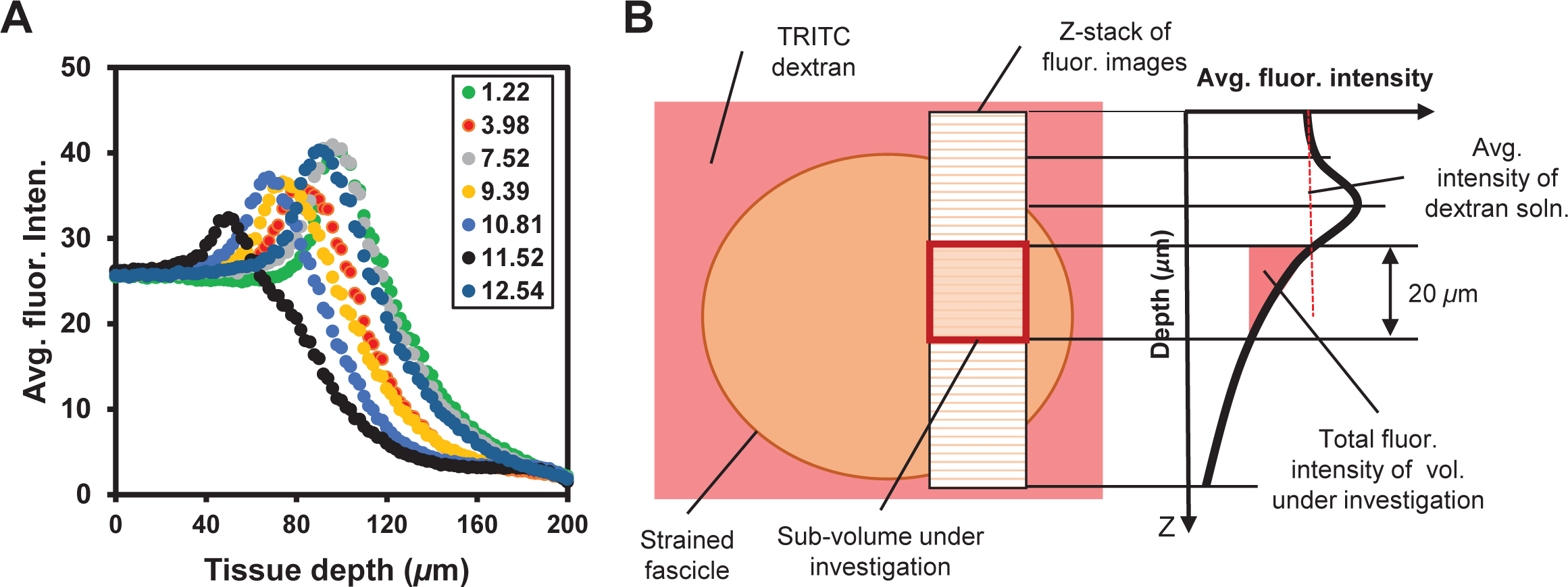
Quantification of permeation with different strain carrying regions of a heterogeneously deformed fascicle. **A.** Average fluorescent intensity with different ROIs of a heterogeneously deformed fascicle, kept overnight in TRITC-dextran (R_s_ = 1.4 nm), varied with the tissue depth (z-direction). Figure legends represent percent strain values within ROIs. **B.** Schematic of a fascicle’s cross-section and a representative curve showing area-averaged fluorescent intensity variation with the tissue depth. The observed peak is likely to result from the curvature of the fascicle surface. Volumetric averaging the fluorescent intensity measurements is used for comparison among different sub-volumes of a fascicle. Each sub-volume under investigation lies at a fixed distance from the fascicle surface and, a constant z-depth of 20 *µ*m is selected to represent the sub-volume (as larger z-depths were prone to fluorescent signal attenuation due to the sample thickness). Top surface of the sub-volume was selected at a z-position such that the average fluorescent intensity becomes equal to the average fluorescent intensity of free-dextran solution i.e., the point of intersection of average intensity with depth curve and line representing average fluorescent intensity of free-dextran solution.

**Fig.S18.**
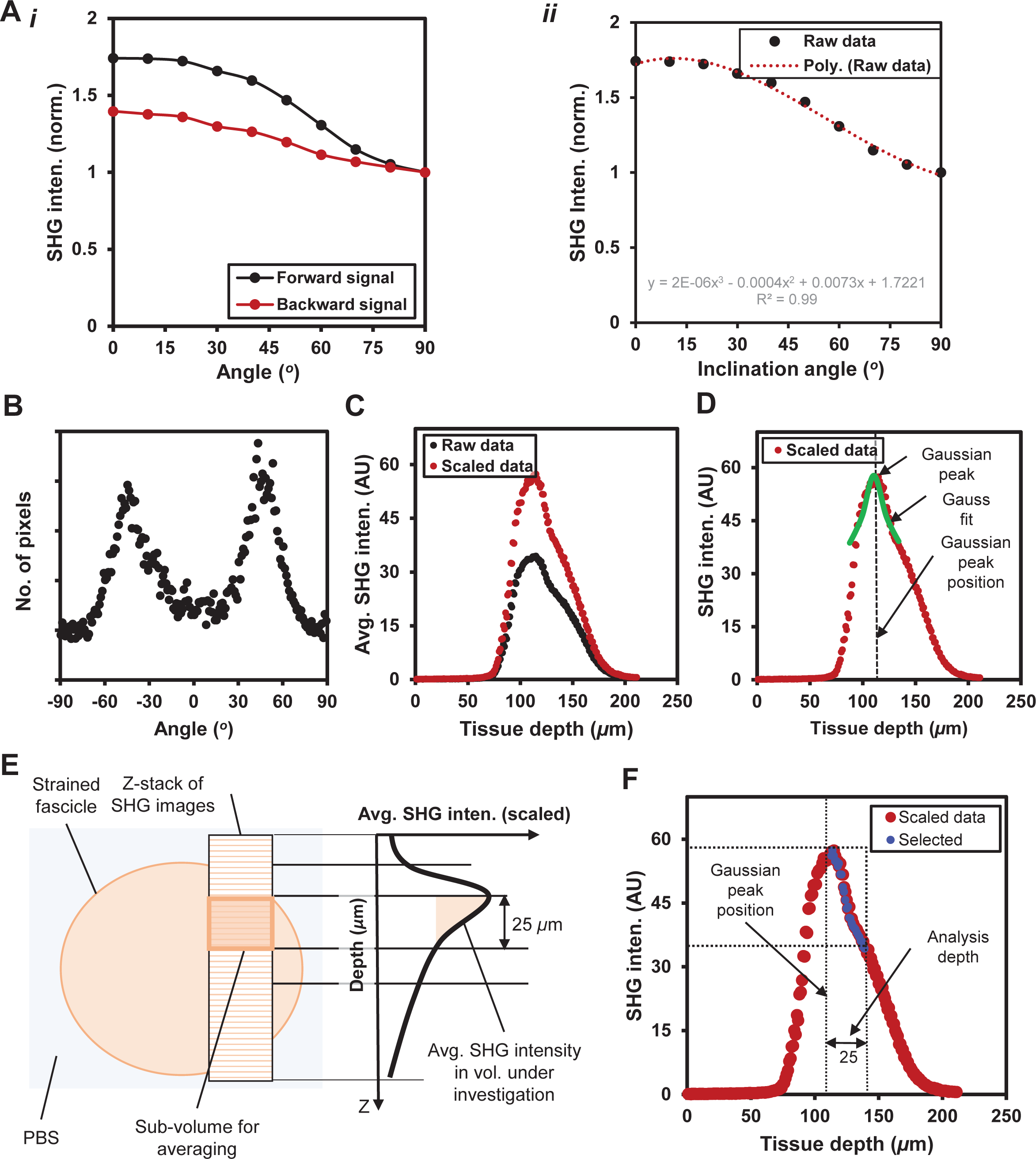
Measurement of absolute SHG intensity within a ROI of mechanically strained fascicle. **A.** (i) SHG intensity varies as a function of inclination angle between fascicle longitudinal-axis and the direction of incident light polarization. (ii) Curve fitting was done to obtain interpolated values of average SHG intensity at different angular positions. **B.** Pixel-wise collagen fiber orientation distribution, at an image plane based on structure tensor, shows that most collagen fibers are aligned along two different directions. **C.** The absolute SHG intensity of an image (marked as red-dots) is determined by the scaling of the experimentally observed average SHG intensity (i.e., raw intensity shown by black-dots). The scaling is based on collagen fiber orientation distribution and angular inclination (orientation of each pixel with the image w.r.t. the direction of incident light polarization) using the interpolation curve. Regions around the troughs in the tendon crimp pattern were excluded due to out of plane fiber alignment. **D.** The location of Gauss-peak on absolute SHG intensity versus tissue depth data has been used to predict the surface of the fascicle within each z-stack of a ROI. **E-F.** The position of sub-volume, used for volumetric averaging absolute SHG signal, considers Gauss-peak as sample surface while sub-volume depth is kept as 25 *µ*m (for all measurements). The selected points used for volumetric averaging are marked as blue-dots.

**Fig.S19.**
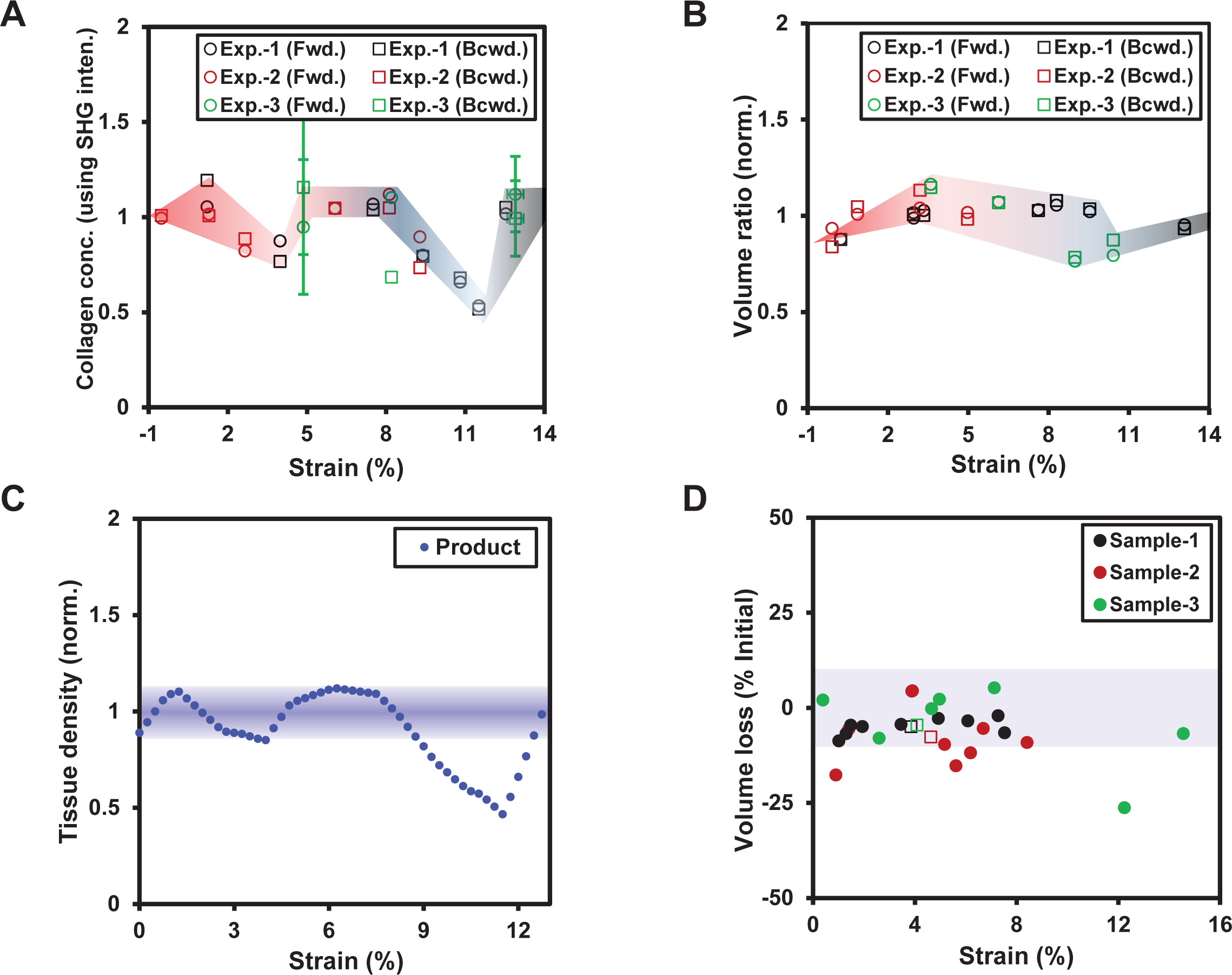
Tissue mass density is not significantly affected by physiological strains in the fascicles. **A.** Average collagen concentration for within a deformed ROIs is measured as square-root of the absolute SHG intensity (Fig.S18E,F). The estimated collagen concentrations at different strain values are normalized w.r.t. average SHG signal value with heterogeneously deformed fascicle (for strains values up to ∼8%). The estimated collagen concentrations based on forward and backward SHG intensities were averaged at each strain value (n = 3). **B.** The normalized volume ratio of a fascicle region is measured as the ratio of deformed volume to its initial volume. The region between parallel consecutive photobleach-stripes in a heterogeneously deformed fascicle was selected for each volume ratio measurement where the cross-sectional geometric limits of each region of the fascicle was represented by SHG intensity. The measurements based on forward and backward SHG intensities were averaged to obtain final volume ratio of a deformed fascicle region (n = 3). **C.** Local tissue mass density in a region of a heterogeneously deformed fascicle is defined as the product of collagen concentration in the deformed volume and volume ratio values. The product of piecewise lines fits to collagen concentration and volume ratio values, shows insignificant variation (< ±10% of its magnitude) of tissue mass density for fascicle strains up to ∼8%. **D.** Percentage volume loss based on Poisson’s ratio (PR) at increasing strain magnitudes within uniaxially deformed fascicle stays within ±10% of its average magnitude. Filled circles indicate local PR of each region for sample while empty squares indicate global PR values based on two extreme photobleach-stripes on the fascicles (n = 3)

**Fig.S20.**
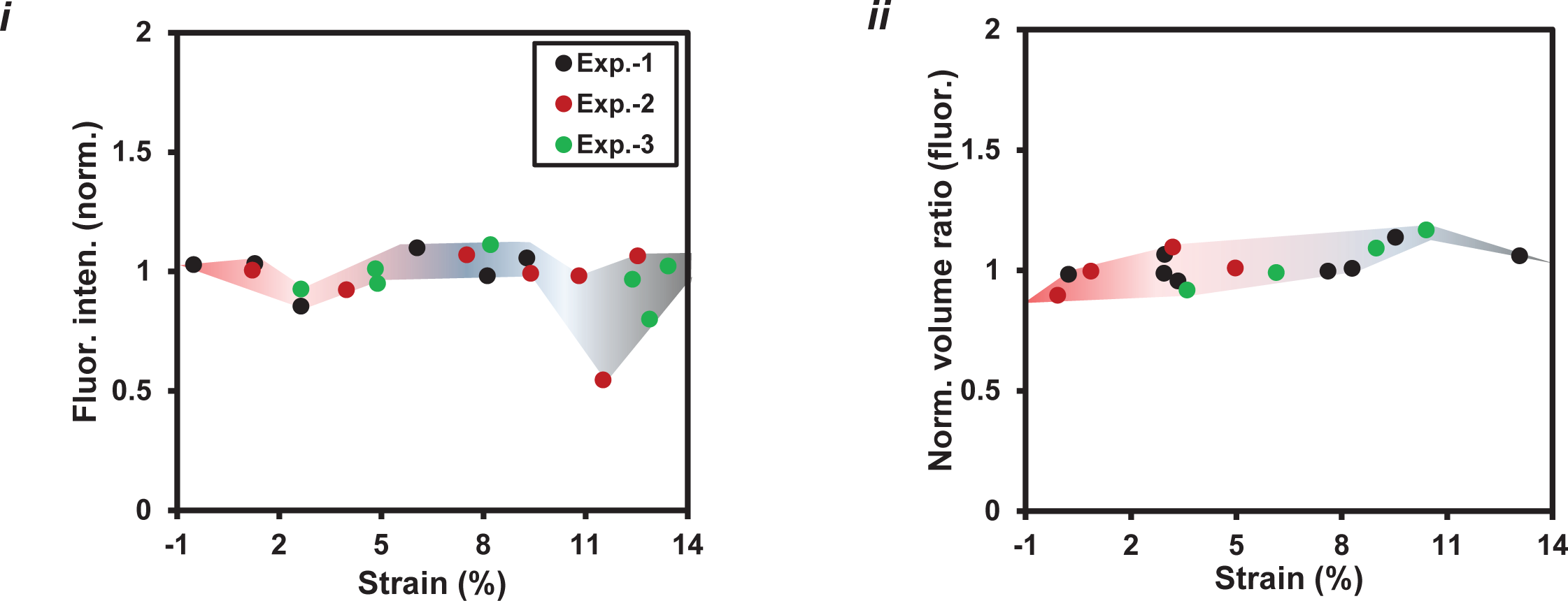
Collagen concentration and volumetric measurements within a fascicle based on fluorescent labelling by an exogenous fluorophore i.e., Col-F dye. (i-ii) Both collagen concentration and volume ratio values at different strain magnitudes shows a deviation from that of SHG based measurements (n = 3).

**Fig.S21.**
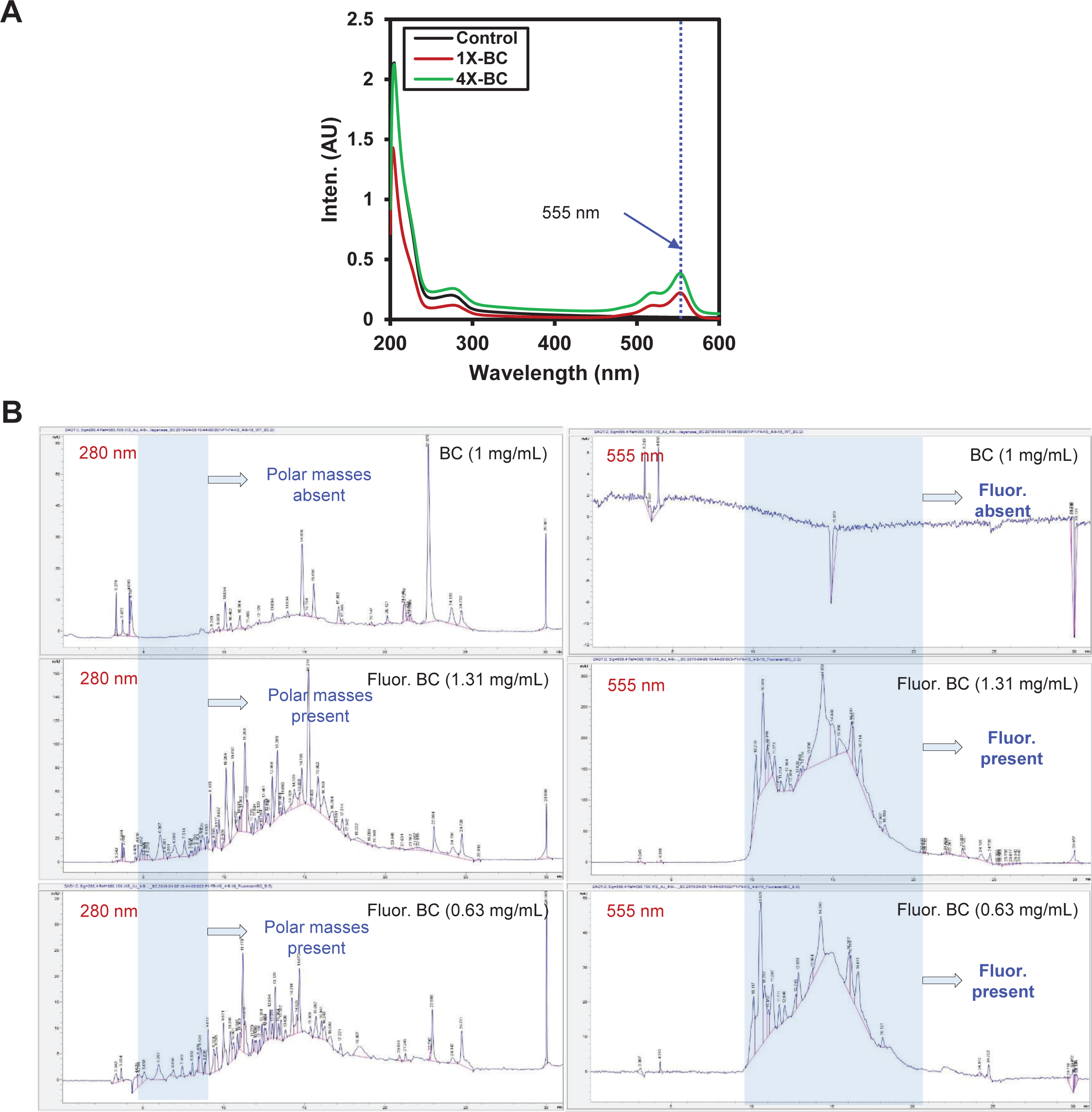
Fluorescent labelling of BC molecules at their N-terminus. **A.** Higher proportion of BC w.r.t. fluorophore (Alexa Fluor™ 555 NHS Ester) results in higher labelling efficiency as indicated by UV scan of fluorescently labelled molecules at 555 nm. 4X = 50 *μ*L of fluorophore + 500 *μ*L of BC (20 mg/mL); 1X = 50 *μ*L of fluorophore + 1000 *μ*L of BC (10 mg/mL). Fluorophore concentration = 10 mg/mL. **B.** Unlabelled and fluorescently labelled BC samples (at ∼1 mg/mL) on high-performance liquid chromatography (HPLC) show the presence of fluorescently labelled masses (at 555 nm). Peaks at 280 nm (highlighted in sky-blue) indicate presence of polar masses in fluorescent-BC samples resulting from self degradation of fluorescent-BC.

**Fig.S22.**
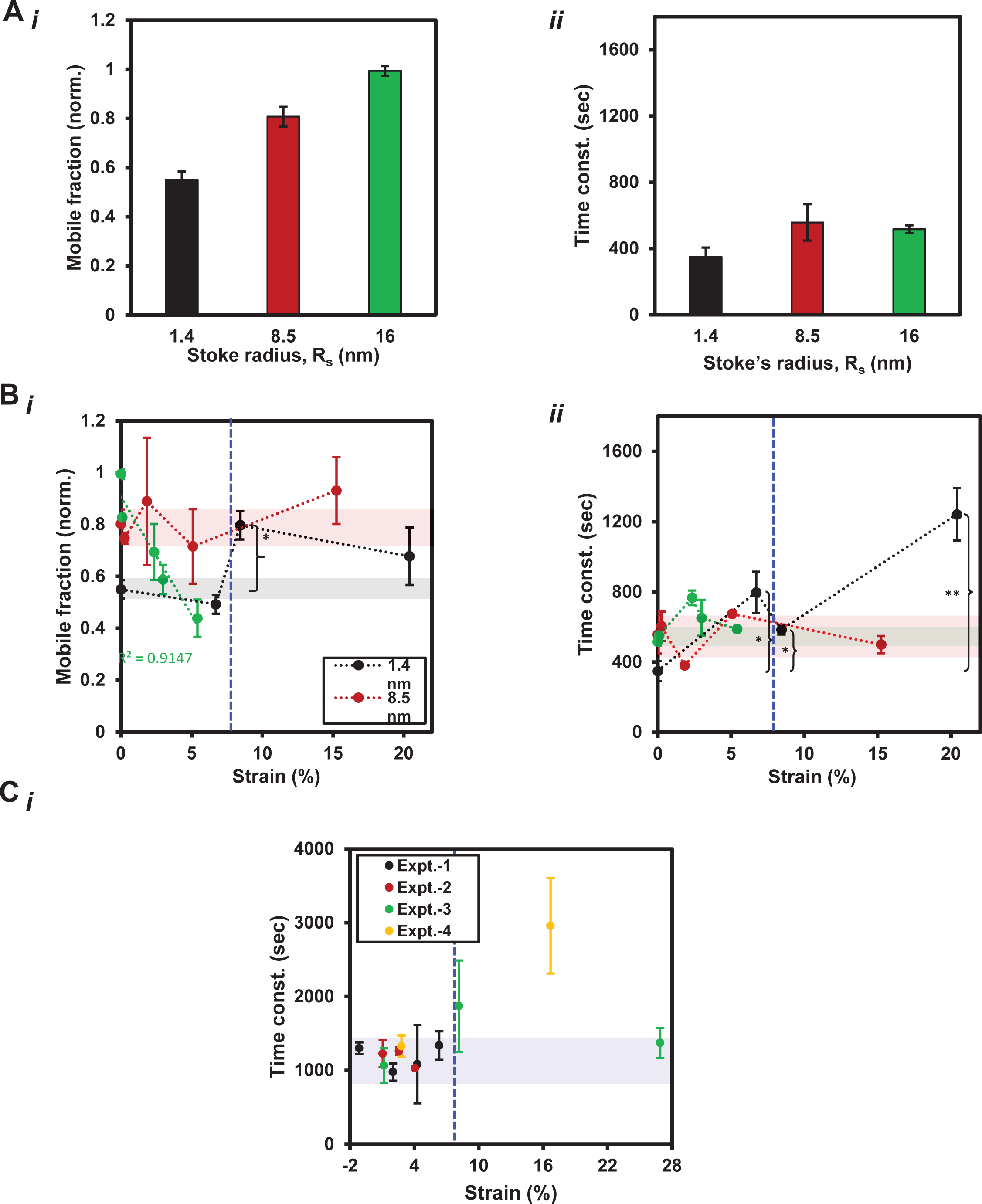
Mobility measurements of different size TRITC-dextran and fluorescent-BC molecules in mechanically deformed fascicles using FRAP method. **A.** Mobile fractions and time constants for different TRITC-dextran molecules (Stoke’s radii (R_s_) = 1.4 nm, 8.5 nm and 16 nm) in undeformed tendon fascicles. (i) Mobile fraction values increase with an increase in the size of TRITC-dextran molecules. (ii) Smaller-size TRITC-dextran molecules (R_s_ = 1.4 nm) move faster than the larger-size TRITC-dextran molecules (R_s_ = 8.5 nm and 16 nm) (n = 3; All error bars indicate ±SEM). **B.** Mobility of different size TRITC-dextran molecules within a mechanically strained fascicles is measured as a function the magnitude of mechanical strain. FRAP measurements of different size TRITC-dextran molecules (indicated by R_s_) are shown by (i) Mobile fraction and (ii) time constant values. Mechanical strain values up to ∼8% (indicated by blue line) do not influence mobile fraction of small size dextran molecules (R_s_ = 1.4 nm) while the mobile fraction values of larger size dextrans (R_s_ = 16 nm) decreases linearly with increasing mechanical strain values. Time constant values show that smaller dextran molecules (R_s_ = 1.4 nm) tend to move slower at increased strain values while the rate of mobility of the larger-size dextran molecules (R_s_ = 8.5 nm and 16 nm) do not change significantly with increasing strain magnitudes (n = 3 fascicles per TRITC dextran molecule; All error bars indicate ±SEM; * Two-tailed *t*-test: * = p < 0.05, ** = p < 0.01, *** = p < 0.001). **C.** Time constants of fluorescent-BC molecules are unaffected by strain values up to ∼8% (indicated by blue line) within the fascicles while larger strains (i.e., > 8%) cause an increase in the rates of mobility (n = 4 fascicles per collagenase molecule; All error bars indicate ±SEM; * Two-tailed *t*-test: * = p < 0.05, ** = p < 0.01, *** = p < 0.001) (n = 4 fascicles per collagenase molecule; All error bars indicate ±SEM).

**Fig.S23.**
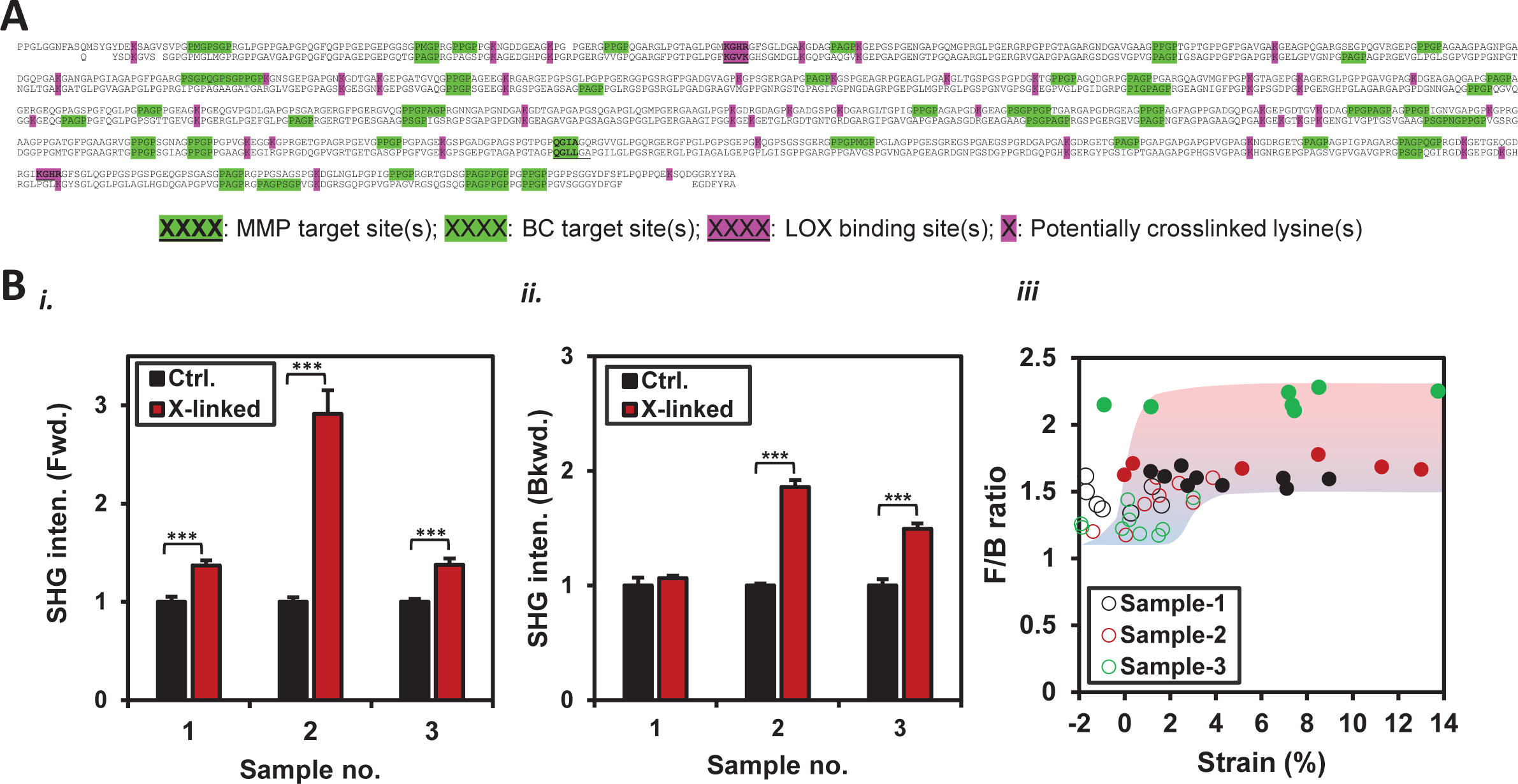
Structural changes in a mice tendon fascicle carrying artificial collagen crosslinks. **A.** Cleavage- (targeted by MMPs or BCs) and crosslink-sites (targeted by transglutaminase (TGM) and lysyl oxidases/hydroxylases (LOXs)) within aligned (using UniProt) alpha-1 and alpha-2 chains of collagen-I shown by FASTA sequence(s). Specie: mice. **B.** TGM-crosslinks induced changes in collagen fibril structure of unstrained tendon fascicles and the influence of mechanical strain on collagen fibrillar structure are captured by SHG signal. (i-ii) Significantly increased SHG signal in forward (Fwd.) and backward (Bkwd.) directions of a strain-free TGM-crosslinked fascicle is observed as compared to its negative control counterpart (i.e., without TGM-crosslinks, n = 3). (iii) F/B ratio magnitudes within uniaxially deformed TGM-crosslinked fascicles (represented by filled markers) are significantly different (with p < 0.001) from F/B ratio values of the strain-free control samples (i.e., without TGM-crosslinks and shown as empty markers). An altered inter-molecular spacing among collagen molecules within collagen fibrils is likely to contribute towards the observed changes in the SHG signal (n = 3 fascicles; All error bars indicate ±SEM; * Two-tailed *t*-test: * = p < 0.05, ** = p < 0.01, *** = p < 0.001).

**Fig.S24.**
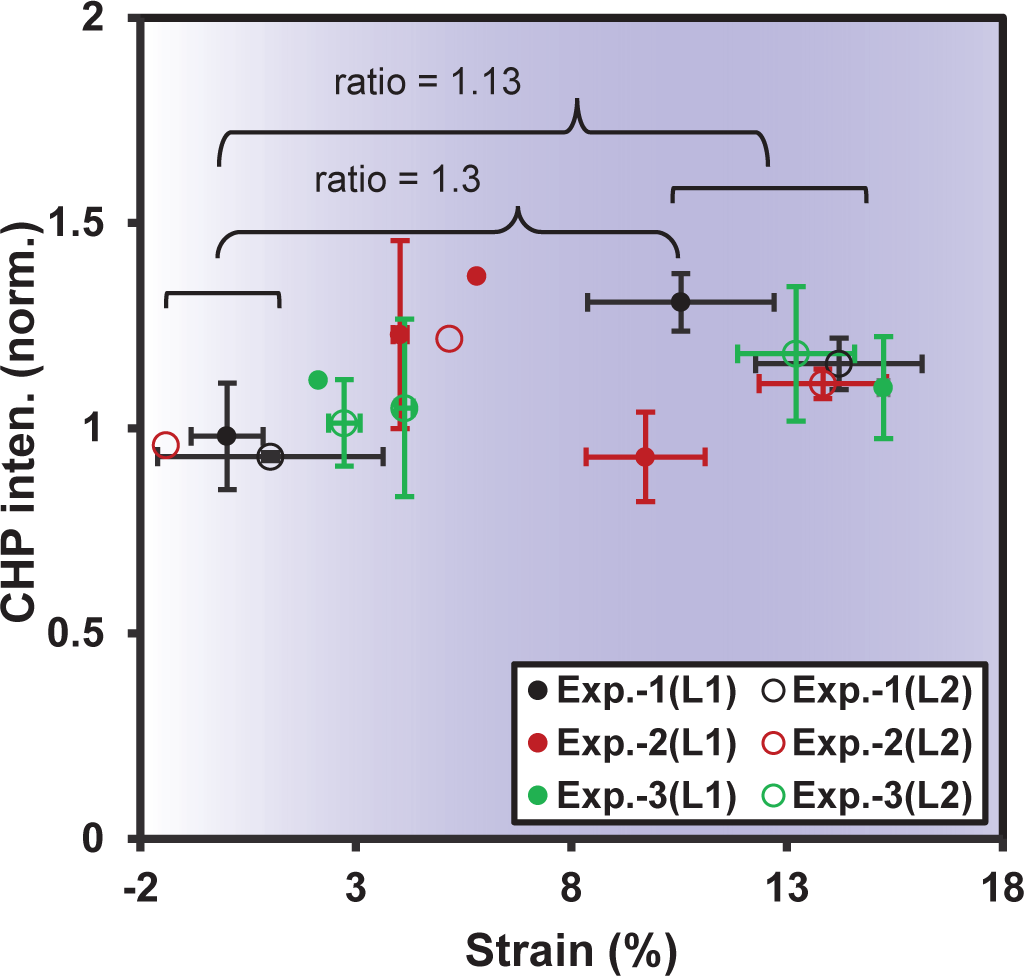
Collagen hybridizing peptide (CHP) shows increased binding towards the high strain regions of a heterogeneously deformed fascicle. Increased CHP intensity indicates an increased unfolded triple-helical collagen level at larger strains (i.e., > ∼8%) as compared to the physiologically strained fascicle regions (strains up to 8%). L1, L2 indicate two different locations of a fascicle situated more than 1000 um away from each other. (CHP = -[-glycine(G)–proline(P)–hydroxyproline(O)-]_c_; CF = carboxyfluorescein, c = 6-10) (n = 3 fascicles; Error bars indicate ±SEM).

**Fig.S25.**
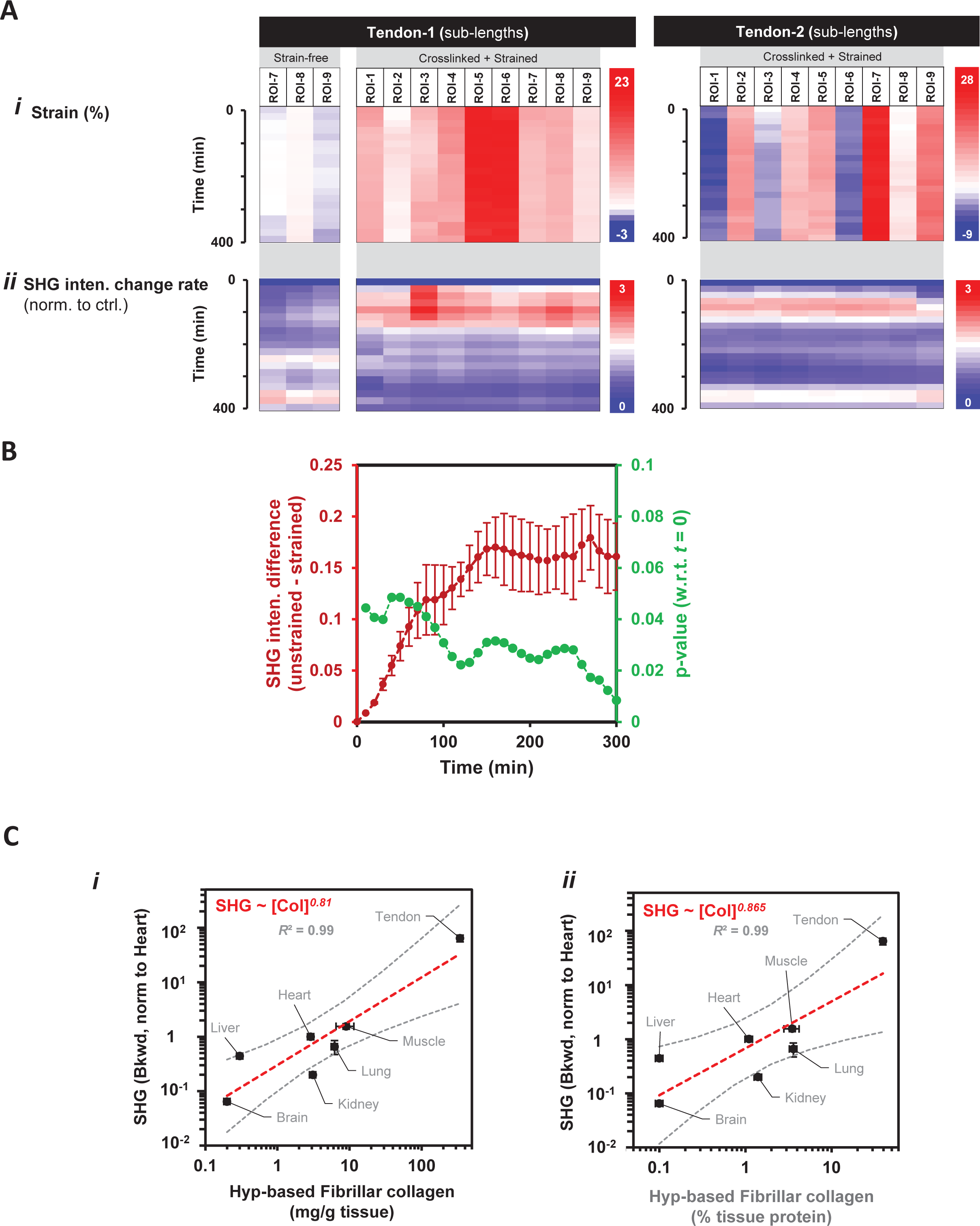
Uniaxially deformed TGM-crosslinked fascicle exposed to BC degrade much faster than its strain-free counterpart without TGM-crosslinks. **A.** TGM-crosslinked fascicle subjected to uniaxial deformation and subsequently exposed to BC for collagen degradation. Temporal values of (i) Strain magnitude and (ii) Collagen degradation rate (for calculations, see Fig.4C) for different ROIs of strain-free and strained fascicles show that the initial strain values remains almost constant while degradation rates increase significantly. **B.** The difference between average SHG intensities of strain-free and strained fascicles (TGM-crosslinked) increases with time (averaged over three experiments, red markers). Following collagen degradation by 1 hour, SHG intensities become significantly different (p-values < 0.05, green markers) from respective initial SHG intensity values (i.e., *t* = 0 for strained or unstrained sub-lengths) (n = 3 fascicles; All error bars indicate ±SEM). **C.** SHG signal scales with tissue fibrillar collagen level measured based on hydroxyproline (Hyp) content defined as either mg/g of wet weight or percentage total protein in tissue (per Tarnutzer et al., 2023 PMID: 36934197).

**Fig.S26:**
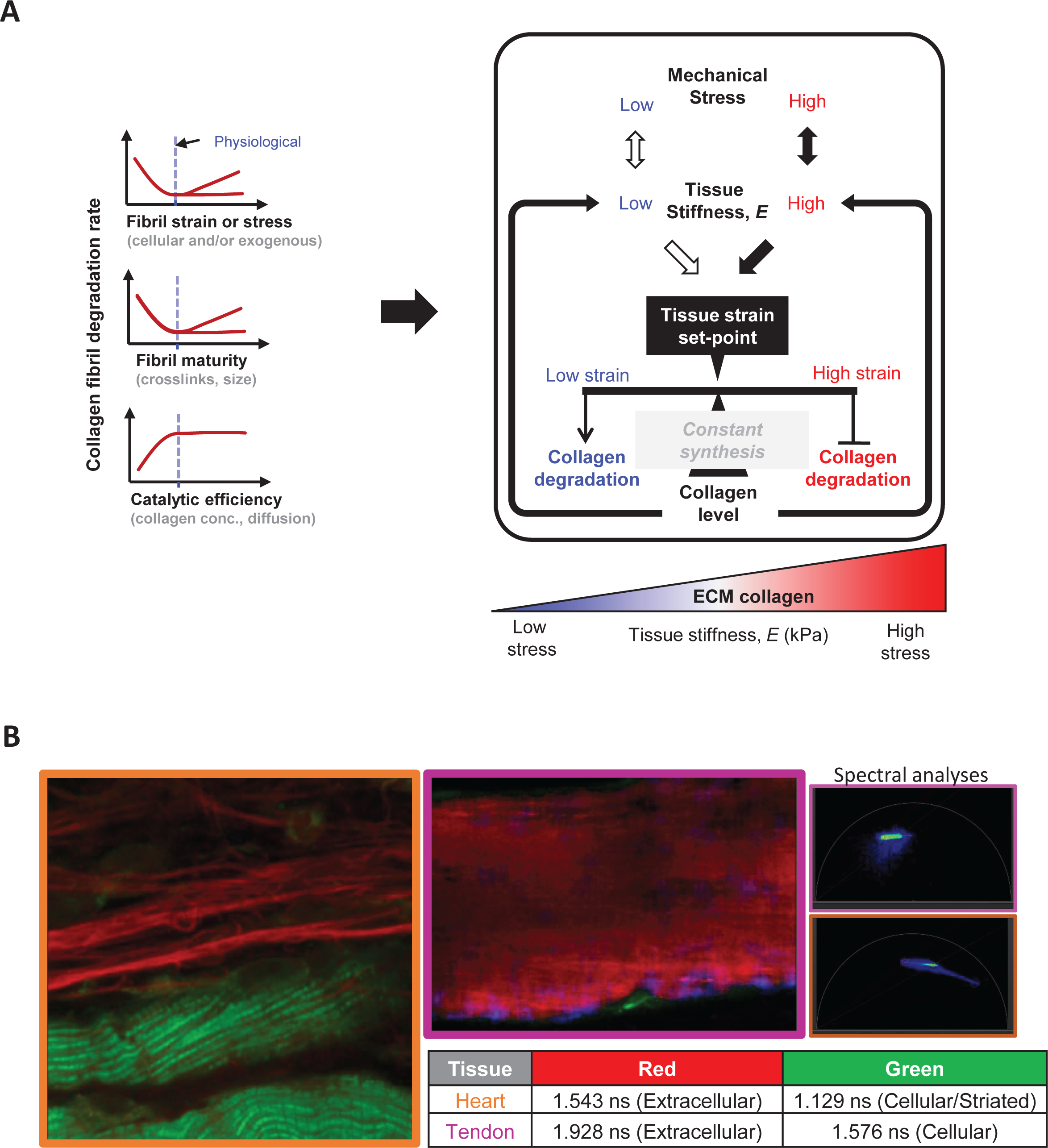
**A.** Working hypothesis for degradative sculpting of tissues. Physiologically, stiffer tissues withstanding larger mechanical stresses carry dense collagen than soft tissues sustaining lower mechanical stresses. Tissue strain set-point controls collagen levels during tissue remodeling irrespective of both density and synthesis of collagen. Strain values larger than strain set-point of a tissue tend to suppress its collagen degradation whereas lower strain values tend to promote collagen degradation **B.** Label-free auto-Fluorescence Lifetime Imaging (aFLIM) of heart and tendon tissues (isolated from adult mouse) (Ex: 440 nm; Em: 450-600 nm) for separating cellular and extra-cellular components of a tissue.

**Fig.S27.**
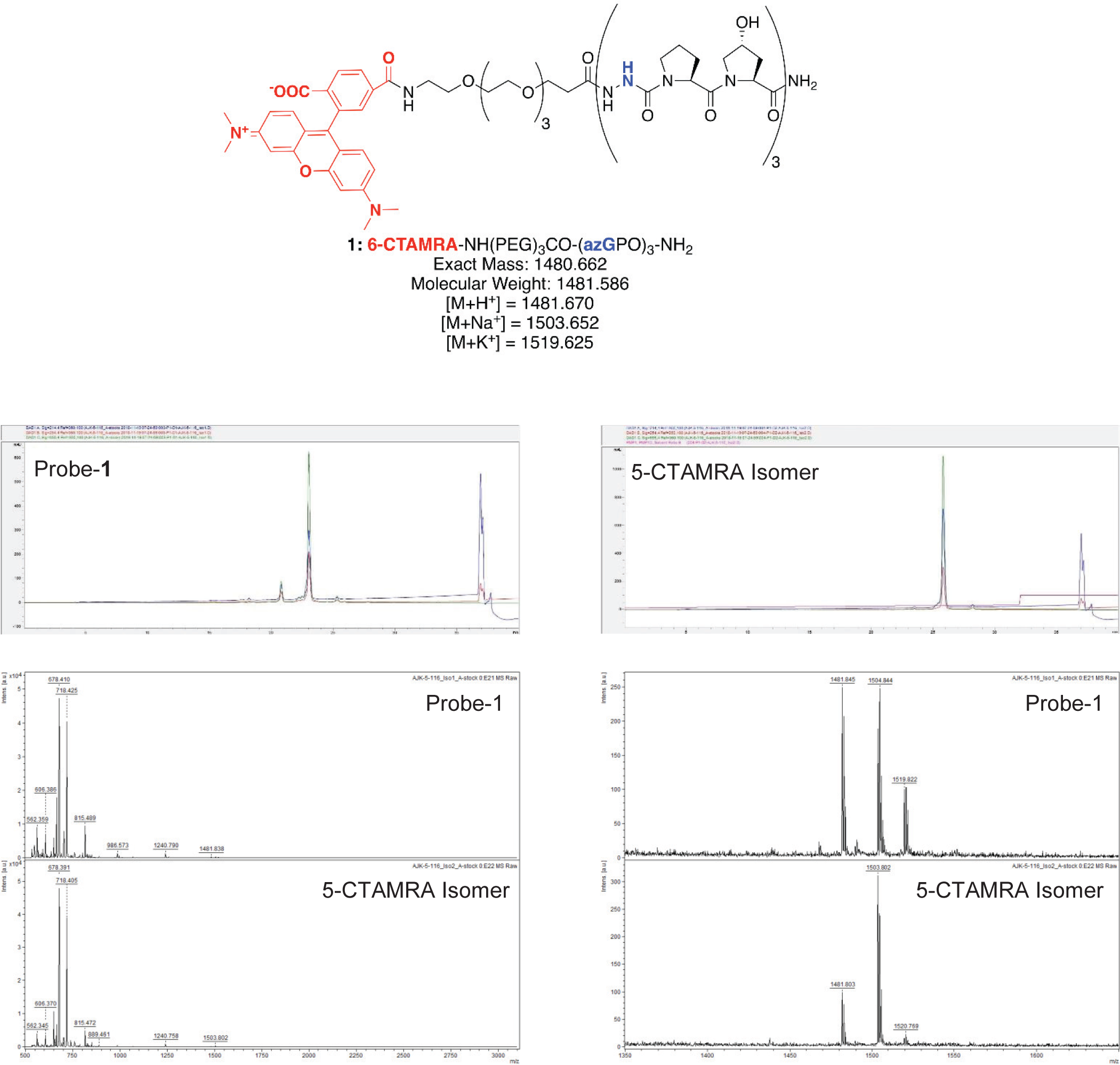
Validation data for Aza-peptide probe-1. (Top) Backbone structure and mass data, including predicted ion adducts for MALDI-TOF MS in positive ion mode. (Middle) HPLC trace for purified product (10-30% ACN/H_2_O (0.1% TFA), 80 °C, 30 min). (Bottom) MALDI-TOF MS spectrum represented in two views: wide mass range (left) and zoomed mass range (right), included to clarify the presence of ion adducts in this data.

**Fig.S28.**
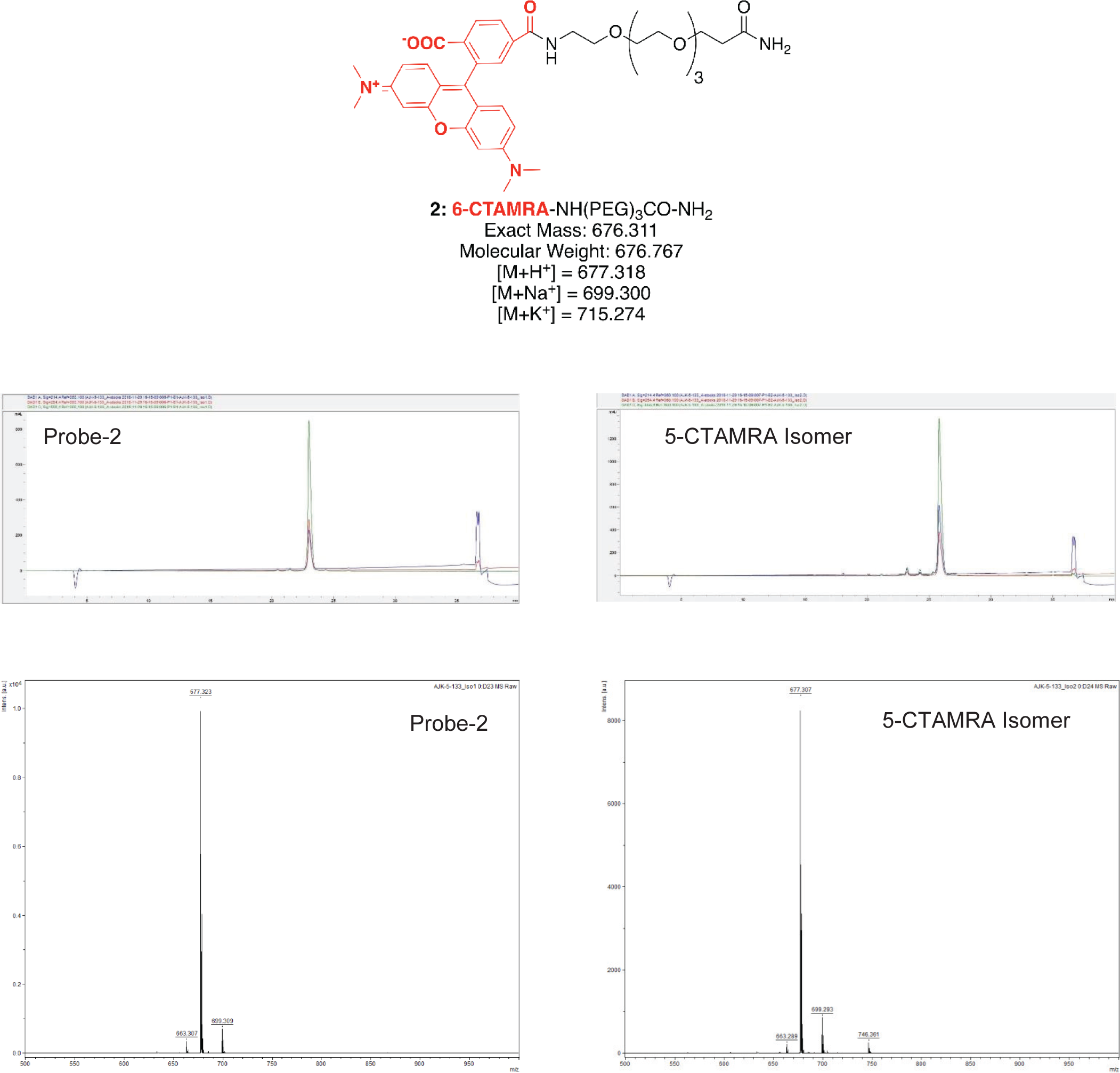
Validation data for 6-CTAMRA-PEG control probe-2. (Top) Backbone structure and mass data, including predicted ion adducts for MALDI-TOF MS in positive ion mode. (Middle) HPLC trace for purified product (10-30% ACN/H_2_O (0.1% TFA), 80 °C, 30 min). (Bottom) MALDI-TOF MS spectrum.

## References

1. Gennes, P.G.d., Scaling concepts in polymer physics. 1979, Ithaca, N.Y.: Cornell University Press. 324 p.

2. Swift, J., I.L. Ivanovska, A. Buxboim, T. Harada, P.C. Dingal, J. Pinter, J.D. Pajerowski, K.R. Spinler, J.W. Shin, M. Tewari, F. Rehfeldt, D.W. Speicher, and D.E. Discher, Nuclear lamin-A scales with tissue stiffness and enhances matrix-directed differentiation. Science, 2013. 341(6149): p. 1240104.

3. Vashisth, M., S. Cho, J. Irianto, Y. Xia, M. Wang, B. Hayes, D. Wieland, R. Wells, F. Jafarpour, A. Liu, and D.E. Discher, Scaling concepts in ‘omics: Nuclear lamin-B scales with tumor growth and often predicts poor prognosis, unlike fibrosis. Proc Natl Acad Sci U S A, 2021. 118(48).

4. Tabula Muris, C., c. Overall, c. Logistical, c. Organ, processing, p. Library, sequencing, a. Computational data, a. Cell type, g. Writing, g. Supplemental text writing, and i. Principal, Single-cell transcriptomics of 20 mouse organs creates a Tabula Muris. Nature, 2018. 562(7727): p. 367–372.

5. Yang, Y.L., L.M. Leone, and L.J. Kaufman, Elastic moduli of collagen gels can be predicted from two-dimensional confocal microscopy. Biophys J, 2009. 97(7): p. 2051–60.

6. Raub, C.B., A.J. Putnam, B.J. Tromberg, and S.C. George, Predicting bulk mechanical properties of cellularized collagen gels using multiphoton microscopy. Acta Biomater, 2010. 6(12): p. 4657–65.

7. Stein, A.M., D.A. Vader, D.A. Weitz, and L.M. Sander, The Micromechanics of Three-Dimensional Collagen-I Gels. Complexity, 2011. 16(4): p. 22–28.

8. Cho, S., M. Vashisth, A. Abbas, S. Majkut, K. Vogel, Y. Xia, I.L. Ivanovska, J. Irianto, M. Tewari, K. Zhu, E.D. Tichy, F. Mourkioti, H.-Y. Tang, R.A. Greenberg, B.L. Prosser, and D.E. Discher, Mechanosensing by the lamina protects against nuclear rupture, DNA damage, and cell cycle arrest. Dev. Cell., 2019. 49(6): p. 920–935.e5.

9. Chaudhuri, O., J. Cooper-White, P.A. Janmey, D.J. Mooney, and V.B. Shenoy, Effects of extracellular matrix viscoelasticity on cellular behaviour. Nature, 2020. 584(7822): p. 535–546.

10. Ivanovska, I.L., J. Swift, K. Spinler, D. Dingal, S. Cho, and D.E. Discher, Cross-linked matrix rigidity and soluble retinoids synergize in nuclear lamina regulation of stem cell differentiation. Mol Biol Cell, 2017. 28(14): p. 2010–2022.

11. Stolz, M., R. Gottardi, R. Raiteri, S. Miot, I. Martin, R. Imer, U. Staufer, A. Raducanu, M. Duggelin, W. Baschong, A.U. Daniels, N.F. Friederich, A. Aszodi, and U. Aebi, Early detection of aging cartilage and osteoarthritis in mice and patient samples using atomic force microscopy. Nat Nanotechnol, 2009. 4(3): p. 186–92.

12. Lopez-Garcia, M.D., D.J. Beebe, and W.C. Crone, Young’s modulus of collagen at slow displacement rates. Biomed Mater Eng, 2010. 20(6): p. 361–9.

13. Calo, A., Y. Romin, R. Srouji, C.P. Zambirinis, N. Fan, A. Santella, E. Feng, S. Fujisawa, M. Turkekul, S. Huang, A.L. Simpson, M. D’Angelica, W.R. Jarnagin, and K. Manova-Todorova, Spatial mapping of the collagen distribution in human and mouse tissues by force volume atomic force microscopy. Sci Rep, 2020. 10(1): p. 15664.

14. Konig, K., A. Ehlers, I. Riemann, S. Schenkl, R. Buckle, and M. Kaatz, Clinical two-photon microendoscopy. Microsc Res Tech, 2007. 70(5): p. 398–402.

15. Williams, R.M., W.R. Zipfel, and W.W. Webb, Interpreting second-harmonic generation images of collagen I fibrils. Biophys J, 2005. 88(2): p. 1377–86.

16. Lutz, V., M. Sattler, S. Gallinat, H. Wenck, R. Poertner, and F. Fischer, Characterization of fibrillar collagen types using multi-dimensional multiphoton laser scanning microscopy. Int J Cosmet Sci, 2012. 34(2): p. 209–15.

17. Majkut, S., T. Idema, J. Swift, C. Krieger, A. Liu, and D.E. Discher, Heart-specific stiffening in early embryos parallels matrix and myosin expression to optimize beating. Curr Biol, 2013. 23(23): p. 2434–9.

18. Green, E.M., H. Wakimoto, R.L. Anderson, M.J. Evanchik, J.M. Gorham, B.C. Harrison, M. Henze, R. Kawas, J.D. Oslob, H.M. Rodriguez, Y. Song, W. Wan, L.A. Leinwand, J.A. Spudich, R.S. McDowell, J.G. Seidman, and C.E. Seidman, A small-molecule inhibitor of sarcomere contractility suppresses hypertrophic cardiomyopathy in mice. Science, 2016. 351(6273): p. 617–21.

19. Mullard, A., FDA approves first cardiac myosin inhibitor. Nat Rev Drug Discov, 2022. 21(6): p. 406.

20. Harn, H.I., S.P. Wang, Y.C. Lai, B. Van Handel, Y.C. Liang, S. Tsai, I.M. Schiessl, A. Sarkar, H. Xi, M. Hughes, S. Kaemmer, M.J. Tang, J. Peti-Peterdi, A.D. Pyle, T.E. Woolley, D. Evseenko, T.X. Jiang, and C.M. Chuong, Symmetry breaking of tissue mechanics in wound induced hair follicle regeneration of laboratory and spiny mice. Nat Commun, 2021. 12(1): p. 2595.

21. Grgic, J., Use It or Lose It? A Meta-Analysis on the Effects of Resistance Training Cessation (Detraining) on Muscle Size in Older Adults. Int J Environ Res Public Health, 2022. 19(21).

22. Warden, S.J., S.M. Mantila Roosa, M.E. Kersh, A.L. Hurd, G.S. Fleisig, M.G. Pandy, and R.K. Fuchs, Physical activity when young provides lifelong benefits to cortical bone size and strength in men. Proc Natl Acad Sci U S A, 2014. 111(14): p. 5337–42.

23. Dewey, F.E., D. Rosenthal, D.J. Murphy, Jr., V.F. Froelicher, and E.A. Ashley, Does size matter? Clinical applications of scaling cardiac size and function for body size. Circulation, 2008. 117(17): p. 2279–87.

24. Heymsfield, S.B., T. Chirachariyavej, I.J. Rhyu, C. Roongpisuthipong, M. Heo, and A. Pietrobelli, Differences between brain mass and body weight scaling to height: potential mechanism of reduced mass-specific resting energy expenditure of taller adults. J Appl Physiol (1985), 2009. 106(1): p. 40–8.

25. Tominari, T., R. Ichimaru, K. Taniguchi, A. Yumoto, M. Shirakawa, C. Matsumoto, K. Watanabe, M. Hirata, Y. Itoh, D. Shiba, C. Miyaura, and M. Inada, Hypergravity and microgravity exhibited reversal effects on the bone and muscle mass in mice. Sci Rep, 2019. 9(1): p. 6614.

26. Walls, S., S. Diop, R. Birse, L. Elmen, Z. Gan, S. Kalvakuri, S. Pineda, C. Reddy, E. Taylor, B. Trinh, G. Vogler, R. Zarndt, A. McCulloch, P. Lee, S. Bhattacharya, R. Bodmer, and K. Ocorr, Prolonged Exposure to Microgravity Reduces Cardiac Contractility and Initiates Remodeling in Drosophila. Cell Rep, 2020. 33(10): p. 108445.

27. Prange, H.D., J.F. Anderson, and H. Rahn, Scaling of Skeletal Mass to Body Mass in Birds and Mammals. The American Naturalist, 1979. 113(1): p. 103–122.

28. Muchlinski, M.N., J.J. Snodgrass, and C.J. Terranova, Muscle mass scaling in primates: an energetic and ecological perspective. Am J Primatol, 2012. 74(5): p. 395–407.

29. Stahl, W.R., Organ weights in primates and other mammals. Science, 1965. 150(3699): p. 1039–42.

30. Wang, Z., P. Deurenberg, W. Wang, A. Pietrobelli, R.N. Baumgartner, and S.B. Heymsfield, Hydration of fat-free body mass: review and critique of a classic body-composition constant. Am J Clin Nutr, 1999. 69(5): p. 833–41.

31. Flynn, B.P., A.P. Bhole, N. Saeidi, M. Liles, C.A. Dimarzio, and J.W. Ruberti, Mechanical strain stabilizes reconstituted collagen fibrils against enzymatic degradation by mammalian collagenase matrix metalloproteinase 8 (MMP-8). PLoS One, 2010. 5(8): p. e12337.

32. Saini, K., S. Cho, L.J. Dooling, and D.E. Discher, Tension in fibrils suppresses their enzymatic degradation - A molecular mechanism for ‘use it or lose it’. Matrix Biol, 2019.

33. Huang, C. and I.V. Yannas, Mechanochemical studies of enzymatic degradation of insoluble collagen fibers. J Biomed Mater Res, 1977. 11(1): p. 137–54.

34. Nabeshima, Y., E.S. Grood, A. Sakurai, and J.H. Herman, Uniaxial tension inhibits tendon collagen degradation by collagenase in vitro. J Orthop Res, 1996. 14(1): p. 123–30.

35. Wyatt, K.E., J.W. Bourne, and P.A. Torzilli, Deformation-dependent enzyme mechanokinetic cleavage of type I collagen. J Biomech Eng, 2009. 131(5): p. 051004.

36. Ruberti, J.W. and N.J. Hallab, Strain-controlled enzymatic cleavage of collagen in loaded matrix. Biochem Biophys Res Commun, 2005. 336(2): p. 483–9.

37. Zareian, R., K.P. Church, N. Saeidi, B.P. Flynn, J.W. Beale, and J.W. Ruberti, Probing collagen/enzyme mechanochemistry in native tissue with dynamic, enzyme-induced creep. Langmuir, 2010. 26(12): p. 9917–26.

38. Chung, L., D. Dinakarpandian, N. Yoshida, J.L. Lauer-Fields, G.B. Fields, R. Visse, and H. Nagase, Collagenase unwinds triple-helical collagen prior to peptide bond hydrolysis. EMBO J, 2004. 23(15): p. 3020–30.

39. Sarkar, S.K., B. Marmer, G. Goldberg, and K.C. Neuman, Single-molecule tracking of collagenase on native type I collagen fibrils reveals degradation mechanism. Curr Biol, 2012. 22(12): p. 1047–56.

40. Plotnikov, S.V., A.C. Millard, P.J. Campagnola, and W.A. Mohler, Characterization of the myosin-based source for second-harmonic generation from muscle sarcomeres. Biophys J, 2006. 90(2): p. 693–703.

41. Xu, H., T. Liang, L. Wei, J.C. Zhu, X. Liu, C.C. Ji, B. Liu, and Z.P. Luo, Nano-elastic modulus of tendon measured directly in living mice. J Biomech, 2021. 116: p. 110248.

42. Aimes, R.T. and J.P. Quigley, Matrix metalloproteinase-2 is an interstitial collagenase. Inhibitor-free enzyme catalyzes the cleavage of collagen fibrils and soluble native type I collagen generating the specific 3/4- and 1/4-length fragments. J Biol Chem, 1995. 270(11): p. 5872–6.

43. Smith, G.N., Jr., K.D. Brandt, and K.A. Hasty, Procollagenase is reduced to inactive fragments upon activation in the presence of doxycycline. Ann N Y Acad Sci, 1994. 732: p. 436–8.

44. Kannus, P., Structure of the tendon connective tissue. Scand J Med Sci Sports, 2000. 10(6): p. 312–20.

45. Zitnay, J.L., Y. Li, Z. Qin, B.H. San, B. Depalle, S.P. Reese, M.J. Buehler, S.M. Yu, and J.A. Weiss, Molecular level detection and localization of mechanical damage in collagen enabled by collagen hybridizing peptides. Nat Commun, 2017. 8: p. 14913.

46. Zega, A., Azapeptides as pharmacological agents. Curr Med Chem, 2005. 12(5): p. 589–97.

47. Fratzl, P., K. Misof, I. Zizak, G. Rapp, H. Amenitsch, and S. Bernstorff, Fibrillar structure and mechanical properties of collagen. J Struct Biol, 1998. 122(1-2): p. 119–22.

48. Diamant, J., A. Keller, E. Baer, M. Litt, and R.G. Arridge, Collagen; ultrastructure and its relation to mechanical properties as a function of ageing. Proc R Soc Lond B Biol Sci, 1972. 180(1060): p. 293–315.

49. Garnero, P., O. Borel, I. Byrjalsen, M. Ferreras, F.H. Drake, M.S. McQueney, N.T. Foged, P.D. Delmas, and J.M. Delaisse, The collagenolytic activity of cathepsin K is unique among mammalian proteinases. J Biol Chem, 1998. 273(48): p. 32347–52.

50. Kirkness, M.W.H. and N.R. Forde, Single-Molecule Assay for Proteolytic Susceptibility: Force-Induced Collagen Destabilization. Biophys J, 2018. 114(3): p. 570–576.

51. Lake, S.P., D.H. Cortes, J.A. Kadlowec, L.J. Soslowsky, and D.M. Elliott, Evaluation of affine fiber kinematics in human supraspinatus tendon using quantitative projection plot analysis. Biomech Model Mechanobiol, 2012. 11(1-2): p. 197–205.

52. Raub, C.B., B.J. Tromberg, and S.C. George, Second-Harmonic Generation Imaging of Self-Assembled Collagen Gels.

53. Tyn, M.T. and T.W. Gusek, Prediction of diffusion coefficients of proteins. Biotechnol Bioeng, 1990. 35(4): p. 327–38.

54. Peter, F. and W. Richard, Nature’s hierarchical materials. Progress in Materials Science, 2007. 52(8): p. 1263–1334.

55. Marc André, M., C. Po-Yu, L. Albert Yu-Min, and S. Yasuaki, Biological materials: Structure and mechanical properties. Progress in Materials Science, 2008. 53(1): p. 1–206.

56. Bourne, J.W. and P.A. Torzilli, Molecular simulations predict novel collagen conformations during cross-link loading. Matrix Biol, 2011. 30(5-6): p. 356–60.

57. Bourne, J.W., J.M. Lippell, and P.A. Torzilli, Glycation cross-linking induced mechanical-enzymatic cleavage of microscale tendon fibers. Matrix Biol, 2014. 34: p. 179–84.

58. Adhikari, A.S., J. Chai, and A.R. Dunn, Mechanical load induces a 100-fold increase in the rate of collagen proteolysis by MMP-1. J Am Chem Soc, 2011. 133(6): p. 1686–9.

59. Adhikari, A.S., E. Glassey, and A.R. Dunn, Conformational dynamics accompanying the proteolytic degradation of trimeric collagen I by collagenases. J Am Chem Soc, 2012. 134(32): p. 13259–65.

60. Chang, S.W., B.P. Flynn, J.W. Ruberti, and M.J. Buehler, Molecular mechanism of force induced stabilization of collagen against enzymatic breakdown. Biomaterials, 2012. 33(15): p. 3852–9.

61. Dingal, P.C. and D.E. Discher, Systems mechanobiology: tension-inhibited protein turnover is sufficient to physically control gene circuits. Biophys J, 2014. 107(11): p. 2734–43.

62. Dooling, L.J., K. Saini, A.A. Anlas, and D.E. Discher, Tissue mechanics coevolves with fibrillar matrisomes in healthy and fibrotic tissues. Matrix Biol, 2022. 111: p. 153–188.

63. Bohm, S., F. Mersmann, M. Tettke, M. Kraft, and A. Arampatzis, Human Achilles tendon plasticity in response to cyclic strain: effect of rate and duration. J Exp Biol, 2014. 217(Pt 22): p. 4010–7.

64. Kuruvilla, S.J., S.D. Fox, D.M. Cullen, and M.P. Akhter, Site specific bone adaptation response to mechanical loading. J Musculoskelet Neuronal Interact, 2008. 8(1): p. 71–8.

65. Abilez, O.J., E. Tzatzalos, H. Yang, M.T. Zhao, G. Jung, A.M. Zollner, M. Tiburcy, J. Riegler, E. Matsa, P. Shukla, Y. Zhuge, T. Chour, V.C. Chen, P.W. Burridge, I. Karakikes, E. Kuhl, D. Bernstein, L.A. Couture, J.D. Gold, W.H. Zimmermann, and J.C. Wu, Passive Stretch Induces Structural and Functional Maturation of Engineered Heart Muscle as Predicted by Computational Modeling. Stem Cells, 2018. 36(2): p. 265–277.

66. Davis, J., L.C. Davis, R.N. Correll, C.A. Makarewich, J.A. Schwanekamp, F. Moussavi-Harami, D. Wang, A.J. York, H. Wu, S.R. Houser, C.E. Seidman, J.G. Seidman, M. Regnier, J.M. Metzger, J.C. Wu, and J.D. Molkentin, A Tension-Based Model Distinguishes Hypertrophic versus Dilated Cardiomyopathy. Cell, 2016. 165(5): p. 1147–1159.

67. Langberg, H., H. Ellingsgaard, T. Madsen, J. Jansson, S.P. Magnusson, P. Aagaard, and M. Kjaer, Eccentric rehabilitation exercise increases peritendinous type I collagen synthesis in humans with Achilles tendinosis. Scand J Med Sci Sports, 2007. 17(1): p. 61–6.

68. Phrommintikul, A., L. Tran, A. Kompa, B. Wang, A. Adrahtas, D. Cantwell, D.J. Kelly, and H. Krum, Effects of a Rho kinase inhibitor on pressure overload induced cardiac hypertrophy and associated diastolic dysfunction. Am J Physiol Heart Circ Physiol, 2008. 294(4): p. H1804–14.

69. Chen, X., O. Nadiarynkh, S. Plotnikov, and P.J. Campagnola, Second harmonic generation microscopy for quantitative analysis of collagen fibrillar structure. Nat Protoc, 2012. 7(4): p. 654–69.

70. Theodossiou, T., G.S. Rapti, V. Hovhannisyan, E. Georgiou, K. Politopoulos, and D. Yova, Thermally induced irreversible conformational changes in collagen probed by optical second harmonic generation and laser-induced fluorescence. Lasers Med Sci, 2002. 17(1): p. 34–41.

71. Dittmore, A., J. Silver, S.K. Sarkar, B. Marmer, G.I. Goldberg, and K.C. Neuman, Internal strain drives spontaneous periodic buckling in collagen and regulates remodeling. Proc Natl Acad Sci U S A, 2016. 113(30): p. 8436–41.

72. Legerlotz, K., J. Dorn, J. Richter, M. Rausch, and O. Leupin, Age-dependent regulation of tendon crimp structure, cell length and gap width with strain. Acta Biomater, 2014. 10(10): p. 4447–55.

73. Szczesny, S.E. and D.M. Elliott, Interfibrillar shear stress is the loading mechanism of collagen fibrils in tendon. Acta Biomater, 2014. 10(6): p. 2582–90.

74. Zuskov, A., B.R. Freedman, J.A. Gordon, J.J. Sarver, M.R. Buckley, and L.J. Soslowsky, Tendon Biomechanics and Crimp Properties Following Fatigue Loading Are Influenced by Tendon Type and Age in Mice. J Orthop Res, 2020. 38(1): p. 36–42.

75. Lutz, V., M. Sattler, S. Gallinat, H. Wenck, R. Poertner, and F. Fischer, Impact of collagen crosslinking on the second harmonic generation signal and the fluorescence lifetime of collagen autofluorescence. Skin Res Technol, 2012. 18(2): p. 168–79.

76. Engler, A.J., S. Sen, H.L. Sweeney, and D.E. Discher, Matrix elasticity directs stem cell lineage specification. Cell, 2006. 126(4): p. 677–89.

77. Arnoczky, S.P., M. Lavagnino, M. Egerbacher, O. Caballero, and K. Gardner, Matrix metalloproteinase inhibitors prevent a decrease in the mechanical properties of stress-deprived tendons: an in vitro experimental study. Am J Sports Med, 2007. 35(5): p. 763–9.

78. Arnoczky, S.P., T. Tian, M. Lavagnino, and K. Gardner, Ex vivo static tensile loading inhibits MMP-1 expression in rat tail tendon cells through a cytoskeletally based mechanotransduction mechanism. J Orthop Res, 2004. 22(2): p. 328–33.

79. Sun, Y.L., A.R. Thoreson, S.S. Cha, C. Zhao, K.N. An, and P.C. Amadio, Temporal response of canine flexor tendon to limb suspension. J Appl Physiol (1985), 2010. 109(6): p. 1762–8.

80. Langberg, H., D. Skovgaard, L.J. Petersen, J. Bulow, and M. Kjaer, Type I collagen synthesis and degradation in peritendinous tissue after exercise determined by microdialysis in humans. J Physiol, 1999. 521 Pt 1: p. 299–306.

81. Dideriksen, K., A.K. Sindby, M. Krogsgaard, P. Schjerling, L. Holm, and H. Langberg, Effect of acute exercise on patella tendon protein synthesis and gene expression. Springerplus, 2013. 2(1): p. 109.

82. Wunderli, S.L., U. Blache, A. Beretta Piccoli, B. Niederost, C.N. Holenstein, F.S. Passini, U. Silvan, L. Bundgaard, U. Auf dem Keller, and J.G. Snedeker, Tendon response to matrix unloading is determined by the patho-physiological niche. Matrix Biol, 2020. 89: p. 11–26.

83. Lavagnino, M. and S.P. Arnoczky, In vitro alterations in cytoskeletal tensional homeostasis control gene expression in tendon cells. J Orthop Res, 2005. 23(5): p. 1211–8.

84. Gardner, K., M. Lavagnino, M. Egerbacher, and S.P. Arnoczky, Re-establishment of cytoskeletal tensional homeostasis in lax tendons occurs through an actin-mediated cellular contraction of the extracellular matrix. J Orthop Res, 2012. 30(11): p. 1695–701.

85. Lavagnino, M., A.E. Brooks, A.N. Oslapas, K.L. Gardner, and S.P. Arnoczky, Crimp length decreases in lax tendons due to cytoskeletal tension, but is restored with tensional homeostasis. J Orthop Res, 2017. 35(3): p. 573–579.

86. Lavagnino, M., S.P. Arnoczky, M. Egerbacher, K.L. Gardner, and M.E. Burns, Isolated fibrillar damage in tendons stimulates local collagenase mRNA expression and protein synthesis. J Biomech, 2006. 39(13): p. 2355–62.

87. Kjaer, M., Role of extracellular matrix in adaptation of tendon and skeletal muscle to mechanical loading. Physiol Rev, 2004. 84(2): p. 649–98.

88. Chang, J., R. Garva, A. Pickard, C.C. Yeung, V. Mallikarjun, J. Swift, D.F. Holmes, B. Calverley, Y. Lu, A. Adamson, H. Raymond-Hayling, O. Jensen, T. Shearer, Q.J. Meng, and K.E. Kadler, Circadian control of the secretory pathway maintains collagen homeostasis. Nat Cell Biol, 2020. 22(1): p. 74–86.

89. Boyera, N., I. Galey, and B.A. Bernard, Effect of vitamin C and its derivatives on collagen synthesis and cross-linking by normal human fibroblasts. Int J Cosmet Sci, 1998. 20(3): p. 151–8.

90. Hall, G., K.B. Tilbury, K.R. Campbell, K.W. Eliceiri, and P.J. Campagnola, Experimental and simulation study of the wavelength dependent second harmonic generation of collagen in scattering tissues. Opt Lett, 2014. 39(7): p. 1897–900.

91. Maller, O., A.P. Drain, A.S. Barrett, S. Borgquist, B. Ruffell, I. Zakharevich, T.T. Pham, T. Gruosso, H. Kuasne, J.N. Lakins, I. Acerbi, J.M. Barnes, T. Nemkov, A. Chauhan, J. Gruenberg, A. Nasir, O. Bjarnadottir, Z. Werb, P. Kabos, Y.Y. Chen, E.S. Hwang, M. Park, L.M. Coussens, A.C. Nelson, K.C. Hansen, and V.M. Weaver, Tumour-associated macrophages drive stromal cell-dependent collagen crosslinking and stiffening to promote breast cancer aggression. Nat Mater, 2021. 20(4): p. 548–559.

92. Diop-Frimpong, B., V.P. Chauhan, S. Krane, Y. Boucher, and R.K. Jain, Losartan inhibits collagen I synthesis and improves the distribution and efficacy of nanotherapeutics in tumors. Proc Natl Acad Sci U S A, 2011. 108(7): p. 2909–14.

93. Chauhan, V.P., I.X. Chen, R. Tong, M.R. Ng, J.D. Martin, K. Naxerova, M.W. Wu, P. Huang, Y. Boucher, D.S. Kohane, R. Langer, and R.K. Jain, Reprogramming the microenvironment with tumor-selective angiotensin blockers enhances cancer immunotherapy. Proc Natl Acad Sci U S A, 2019. 116(22): p. 10674–10680.

94. Pinter, M., A. Weinmann, M.A. Worns, F. Hucke, S. Bota, J.U. Marquardt, D.G. Duda, R.K. Jain, P.R. Galle, M. Trauner, M. Peck-Radosavljevic, and W. Sieghart, Use of inhibitors of the renin-angiotensin system is associated with longer survival in patients with hepatocellular carcinoma. United European Gastroenterol J, 2017. 5(7): p. 987–996.

95. Kutys, M.L. and K.M. Yamada, An extracellular-matrix-specific GEF-GAP interaction regulates Rho GTPase crosstalk for 3D collagen migration. Nat Cell Biol, 2014. 16(9): p. 909–17.

96. Dingal, P.C., A.M. Bradshaw, S. Cho, M. Raab, A. Buxboim, J. Swift, and D.E. Discher, Fractal heterogeneity in minimal matrix models of scars modulates stiff-niche stem-cell responses via nuclear exit of a mechanorepressor. Nat Mater, 2015. 14(9): p. 951–60.

97. Suzuki, K., J.J. Enghild, T. Morodomi, G. Salvesen, and H. Nagase, Mechanisms of activation of tissue procollagenase by matrix metalloproteinase 3 (stromelysin). Biochemistry, 1990. 29(44): p. 10261–70.

98. Kasznel, A.J., *Design, Synthesis, Structural Studies, & Applications of Synthetic Collagen Peptides*. 2020, University of Pennsylvania.

99. Biela, E., J. Galas, B. Lee, G.L. Johnson, Z. Darzynkiewicz, and J.W. Dobrucki, Col-F, a fluorescent probe for ex vivo confocal imaging of collagen and elastin in animal tissues. Cytometry A, 2013. 83(6): p. 533–9.

100. Lacomb, R., O. Nadiarnykh, S.S. Townsend, and P.J. Campagnola, Phase Matching considerations in Second Harmonic Generation from tissues: Effects on emission directionality, conversion efficiency and observed morphology. Opt Commun, 2008. 281(7): p. 1823–1832.

101. Williams, R.M., J.B. Shear, W.R. Zipfel, S. Maiti, and W.W. Webb, Mucosal mast cell secretion processes imaged using three-photon microscopy of 5-hydroxytryptamine autofluorescence. Biophys J, 1999. 76(4): p. 1835–46.

102. Jayyosi, C., G. Fargier, M. Coret, and K. Bruyere-Garnier, Photobleaching as a tool to measure the local strain field in fibrous membranes of connective tissues. Acta Biomater, 2014. 10(6): p. 2591–601.

103. Lee, J.C., D.T. Wong, and D.E. Discher, Direct measures of large, anisotropic strains in deformation of the erythrocyte cytoskeleton. Biophys J, 1999. 77(2): p. 853–64.

104. Rezakhaniha, R., A. Agianniotis, J.T. Schrauwen, A. Griffa, D. Sage, C.V. Bouten, F.N. van de Vosse, M. Unser, and N. Stergiopulos, Experimental investigation of collagen waviness and orientation in the arterial adventitia using confocal laser scanning microscopy. Biomech Model Mechanobiol, 2012. 11(3-4): p. 461–73.

105. Avila, F.J. and J.M. Bueno, Analysis and quantification of collagen organization with the structure tensor in second harmonic microscopy images of ocular tissues. Appl Opt, 2015. 54(33): p. 9848–54.

106. Phair, R.D., S.A. Gorski, and T. Misteli, Measurement of dynamic protein binding to chromatin in vivo, using photobleaching microscopy. Methods Enzymol, 2004. 375: p. 393–414.

107. Smith, M.S., W.M. Billings, F.G. Whitby, M.B. Miller, and J.L. Price, Enhancing a long-range salt bridge with intermediate aromatic and nonpolar amino acids. Org Biomol Chem, 2017. 15(28): p. 5882–5886.

108. Stahl, P.J., J.C. Cruz, Y. Li, S. Michael Yu, and K. Hristova, On-the-resin N-terminal modification of long synthetic peptides. Anal Biochem, 2012. 424(2): p. 137–9.

109. Cai, X., Z. Liu, S. Zhao, C. Song, S. Dong, and J. Xiao, A single stranded fluorescent peptide probe for targeting collagen in connective tissues. Chem Commun (Camb), 2017. 53(87): p. 11905–11908.

110. Hyun, S., L. Li, K.C. Yoon, and J. Yu, An amphipathic cell penetrating peptide aids cell penetration of cyclosporin A and increases its therapeutic effect in an in vivo mouse model for dry eye disease. Chem Commun (Camb), 2019. 55(91): p. 13657–13660.

111. Bonkowski, B., J. Wieczorek, M. Patel, C. Craig, A. Gravelin, and T. Boncher, Basic Concepts of using Solid Phase Synthesis to Build Small Organic Molecules using 2-Chlorotrityl Chloride Resin. Modern Chemistry & Applications, 2013. 1(4): p. 113.

